# Iron acquisition by a commensal bacterium modifies host nutritional immunity during *Salmonella* infection

**DOI:** 10.1101/2023.06.25.546471

**Authors:** Luisella Spiga, Ryan T. Fansler, Yasiru R. Perera, Nicolas G. Shealy, Matthew J. Munneke, Teresa P. Torres, Holly E. David, Andrew Lemoff, Xinchun Ran, Katrina L. Richardson, Nicholas Pudlo, Eric C. Martens, Zhongyue J. Yang, Eric P. Skaar, Mariana X. Byndloss, Walter J. Chazin, Wenhan Zhu

## Abstract

During intestinal inflammation, host nutritional immunity starves microbes of essential micronutrients such as iron. Pathogens scavenge iron using siderophores, which is counteracted by the host using lipocalin-2, a protein that sequesters iron-laden siderophores, including enterobactin. Although the host and pathogens compete for iron in the presence of gut commensal bacteria, the roles of commensals in nutritional immunity involving iron remain unexplored. Here, we report that the gut commensal *Bacteroides thetaiotaomicron* acquires iron in the inflamed gut by utilizing siderophores produced by other bacteria including *Salmonella,* via a secreted siderophore-binding lipoprotein termed XusB. Notably, XusB-bound siderophores are less accessible to host sequestration by lipocalin-2 but can be “re-acquired” by *Salmonella*, allowing the pathogen to evade nutritional immunity. As the host and pathogen have been the focus of studies of nutritional immunity, this work adds commensal iron metabolism as a previously unrecognized mechanism modulating the interactions between pathogen and host nutritional immunity.

## Introduction

The human large intestine is home to a vast number of microorganisms comprising fungi, archaea, viruses, and bacteria^1, 2^. Similar to other ecosystems, the gut microbial community is subjected to constant perturbations including changes in diet, use of antibiotics, and intestinal inflammation^3^. Such disturbances cause transient changes in the composition and function, or even reorganization of the gut ecosystem to an alternative equilibrium^4^. In the face of disturbance, the intestinal microbial community can persist in equilibrium, or return to the original state as the perturbation passes, which together describes the ecosystem’s resilience^5^. When perturbations exceed the resilience capacity, the microbiome can be driven into imbalanced states, with the hallmark feature being a permanently decreased species diversity^6^. Altered composition and function of the gut microbiota have been associated with the exacerbation of a wide range of infectious and non-communicable diseases ^6–8^. Despite being a pillar in maintaining the structure and function of the human gut microbiota over time^3^, our understanding of microbiota resilience remains limited.

As the nutrient landscape shapes the structure and function of the inhabiting microbial ecosystem^9^, changes in nutrient availabilities constitute one of the most critical perturbations to the gut microbiome^10^. One of the essential nutrients is iron, which nearly all organisms require as a cofactor in metalloproteins that carry out essential cellular functions (e.g. energy generation and DNA replication)^11^. Vertebrate hosts exploit the microbial dependency on iron and other nutrient metals to exert control over microbial growth, in a process termed nutritional immunity^12^. During episodes of gut inflammation, mucosal epithelial cells and immune cells limit bacterial growth by producing iron-scavenging proteins such as lactoferrin^13^. Pathogens, including *Salmonella enterica* serovar Typhimurium (*S*. Tm), compete for iron with the host by producing, small, high affinity iron chelators termed siderophores, including enterobactin^14^. This microbial iron capture strategy is counteracted by the host protein lipocalin-2, which sequesters iron laden enterobactin^15^. *S*. Tm, in turn, can produce a “stealth siderophore” salmochelin^16–18^, a glycosylated derivative of enterobactin not recognized by lipocalin-2^19^, to evade nutritional immunity defense and thrive in the inflamed gut^20, 21^.

Decades of research have characterized how siderophores allow pathogens such as *S*. Tm to survive and thrive in the inflamed gut^11, 20–23^. While the host and pathogen have long been considered the primary essential facets of nutritional immunity, it is important to note that the host-pathogen competition for iron occurs in the presence of gut commensal bacteria that are substantially more abundant than the pathogens^24–27^. In contrast to well-studied bacterial pathogens, how commensals cope with iron restriction during intestinal inflammation remains largely unexplored. Our previous work demonstrated that *B. thetaiotaomicron*, a representative of human gut commensal bacteria, sustains resilience in the inflamed gut by acquiring siderophores produced by other microbes (xenosiderophores), including *S*. Tm^28^. However, the molecular basis of commensal siderophore acquisition remains elusive. More importantly, whether commensal bacteria assume bystander roles or actively modify iron competition between host nutritional immunity and pathogens remains undefined.

Here, we show that *B. thetaiotaomicron* secretes a lipoprotein onto the cell surface and into the extracellular milieu to acquire xenosiderophores. Siderophores bound to this protein are shielded from lipocalin-2-mediated sequestration but are readily utilized by *S*. Tm. By recapturing siderophores from commensal bacteria, *S*. Tm can recover a pool of siderophores that are otherwise sequestered by host proteins. Our findings implicate commensal iron acquisition as a previously unrecognized facet of nutritional immunity.

## RESULTS

### *B. thetaiotaomicron* employs a specialized mechanism for xenosiderophore acquisition

In *B. thetaiotaomicron*, the <u>X</u>enosiderophore <u>U</u>tilization <u>S</u>ystem (Xus) operon encodes three genes (XusABC) that are indispensable for the utilization of xenosiderophore^28^ (**Fig. 1A**). While XusA contributes to *B. thetaiotaomicron* resilience in animal models of infectious and non-infectious colitis^28^, the roles of the additional two genes in this operon to *B. thetaiotaomicron* resilience *in vivo* remain undefined. Because secreted bacterial proteins shape interbacterial and host-microbe interactions by modulating nutrient competition (^29–33^, reviewed in^34^), we sought to determine whether *B. thetaiotaomicron* acquires siderophores using surface-exposed components.

**Figure 1:**
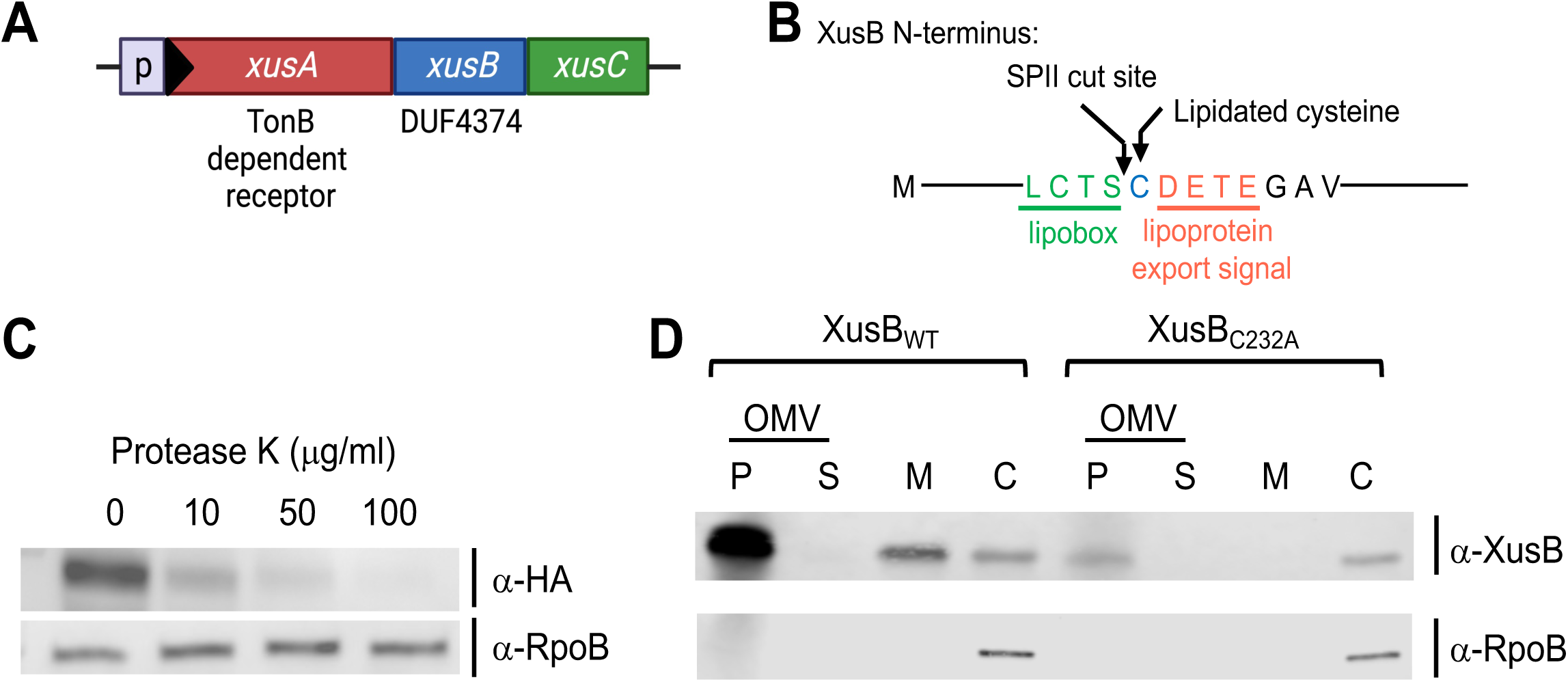
XusB is a surface-exposed protein that is enriched in outer membrane vesicles. (**A**) Schematic representation of the *xusABC* operon in *B. thetaiotaomicron* VPI-5482. (**B**) Sequence features of XusB N-terminus. (**C**) *B. thetaiotaomicron* cells were cultured in iron-limiting media and treated with indicated concentrations of protease K. Degradation of HA-tagged XusB on whole *B. thetaiotaomicron* cells was assessed via western blots. RpoB is a cytoplasmic control. (**D**) *B. thetaiotaomicron* cells were cultured in iron-limiting media and fractionated into outer membrane vesicles (OMV-P), cell-free supernatant (OMV-S), membrane (M), and cytoplasm/periplasm (C) fractions. Indicated fractions were resolved on SDS-PAGE gel and probed for XusB and RpoB. RpoB is a cytoplasmic control.

Sequence analysis predicts that XusA encodes multi-stranded antiparallel β-barrels that are typical of outer membrane-embedded TonB-dependent receptors^35^, while XusC harbors a conserved PepSY domain, a feature of a reductase family found on the bacterial inner membrane^36^. In contrast, XusB encodes a lipobox-like sequence at its N-terminus, a motif typical of bacterial lipidated proteins^37^. The lipobox-like sequence is followed by a putative conserved cysteine (Cys-23) and a lipoprotein export signal (LES), features that allow *Bacteroidetes* lipoprotein to be flipped on the outer leaflet of the outer membrane (**Fig. 1B**)^38^. Based on these features, we predicted that XusB is a surface-exposed lipoprotein. To test this hypothesis, we treated intact *B. thetaiotaomicron* cells with increasing concentrations of protease K, which degrades surface-exposed proteins. We then resolved the whole cell lysate on an SDS-PAGE gel. While the intracellular protein RpoB (RNA polymerase β-subunit) was protected from proteolysis, XusB was readily degraded by extracellular protease K in a dose-dependent manner, consistent with the notion that XusB is a surface-exposed protein (**Fig. 1C**). Moreover, cellular fractionation experiments revealed that XusB is associated with the cellular membrane (**Fig. 1D**). Substituting the conserved (Cys-23) for alanine abolished XusB membrane association, consistent with the notion that this residue is important for correct cellular localization (**Fig. 1B and 1D**). Mutating all four LES residues (Asp24, Glu25, Thr26, and Glu27; XusB_lesMT_) to alanine diminished the XusB levels in all fractions despite comparable transcription levels to the wild-type counterpart, indicating that these residues are important to the stability of XusB (**Supplementary Fig. 1A**). Of note, XusB is enriched in the bacterial culture supernatant (**Fig. 1D**, first lane). Further fractionation of cell-free supernatant via ultracentrifugation revealed that XusB was present in outer membrane vesicles (OMVs), small, spherically bilayered vesicles derived from the outer membrane of diverse Gram negative bacteria^39^. This is in accordance with previous reports demonstrating that XusB was differentially enriched in the OMV fraction in both bacterial cultures and in the murine gastrointestinal tract^40, 41^. Consistent with the notion that localization to the cell exterior is required for incorporation into OMVs^40^, mutating Cys23 or LES residues to alanine expectedly impaired the recruitment of XusB into OMVs (**Fig. 1D** and **Supplementary Fig. 1A**). Based on these observations, we hypothesize that *B. thetaiotaomicron* secretes XusB into the extracellular milieu to aid in the capture and import iron in the form of xenosiderophores.

### XusB directly binds to enterobactin with high affinity

Previous work revealed that catecholate siderophores, including enterobactin and salmochelin, sustain *B. thetaiotaomicron* resilience and are ligands for the Xus system^28, 42^. To explore the role of XusB in extracellular siderophore capture, we tested whether recombinant XusB binds to enterobactin. Recombinant XusB has limited solubility in aqueous buffers, so we generated a XusBΔN construct lacking the 40 N-terminal residues encoding the lipobox and LES, which are not likely involved in ligand binding. As anticipated, XusBΔN exhibited greatly improved solubility, which enabled isolation of highly pure recombinant protein and subsequent biophysical and structural analyses. When XusBΔN and iron-laden enterobactin (Fe-Ent) were co-incubated at a 1:1 ratio, we were able to isolate the complex on size exclusion chromatography (**Fig. 2A**); enterobactin (0.66 kDa) co-eluted in the 50 kDa XusB (280 nm) peak as reflected in the observation of UV signal in the absorbance at 550 nm^43^ that is characteristic for Fe-Ent. XusB is expected to bind Fe-enterobactin with high affinity to compete with the invading pathogen. To test this hypothesis, we measured the affinity by monitoring the intrinsic tryptophan fluorescence of XusB as it is titrated with Fe-Ent^44^. We observed a concentration-dependent quenching of XusB intrinsic fluorescence as Fe-Ent was added, indicating the microenvironment of one or more tryptophan residues changes as enterobactin is bound (**Fig. 2B**). A dissociation constant of 148 nM was extracted from the data. These results support a model in which *B. thetaiotaomicron* utilizes XusB to capture the xenosiderophore Fe-Ent.

**Figure 2:**
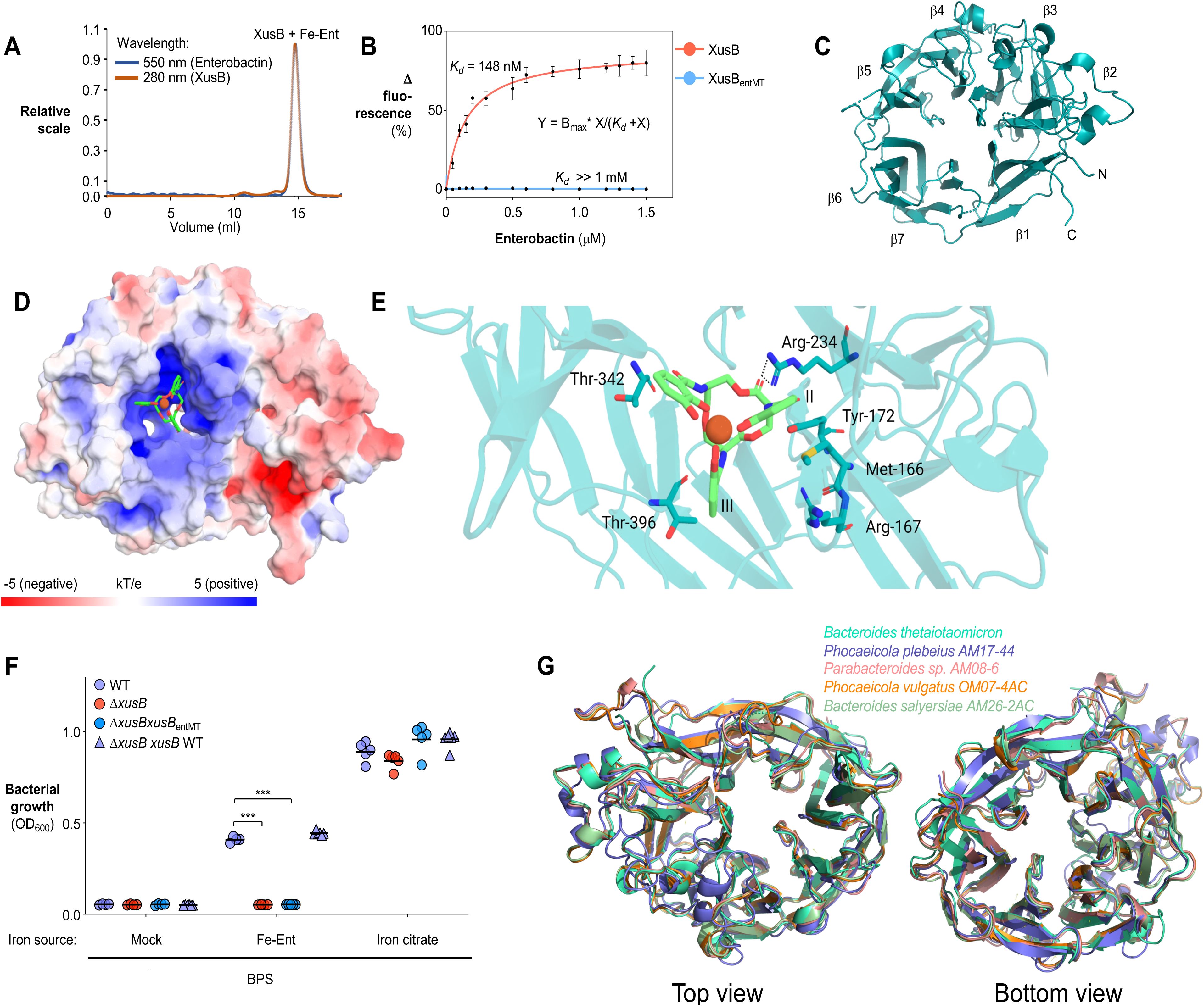
*Bacteroidetes* encode homologs of XusB, which binds to enterobactin with high affinity. (**A**) Size exclusion chromatography traces for recombinant XusB (280 nm) incubated with Fe-enterobactin (550 nm). (**B**) Changes of intrinsic tryptophan fluorescence (Δ fluorescence) in XusB upon binding to enterobactin. The dissociation constant was derived by fitting Ent concentrations vs. fluorescence intensity changes to a one-site specific binding model. (**C**) The overall structure of XusBΔN. N- and C-termini are indicated, and blades are numbered. Each β-blade consists of four β-strands (ABCD). (**D**) Overall electrostatic profile of XusB docked with Fe-Ent. Enterobactin is shown as sticks with carbon atoms colored in green, nitrogen in blue, and oxygen in red, with the Fe^3+^ represented as an orange sphere. The surface electrostatic field was calculated using Adaptive Poisson-Boltzmann Solver (ABPS). The color scale represents electrostatic potentials in units of kT/e ranging from −5 (red, negatively charged) to +5 (blue, positively charged). (**E**) Close-up of XusBΔN residues interacting with Fe^3+^-Ent. Residues and catechol arms of Fe^3+^-Ent are numbered. (**F**) *B. thetaiotaomicron* strains were cultured in BHIS medium, followed by subculturing in iron-limited (200 μmol/L of BPS) SDM medium supplemented with 0.5 μM enterobactin. Growth of *B. thetaiotaomicron* was determined by measuring the optical density (OD_600_). (**G**) Overlay of ΔN-terminus structures of XusB and its homologs. Bars represent the geometric mean. ***, *P* < 0.001.

### Fe-enterobactin binds to the central channel of the XusB β-propeller

To understand the molecular basis for enterobactin binding by XusB, we set out to determine the three-dimensional structure of XusB in the absence and presence of the siderophore, using the XusBΔN construct. Crystallization trials on the isolated protein produced crystals that diffracted to 2.5 Å. Although XusB does not have substantial homology with any protein whose structure is available in the Protein Data Bank, the phase problem could be solved by molecular replacement using a model generated by AlphaFold2^45^. XusBΔN forms a seven-bladed β-propeller arranged around a pseudo seven-fold axis with a deep and wide pocket in the middle of the propeller and an axial channel running through the center of the propeller (**Fig. 2C, Supplementary Table 1**), features that are unique among bacterial enterobactin receptors. The edges of the β- strands, together with Thr296 and Tyr445, delineate a ligand-accessible, polar calyx in the center of the propeller. Consistent with the notion that seven-bladed β-propeller proteins can use such pockets to bind and transport ligands^46^, the central channel is highly positively charged (**Fig. 2D**), well suited to bind the negatively charged Fe enterobactin.

To understand the details of Fe-enterobactin binding to XusB, we set out to determine the crystal structure of the complex. However, extensive crystallization trials attempting both soaking the ligand into XusBΔN crystals or co-crystallization of the protein and ligand yielded no diffraction-quality crystals, possibly due to the partial degradation of Fe-Ent in the process of crystallization, as reported previously^15^. Since a crystallographic approach was not feasible, we sought to obtain a structural model using a computational docking approach starting from the structure of the free protein.

Confidence in a computational model is dependent on having an accurate starting structure, knowledge of where the ligand binds to the protein, and experimental validation. We were confident that Fe-Ent binds in the XusB calyx. Small angle X-ray scattering (SAXS) was used to determine if the structure of XusB is perturbed upon binding of Fe-Ent. Back-calculations of the SAXS profile from the crystal structure fit very well with the experimental data for the free protein (*x^2^* = 1.33) (**Supplementary Fig. 1B**). Moreover, comparisons of the SAXS data for the protein in the absence and presence of Fe enterobactin show very little change in XusBΔN upon binding (**Supplementary Fig. 1C**). These data indicate that the crystal structure serves as an excellent starting structure for the docking of Fe-Ent to XusB.

Enterobactin has a sufficient degree of structural flexibility that docking calculations with two different Fe-Ent starting conformations were performed to ensure the conclusions drawn were accurate. The two Fe-enterobactin structures were then each docked to the structure of XusBΔN using the AutoDock server^47^. The docked structures were similar, revealing that Ent bound deep within the highly basic concave surface of XusB, contacting residues from “A” strand in β3, β4, β5, and β6, including Arg234, which makes multiple hydrogen bonds to Fe^3+^-Ent (**Fig. 2D&E**). The catechol arms I and II of the Fe^3+^-Ent sit between β4, β5, and β6 making hydrophobic interactions, whereas arm III makes minimal interactions and is partially exposed to the solvent (**Fig. 2E**).

To validate the structural model, six XusB residues contacting Ent were mutated to alanine (XusB_entMT_). These mutations completely abolished enterobactin binding despite displaying similar protein levels and cellular localization as the wild-type counterpart (**Fig. 2B** and **Supplementary Fig. 1D**). Moreover, a *B. thetaiotaomicron* strain expressing the enterobactin-binding deficient mutant (XusB_entMT_) failed to utilize enterobactin as the sole iron source (**Fig. 2F**), further highlighting the importance of the highly positively charged concave surface of XusBΔN for binding the negatively charged enterobactin.

### *Bacteroidetes* members encode structural homologs of XusB

Structural analysis enabled us to interrogate whether XusB represents an example of a broader nutrient acquisition strategy utilized by diverse gut commensal bacteria. To investigate the distribution of XusB homologs in the human microbiome, we performed a Blastp search in 1,500 human gut metagenome-assembled genomes^48^. While multiple *Bacteroidetes* species encode XusB homologs, with conserved genetic synteny similar to the Xus operon, the primary sequences are poorly preserved (**Supplementary Fig. 2A-B**). We thus explored whether these proteins are structurally conserved using AlphaFold2, which has been shown to accurately predict tertiary structure of single-domain globular proteins such as XusB^45^. Indeed, as noted above, the AlphaFold2 predicted XusBΔN structure was sufficiently accurate to be used as a starting model for experimental determination of the X-ray crystal structure, with a pair-wise (predicted vs. experimental structure) root mean square deviation (RMSD) value of 0.553 Å. We thus used AlphaFold2 to predict the tertiary structures of XusB homologs. We found all predicted structures of XusB homologs share a flexible region at N-terminus and adopt a characteristic seven-bladed β-propeller fold, a feature of the conserved <u>d</u>omain of <u>u</u>nknown function 4374 (DUF4374) that is exclusively found in the *Bacteroidetes* phylum. Of note, the models of these homologs have high overall similarity to XusB (RMSD ∼1 Å, **Fig. 2G**, **Supplementary Fig. 3A**), but are structurally distinct from other *Bacteroidetes* nutrient binding lipoproteins, as well as siderophore binding proteins in Gram-positive bacteria (rmsd>>10 Å, **Supplementary Fig. 3A**). Additionally, all homologs exhibit a positively charged central calyx, similar to XusB (**Supplementary Fig. 3B**), suggesting that these proteins may bind to negatively charged ligands such as enterobactin. Together, these data suggest that XusB and its homologs represent an example of a conserved nutrient acquisition strategy among members of *Bacteroidetes*.

### XusB can function as an early step in extracellular siderophore capture

Because XusB binds to enterobactin with high affinity, is indispensable in xenosiderophore utilization, and is secreted into the extracellular milieu, we sought to explore if XusB functions extracellularly as an early step for siderophore acquisition. To this end, we first cultured indicated *B. thetaiotaomicron* strains in a modified semi-defined medium^49^, followed by the addition of an extracellular iron chelating agent bathophenanthroline disulfonate (BPS) to model iron-limiting conditions. We collected the cell-free fraction of *B. thetaiotaomicron* cultures and separated the OMV and the supernatant fractions. We then loaded these fractions with Fe-Ent, removed unbound siderophores by repeat ultrafiltration, fed them to recipient cells, and measured bacterial growth (**Fig. 3A**).

**Figure 3:**
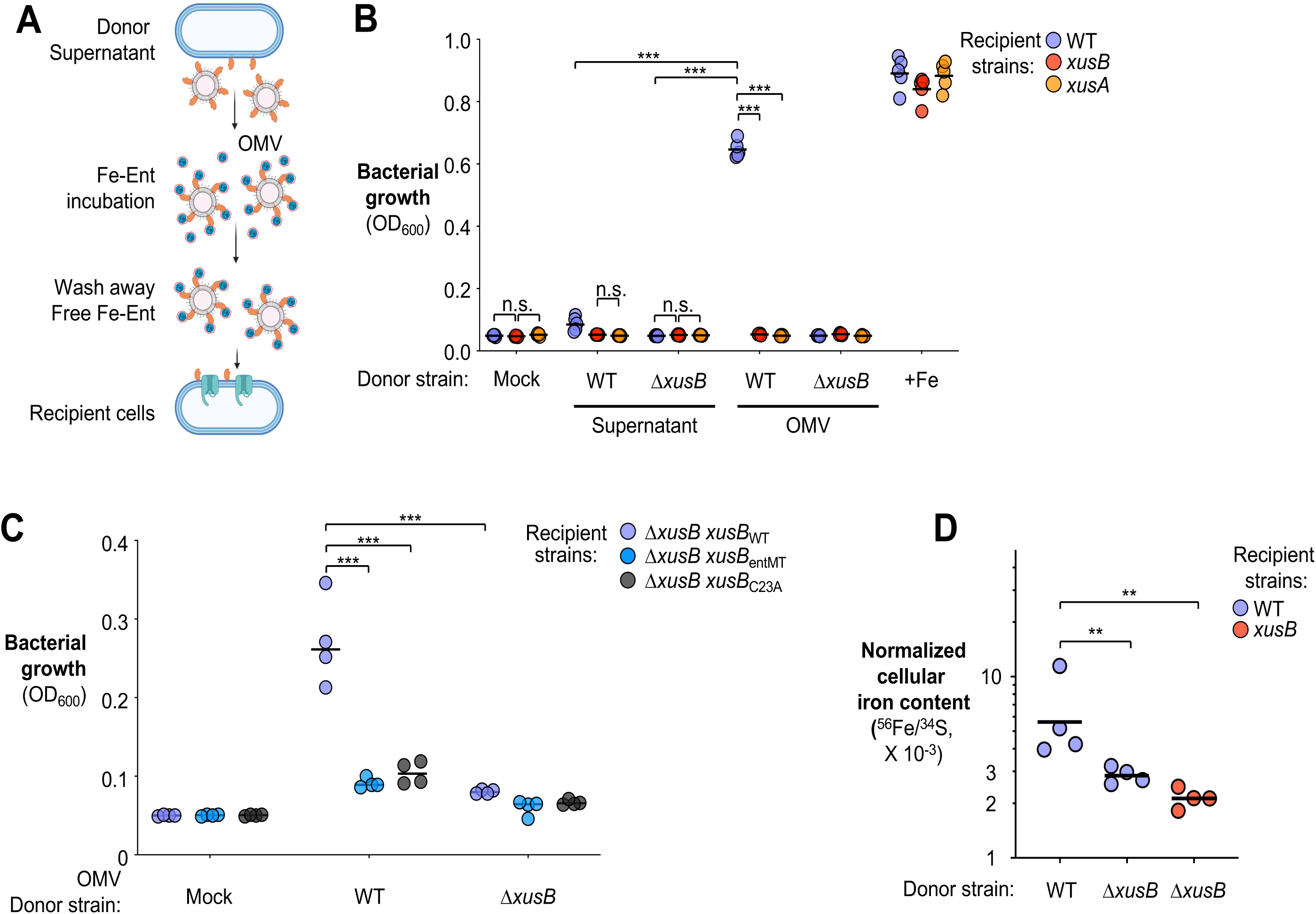
XusB can function as an early step in xenosiderophore acquisition. (**A-C**) Supernatant from donor *B. thetaiotaomicron* strains grown in iron-limited medium (BPS-supplemented SDM medium) was filter-sterilized, loaded with Fe-Ent, ultracentrifuged to collect OMV fraction, washed to remove unbound ligand, and introduced to recipient cells in BPS-supplemented SDM medium. (**A**) Schematic representation and (**B-C**) Bacterial growth measured by OD_600_. Bars represent the geometric mean. (**D**) The *B. thetaiotaomicron* wild-type strain and an isogenic *xusB* mutant were cultured in iron-deprived media to exhaust endogenous iron before subcultured in the presence of Fe-Ent loaded OMVs collected from indicated strains. ICP-MS was used to assess cellular iron levels. Bars represent the geometric mean. **, *P* < 0.01; ***, *P* < 0.001; n.s., not statistically significant.

Under iron limiting conditions (BPS supplemented semi-defined media), the Fe-Ent loaded OMV fraction derived from the wild-type *B. thetaiotaomicron* strain supported robust growth of the wild-type recipient cells, but not cells lacking XusB or the enterobactin transporter XusA (**Fig. 3B**). By contrast, the remaining supernatant after depleting OMVs by ultracentrifugation did not promote growth of the same recipient cells (**Fig. 3B**). The OMV fraction collected from XusB-deficient donors did not promote the growth of wild type recipient cells (**Fig. 3B**), suggesting that the growth of the recipient cells was not due to contaminating free Fe-Ent associated with the OMVs. Further, Fe-Ent feeding required the donor cells expressing the wild-type, but not a missorted mutant, nor an enterobactin binding-deficient mutant of XusB (**Fig. 3C**). Conversely, Fe-Ent loaded XusB also failed to rescue the growth of recipient cells expressing the enterobactin-binding deficient mutant (**Supplementary Fig. 4A**). Consistent with the role of XusB in capture and delivery of iron in enterobactin, supplementation of Fe-Ent-laden OMVs derived from XusB^+^ *B. thetaiotaomicron*, but not the *xusB*-deficient mutant, significantly increased the cellular iron levels in wild-type cells (**Fig. 3D**). These data suggest that OMV-associated XusB can function in the extracellular milieu to capture enterobactin to promote *B. thetaiotaomicron* growth during iron starvation, and this process requires the presence of functional XusB on the surface the recipient cells.

### XusB sustains *B. thetaiotaomicron* resilience during *S*. Tm infection

XusB assumes an indispensable role in *B. thetaiotaomicron* xenosiderophores utilization *in vitro* by serving as the initial step in extracellular siderophore capture (**Fig. 3**). Given the crucial role of xenosiderophore utilization in sustaining commensal fitness in the inflamed gut^28^, we hypothesize that XusB contributes to *B. thetaiotaomicron* resilience during intestinal inflammation. To test this hypothesis, we explored the fitness advantage conferred by XusB in mice challenged by *S*. Tm, a pro-inflammatory pathogen that induces robust iron limitation in the gastrointestinal tract and produces enterobactin, which is accessible to *B. thetaiotaomicron* in a XusB-dependent manner (Ref^20^ and **Fig. 2F**). As conventionally raised mice are resistant to *B. thetaiotaomicron* colonization^50^, we treated C57BL/6 mice with an antibiotic cocktail to promote *B. thetaiotaomicron* engraftment^51^. We then inoculated these mice with an equal mixture of the *B. thetaiotaomicron* wild-type strain and an isogenic mutant lacking *xusB* (Δ*xusB*), or an isogenic strain that expresses the enterobactin binding deficient XusB mutant (Δ*xusB xusB*_entMT_). Two days post-colonization, groups of mice were either mock treated with LB broth or intragastrically challenged with the *S*. Tm wild-type strain or a mutant (*entB*) deficient in producing both enterobactin and salmochelin (**Fig. 4A**). Four days post *S*. Tm challenge, we collected the intestinal contents and quantified the abundance of *B. thetaiotaomicron* and *S*. Tm strains. As anticipated, the Δ*xusB* mutant displayed fitness levels similar to that of the wild-type strain in mock-treated animals, but was outcompeted in *S*. Tm challenged mice (**Fig. 4B**). The *xusB*entMT was similarly outcompeted by its wild type counterpart during *S*. Tm infection, highlighting the role of enterobactin binding in XusB as a resilience factor (**Fig. 4B**). Notably, the fitness advantage conferred by the XusB was diminished in mice challenged with an S. Tm *entB* mutant^52^, consistent with the notion that the experimentally introduced *S*. Tm is a major source of catecholate siderophores. (**Fig. 4B**).

**Figure 4:**
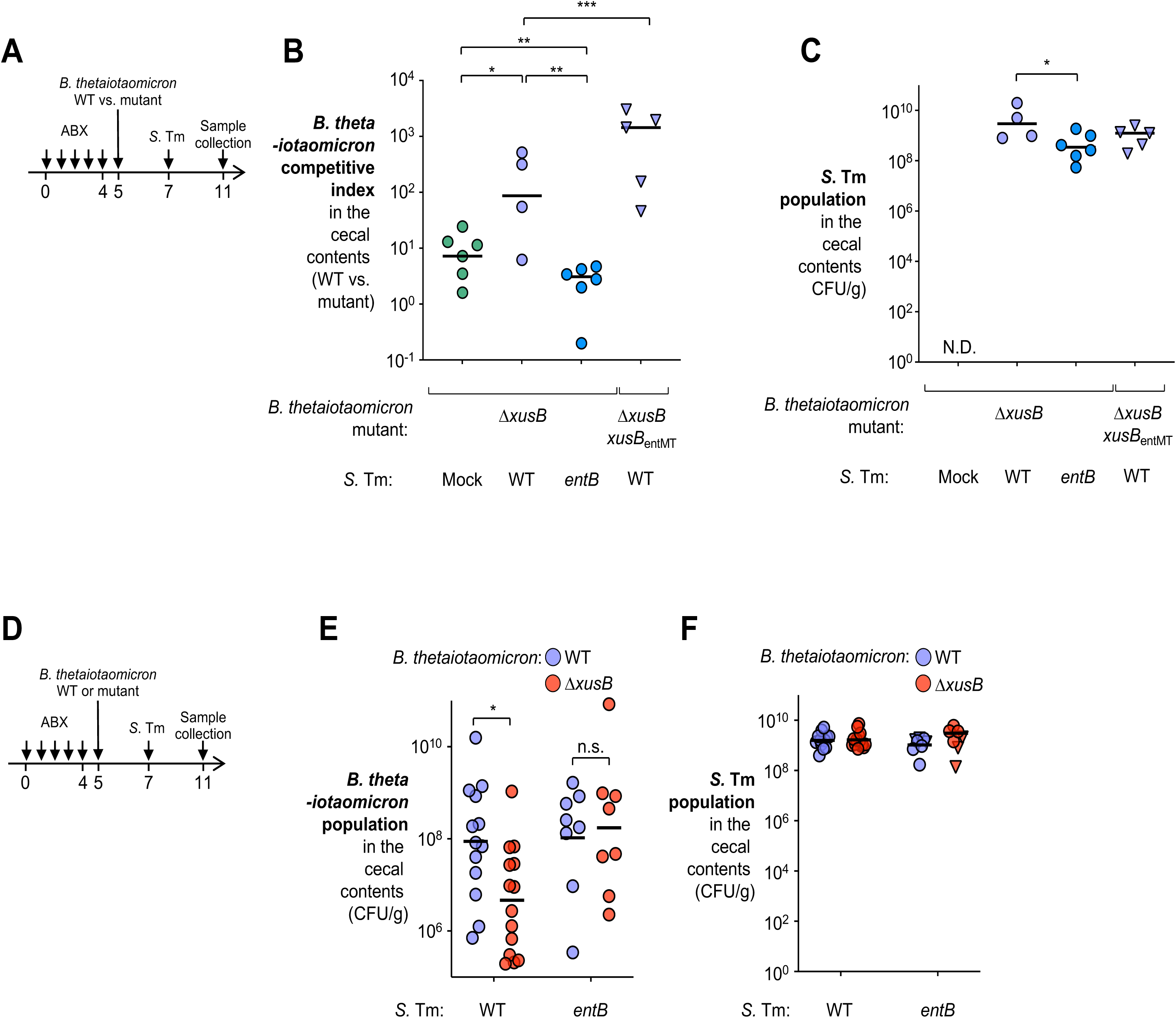
Role of *xusB* in *B. thetaiotaomicron* resilience during murine *S*. Tm infection. (**A-C**) Groups of C57BL/6 mice were treated with a cocktail of antibiotics, followed by intragastrical inoculation of an equal mixture of the *B. thetaiotaomicron* wild-type strain and an isogenic mutant (Δ*xusB*), or a strain expressing the enterobactin-binding deficient mutant (Δ*xusB xusB*_entMT_). Mice were then either mock-treated (*N* = 6), intragastrically challenged with *S*. Tm SL1344 (WT vs. Δ*xusB*, *N* = 4; WT vs. Δ*xusB xusB*entMT, *N* = 5), or an isogenic *entB* mutant (WT vs. Δ*xusB*, *N* = 6). (**A**) Schematic presentation of the experiment. Four days after infection, the cecal contents were collected, and the abundance of (**B**) *B. thetaiotaomicron* and (**C**) *S*. Tm was determined by plating on selective agar. The competitive index was calculated as the ratio of the two strains in the cecal content, corrected by the ratio in the inoculum. (**D-F**) Groups of C57BL/6 mice were treated with a cocktail of antibiotics, followed by intragastrical inoculation of either the *B. thetaiotaomicron* wild-type strain or an isogenic Δ*xusB* mutant. Mice were challenged with the *S*. Tm SL1344 wild-type strain (*B. thetaiotaomicron* wild-type*, N* = 13; *B. thetaiotaomicron* Δ*xusB, N =* 14) or an isogenic *entB* mutant (*B. thetaiotaomicron* wild type*, N* = 8; *B. thetaiotaomicron* Δ*xusB, N =* 7) for 4 days. The abundance of each *B. thetaiotaomicron* (**E**) and *S*. Tm (**F**) strain in cecal contents was determined by plating on selective agar. Bars represent the geometric mean. *, *P* < 0.05; **, *P* < 0.01; ***; *P* < 0.001; n.s., not statistically significant.

Next, we sought to more directly determine whether XusB sustains *B. thetaiotaomicron* resilience in competition with members of the microbiota during colitis. To explore this, we associated antibiotic-pretreated mice with the *B. thetaiotaomicron* wild-type strain or a Δ*xusB* mutant. Two days later, mice were challenged with the *S*. Tm wild-type strain for four days. Mice challenged with an *S*. Tm *entB* mutant were used as controls (**Fig. 4D**). Quantification of the *B. thetaiotaomicron* intestinal population revealed that the Δ*xusB* mutant displayed a significant fitness defect in mice challenged with the *S*. Tm wild-type strain (**Fig. 4E**). The fitness differences between *B. thetaiotaomicron* strains were diminished in mice infected by the *S*. Tm *entB* mutant (**Fig. 4E**). In both competitive and single colonization settings, while the *S*. Tm strain displayed slightly higher fitness than the *entB* mutant strain, no significant differences in markers of mucosal inflammation, except *Nos2* (*Nitric oxide synthase 2*), were noted (**Fig. 4C, F** and **Supplementary Fig. 5**). Together, these data suggest that XusB participates in xenosiderophore acquisition and sustaining *B. thetaiotaomicron* resilience in the context of infectious colitis.

### XusB mediates inter-species competition for siderophore *in vitro*

Secreted factors mediate interbacterial nutritional competition and shape the composition of the microbial community (^22, 53, 54^, reviewed in^55, 56^). As XusB-containing OMVs capture xenosiderophores, and the delivery of XusB-bound enterobactin requires XusB produced by the recipient cells, we predict that strain-specific factors might be required for the retrieval of XusB-bound enterobactin, and that XusB could sequester enterobactin from other competing species. To explore this possibility, we cultured a *Bacteroides* strain, *B*. *vulgatus,* which expresses XusB homolog, in iron-limiting media supplemented with free Fe-Ent or XusB-bound Fe-Ent. While free Fe-Ent supported robust growth of *B*. *vulgatus*^28^, XusB-bound Fe-Ent failed to rescue the same strain (**Fig. 5A-B**). Notably, XusB was not degraded when incubated with *B. thetaiotaomicron* or *B. vulgatus* (**Supplementary Fig. 6A**). Because other members of the *Bacteroidetes* phylum encode XusB homologs (e.g., *B. vulgatus* and *B. salyersiae*, **Fig. 2G** & **Supplementary Fig. 2**), we sought to explore whether species-specific XusB homologs capture siderophores in a “selfish” manner. Consistent with this notion, the Fe-Ent loaded OMV fraction of *B. vulgatus* could supply Fe-Ent as the sole iron source when *B. vulgatus* was the recipient strain. In contrast, the same OMV fraction blocked *B. salyersiae*, which can readily utilize free Fe-Ent as the sole iron source, from utilizing the xenosiderophore (**Supplementary Fig. 6B**). Together, these results indicate that XusB and its homolog may mediate interbacterial competition for siderophores.

**Figure 5:**
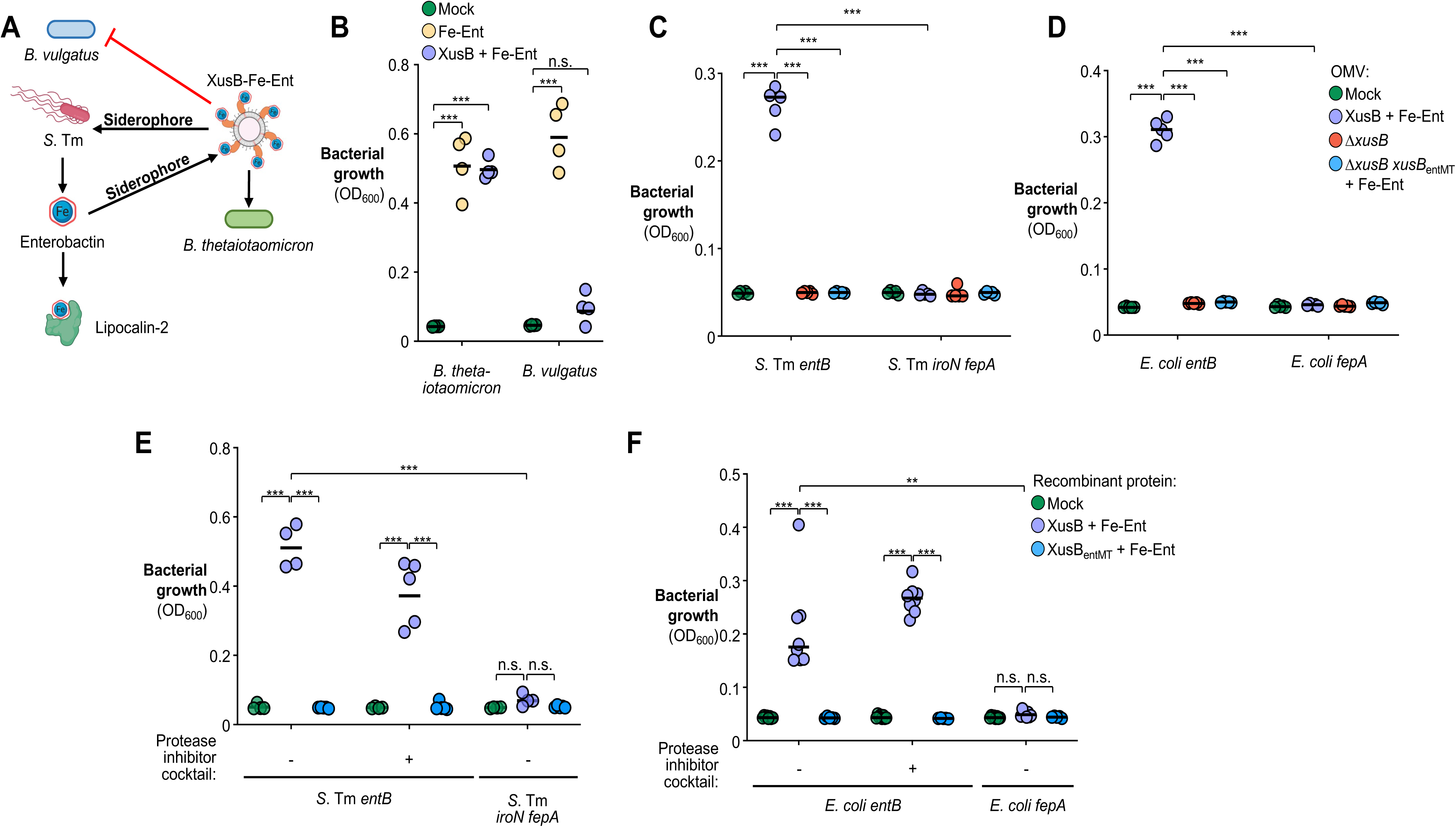
Role of the XusB in interspecies competition for siderophores. (**A**) Model schematic of XusB in modifying enterobactin accessibility. (**B**) *B. thetaiotaomicron* or *B. vulgatus* were inoculated into iron-starved, semi-defined medium (SDM, 200 μmol/L BPS) supplemented with either free Fe-Ent or Fe-Ent complex with recombinant XusB; bacterial growth was measured 48 hours post-inoculation by OD_600_. (**C-D**) Supernatant from donor *B. thetaiotaomicron* strains grown in iron-limited, SDM was filter-sterilized, loaded with Fe-Ent, washed to remove unbound ligand, and introduced to recipient *S*. Tm (**C**) or *E*. coli (**D**) strains grown in SDM supplemented with 200 μmol/L BPS. (**E-F**) Indicated recombinant XusB protein was loaded with Fe-Ent, washed to remove unbound Fe-Ent, and supplemented *S*. Tm (**E**) or *E*. coli (**F**) strains grown in 200 μmol/L BPS supplemented SDM. Bacteria were cultured for 48 hours, and bacterial growth was measured by OD_600_. Bars represent the geometric mean. **; *P* < 0.01; ***; *P* < 0.001; ns, not statistically significant.

To further explore how XusB contributes to interspecies interactions beyond *Bacteroidetes*, we fed Fe-Ent loaded XusB to an *S*. Tm *entB* mutant strain (*Enterobacteriaceae* family, *Proteobacteria* phylum). The *S*. Tm *entB* mutant is deficient in the production of both enterobactin and salmochelin and can only grow in iron-limited conditions when exogenous siderophores are provided^28, 57, 58^. Notably, the *S*. Tm *entB* mutant grew robustly when the Fe-Ent was complexed with XusB^+^ OMVs (**Fig. 5C**). By contrast, OMVs derived from isogenic Δ*xusB* or Δ*xusB xusB*_entMT_ mutants did not support the growth of *S*. Tm *entB* strain, suggesting that XusB-complexed Fe-Ent, rather than contaminating free Fe-Ent, served as the iron source (**Fig. 5C**). Further, Fe-Ent bound to recombinant XusB protein also promoted *S*. Tm *entB* growth in iron starved media (**Fig. 5E**). Notably, another member of the *Enterobacteriaceae* family, *E. coli* BW25113 (isogenic *entB* mutant) also benefited from XusB-bound enterobactin as the sole iron source (**Fig. 5D&F**). The observed interphylum siderophore cross-feeding was not limited to *B. thetaiotaomicron*, as Fe-Ent complexed with OMV fractions derived from other *Bacteroides* members that encode XusB homologs promoted growth of the *S*. Tm *entB* mutant in iron-limiting media (**Supplementary Fig. 6C**). Of note, co-incubation of XusB with *S*. Tm, or *E. coli* did not lead to appreciable degradation of XusB even in the absence of protease inhibitors (**Supplementary Fig. 6A**), suggesting that the utilization of XusB bound enterobactin by *Enterobacteriaceae* was not due to XusB degradation by extracellular proteases to release Fe-Ent. Consistent with this notion, the addition of a protease inhibitor cocktail failed to block the utilization of XusB-bound enterobactin (**Fig. 5E-F**). A small-scale screen using the Keio collection^59^ identified the outer membrane embedded, TonB-dependent enterobactin transporter FepA^60^ to be necessary for utilizing XusB-bound Fe-Ent (**Fig. 5D&F**). Similarly, FepA and IroN^16^, known enterobactin transporters in *S*. Tm, were also required for scavenging enterobactin bound by XusB (**Fig. 5C&E**). These data suggested that *B. thetaiotaomicron* captures siderophores for itself through a mechanism that may mediate siderophore competition with other *Bacteroides* members, but also could be exploited by *Enterobacteriaceae* such as *S.* Tm and *E. coli*.

### Commensal xenosiderophore acquisition modifies nutritional immunity *in vitro*

Gut commensals play key roles in modulating host-pathogen interactions by either conferring resistance or increasing susceptibility to pathogen colonization by modifying nutrient availability^61^. During *S*. Tm infection, the enteric pathogen produces enterobactin and salmochelin^52^ to scavenge iron for growth^62^. While enterobactin is the highest affinity iron chelator known in nature^14^, it can be sequestered by host protein lipocalin-2, constituting an “inaccessible” enterobactin pool to the pathogen^15, 20^. Notably, *S*. Tm secrets enterobactin into the extracellular milieu that is occupied by a high density of commensal bacteria, suggesting that enterobactin can be bound by commensal siderophore-binding proteins such as XusB (*K_d_* = 148 nM, **Fig. 2B**), albeit with a lower affinity than lipocalin-2 (*K_d_* = 0.41 nM)^15^. While the direct exchange of enterobactin between high-affinity binders such as XusB and lipocalin-2 is thermodynamically favorable, the exchange could occur on a time scale of hours^32^. This process could take significantly longer in reality, considering molecular diffusion and exchange in the intestinal lumen occur at a much slower rate than in pure solvent, due to the presence of dense bacterial population, large molecules, and mucous fluid^63^. This suggests that XusB bound enterobactin may represent a distinct siderophore pool that is less accessible to lipocalin-2 but can be exploited by *S*. Tm (**Fig. 5A**). As such, we hypothesized that secreted commensal siderophore binding protein such as XusB could promote *S*. Tm evasion of nutritional immunity by competing for Fe-Ent with the host protein lipocalin-2.

To measure enterobactin accessibility to *S*. Tm in the presence of nutritional immunity components, we competed the *S*. Tm wild-type strain against a Δ*fepA iroN* mutant in semi-defined medium supplemented with iron chelator BPS and recombinant lipocalin-2 (**Fig. 6A**). Because the Δ*fepA iroN* mutant cannot access enterobactin or salmochelin^16^, we predict that the *S*. Tm wild-type strain will outcompete the mutant in the absence of commensal siderophore binding protein, but the fitness advantage of the wild type strain will be significantly greater when XusB is present due to increases access to enterobactin. As predicted, the *S.* Tm wild-type strain outcompeted the Δ*fepA iroN* mutant to a larger magnitude when XusB was added (**Fig. 6B**). Heat inactivation (HI), but not pretreatment using divalent ion chelator Chelex 100, abolished the added fitness advantage of the *S*. Tm wild-type strain, suggesting that XusB, but not the contaminating iron, is responsible for altering the enterobactin pool (**Fig. 6B**). Further, OMVs collected from the *B. thetaiotaomicron* wild-type strain, but not those derived from the *B. thetaiotaomicron* Δ*xusB* mutant, boosted the fitness advantage of *S*. Tm when lipocalin-2 is present (**Fig. 6C**). These data together indicate that a secreted commensal siderophore binding protein can modify enterobactin competition between *S*. Tm and host nutritional immunity components.

**Figure 6:**
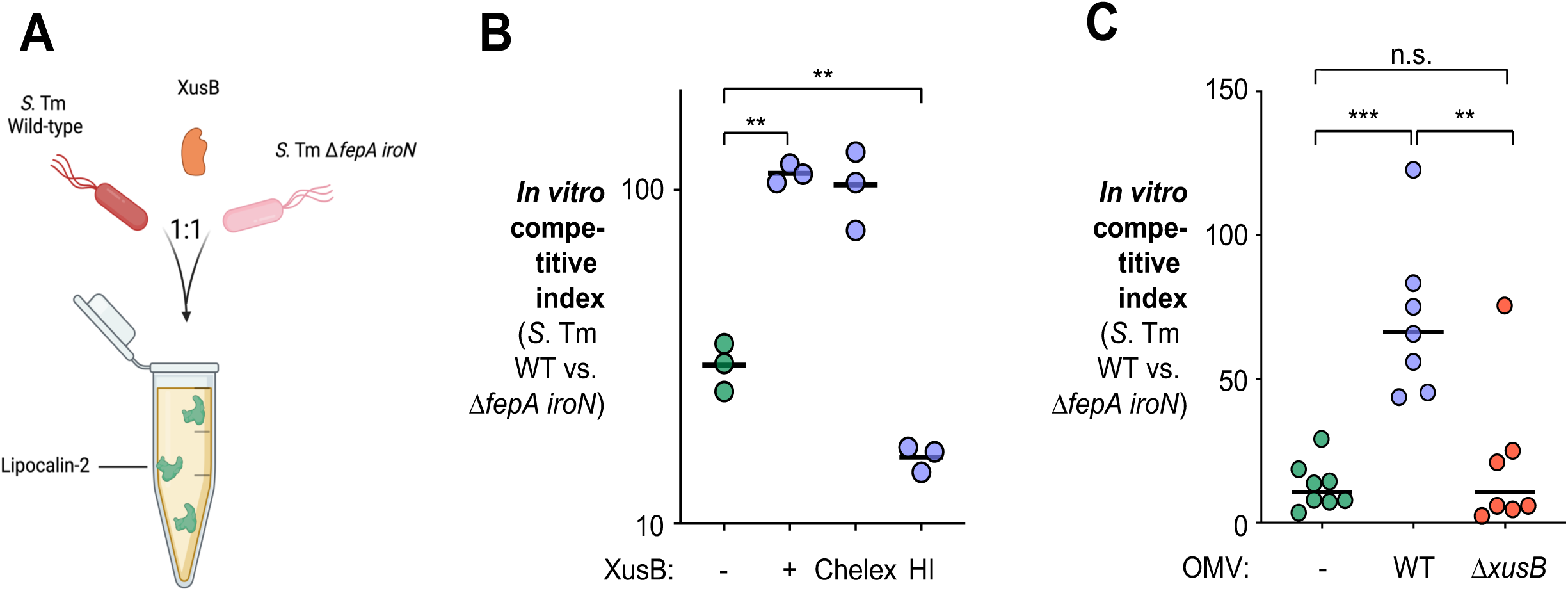
XusB modulates the interactions between *S*. Tm and lipocalin-2 *in vitro*. **(A-C)** An equal mixture of *S*. Tm wild-type strain and an isogenic Δ*fepA iroN* mutant was inoculated into semi-defined medium supplemented with iron chelator BPS and recombinant lipolicalin-2, in the absence or presence of XusB, for 16 hours. Competition index for the *S*. Tm SL1344 wild-type strain vs. the Δ*fepA iroN* mutant strain in the presence of recombinant XusB (**B**) and XusB-containing OMVs (**C**). The competitive index was calculated as the ratio of the two strains in the culture, corrected by the ratio in the inoculum. Chelex: Pretreatment of divalent ion chelating agent Chelex 100. HI: Heat inactivation. Bars represent the geometric mean. **, *P* < 0.01; ***, *P* < 0.001; n.s., not statistically significant.

### Commensal xenosiderophore acquisition modifies nutritional immunity *in vivo*

*S*. Tm relies on catecholate siderophores to overcome iron starvation during infection^20, 23^. While salmochelin evades the sequestration by host protein lipocalin-2^19^, *S*. Tm mutant strains deficient in salmochelin utilization are not completely avirulent in *Lcn2*^+/+^ mice^20^, indicating that a population of enterobactin molecules may escape lipocalin-2 sequestration during infection. One possibility is that commensal siderophore binding proteins such as XusB shield enterobactin from lipocalin-2. By exploiting XusB bound enterobactin, *S*. Tm may recover a pool of enterobactin that is otherwise bound by lipocalin-2. To test this hypothesis, we first explored XusB association with OMVs using a gnotobiotic animal model, which allowed us to measure XusB in the absence of confounding microbes that may express XusB homologs. In this model, we opted to use Swiss Webster mice because Germ-free C57BL/6 mice are highly susceptible to *S*. Tm infection and succumb due to systemic dissemination^64^. We first monoassociated groups of gnotobiotic Swiss Webster mice with either the *B. thetaiotaomicron* wild-type strain or with a Δ*xusB* mutant strain. Seven days later, both groups of mice were challenged with *S*. Tm to induce iron limitation in the gut. We then collected the OMV fraction of filter sterilized homogenate of the intestinal contents and analyzed the proteome (**Fig. 7A**). Only the OMV fraction from mice associated with *B. thetaiotaomicron* wild-type strain yielded XusB-specific peptide fragments, consistent with the idea that *B. thetaiotaomicron* produces OMV-associated XusB under iron-limiting conditions *in vivo* (**Fig. 7B**).

**Figure 7:**
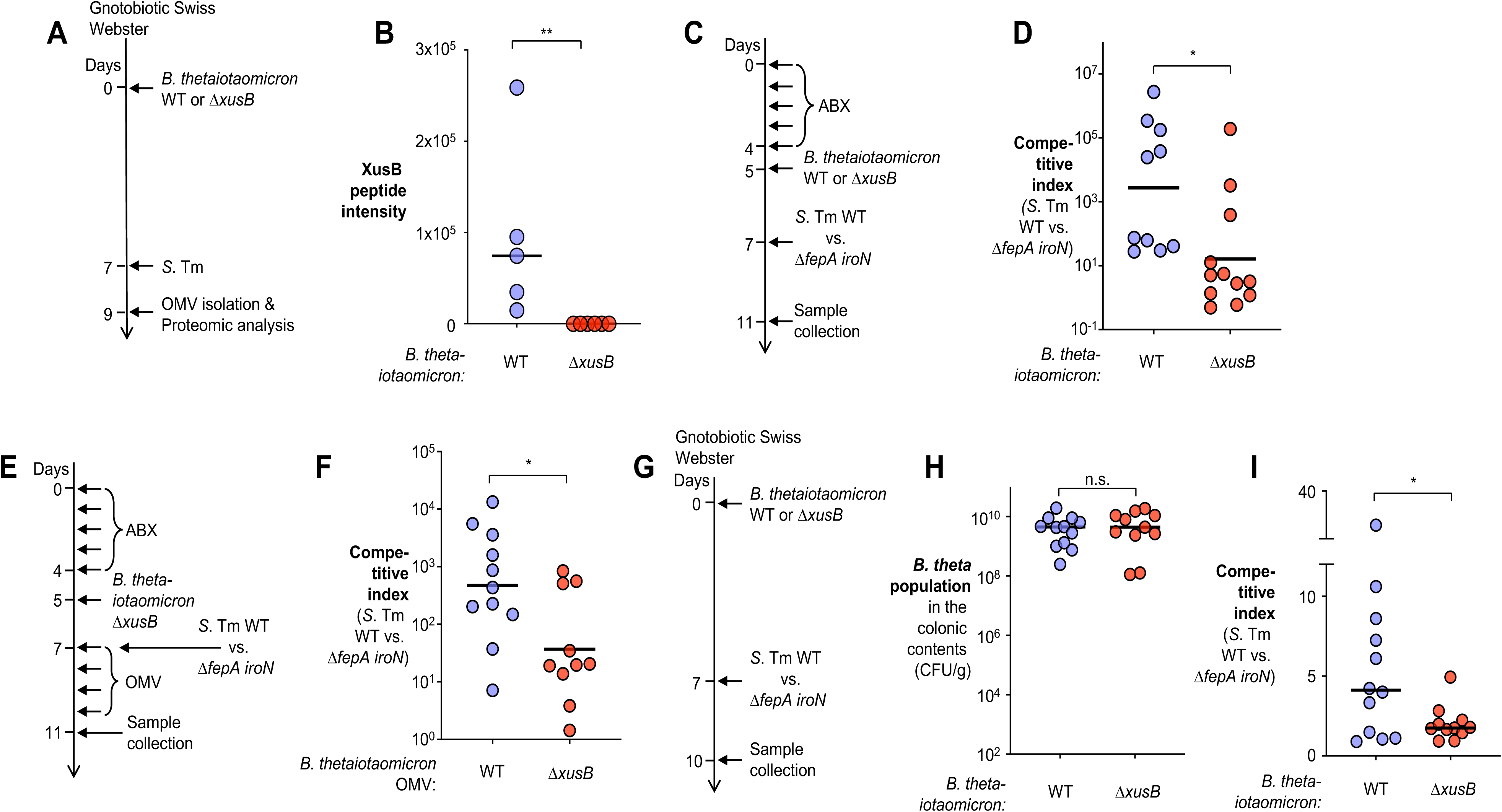
Role of XusB in pathogen-host nutritional immunity interactions during *S*. Tm infection. **(A-B)** Groups of gnotobiotic Swiss Webster mice were monoassociated with either the *B. thetaiotaomicron* wild-type strain (*N* = 5) or the isogenic Δ*xusB* mutant (*N* = 6) for 7 days. Mice were then challenged with the *S*. Tm SL1344 for 2 days. The outer membrane vesicle fraction of the intestinal contents was collected, and the proteome profiled using UHPLC-MS/MS. (**A**) Schematic representation. (**B**) The peptide abundance of XusB measured by UHPLC-MS/MS. (**C-D**) Groups of C57BL/6 mice were treated with a cocktail of antibiotics, followed by intragastrical inoculation of either the *B. thetaiotaomicron* wild type strain (*N* = 10) or an isogenic Δ*xusB* mutant (*N* = 12). Mice were inoculated with an equal mixture of the *S*. Tm SL1344 wild-type strain and the isogenic Δ*fepA iroN* mutant strain. (**C**) Schematic representation of the experiment. 4 days later, cecal contents were collected, and the competitive index (**D**) for the *S*. Tm SL1344 wild-type strain vs. the Δ*fepA iroN* mutant strain was determined. (**E-F**) Groups of C57BL/6 mice were treated with a cocktail of antibiotics, followed by intragastrical inoculation of a Δ*xusB* mutant strain. Two days later, mice were then inoculated with an equal mixture of the *S*. Tm SL1344 wild-type strain and the isogenic Δ*fepA iroN* mutant strain, followed by intragastrical administration of an equal amount of OMVs derived from the *B. thetaiotaomicron* wild type strain (*N* = 11) or the isogenic Δ*xusB* mutant strain (*N* = 10) daily. (**E**) Schematic representation. 4 days later, cecal contents were collected, and the competition index (**F**) for the *S*. Tm SL1344 wild-type strain vs. the Δ*fepA iroN* mutant strain was determined. **(G-I)** Groups of gnotobiotic Swiss Webster mice were monoassociated with either the *B. thetaiotaomicron* wild-type strain (*N* = 12) or the isogenic Δ*xusB* mutant (*N* = 11) for 7 days. Mice were then challenged with an equal mixture of the *S*. Tm SL1344 wild-type strain and the isogenic Δ*fepA iroN* mutant strain for 3 days. (**G**) Schematic representation of the experiment. The abundance of *B. thetaiotaomicron* (**H**) and the competition index (**I**) for *S*. Tm SL1344 wild-type strain vs. the Δ*fepA iroN* mutant strain in the cecal contents were determined. Bars represent the geometric mean. *, *P* < 0.05; **, *P* < 0.01; n.s., not statically significant.

Next, we explored if commensal siderophore acquisition modifies nutritional immunity *in vivo*. To this end, we colonized conventional C57BL/6 mice with the *B. thetaiotaomicron* wild-type strain or an isogenic Δ*xusB* mutant. We then inoculated these mice with an equal mixture of the *S*. Tm wild-type strain and an isogenic Δ*fepA iroN* mutant defective in utilizing both catecholate siderophores. The competitive infection design allows a controlled comparison as the wild-type and mutant bacteria experience the same environment in a single mouse. Four days post inoculation, we collected intestinal contents, quantified the abundance of *S*. Tm strains by plating on selective media and calculated the competitive index as the ratio of the *S.* Tm wild-type vs. the Δ*fepA iroN* strains (**Fig. 7C**). We predict that the *S*. Tm wild-type strain will outcompete the mutant in both groups, but to different magnitudes. If *S*. Tm can access XusB-bound enterobactin, the competitive index will be greater in mice colonized with XusB-producing *B. thetaiotaomicron* than those with the Δ*xusB* mutant. Indeed, the *S*. Tm wild-type strain outcompeted the Δ*fepA iroN* mutant to a greater magnitude in mice colonized with the wild-type *B. thetaiotaomicron* compared to animals associated with the Δ*xusB* mutant (**Fig. 7D**). No significant difference in the levels of nutritional immunity components such as lactoferrin, lipocalin-2, or calprotectin were noted (**Supplementary Figure. 7A**).

Commensal bacteria such as *B. thetaiotaomicron* could modify the susceptibility to *S*. Tm by altering the availability of nutrients other than iron in the gut^25, 65–67^. Because the *B. thetaiotaomicron* wild-type strain displays higher fitness compared to the Δ*xusB* mutant (**Fig. 4B**), we sought to assess the role of XusB in nutritional immunity in the absence of the fitness difference between the two *B. thetaiotaomicron* strains. To this end, we colonized two groups of C57BL/6 mice with the *B. thetaiotaomicron* Δ*xusB* mutant strain, followed by a challenge of an equal mixture of the *S*. Tm wild-type strain and a Δ*fepA iroN* mutant. We then intragastrically administered equal amount of OMVs harvested from wild-type *B. thetaiotaomicron* (WT-OMVs), or an isogenic mutant lacking XusB (Δ*xusB*- OMVs) (**Fig. 7E**). In these mice, there was no significant difference in *B. thetaiotaomicron* fitness or levels of inflammatory markers (**Supplementary Figure. 7B and 8A**). In contrast, while the *S*. Tm wild-type strain outcompeted the Δ*fepA iroN* mutant in mice treated by oral gavage of Δ*xusB*-OMVs, the fitness advantage over the double mutant was significantly greater in mice that received WT-OMVs (**Fig. 7F**), consistent with the notion that XusB alters enterobactin accessibility *in vivo*.

To further exclude the influence of other microbes in the gastrointestinal tract, we monoassociated groups of gnotobiotic Swiss Webster mice with the *B. thetaiotaomicron* wild-type strain or the Δ*xusB* mutant. After seven days, we intragastrically challenged both groups of mice with an equal mixture of *S*. Tm wild-type strain and the Δ*fepA iroN* mutant. Three days post-infection, we collected the intestinal contents and determined the abundance of *B. thetaiotaomicron* and *S*. Tm strains. In contrast to the conventional mice, both *B. thetaiotaomicron* strains were recovered at comparable levels (**Fig. 7H**), likely due to the lack of competition with other gut microbes and a ∼10-fold higher iron content in the gastrointestinal tract of Germ-free animals compared to conventional mice^68^. The latter phenotype is likely due to the dysregulated iron absorption and retention in Germ-free animals as a result of the absence of microbial signals ^68–70^, a defect that cannot be fully restored by short-term monoassociation of a defined bacterial consortium^68^. Despite the increased iron availability, the fitness advantage of the *S*. Tm wild-type strain over the Δ*fepA iroN* mutant was significantly greater in mice associated with the *B. thetaiotaomicron* wild-type strain compared to those associated with the Δ*xusB* mutant strain (**Fig. 7I**). Analysis of transcript levels of inflammation markers revealed a marginal difference in lactoferrin, while the levels of other nutritional immunity components such as lipocalin-2 and calprotectin were undistinguishable between the two treatment groups (**Supplementary Figure. 7C**). Taken together, these experiments demonstrate that iron acquisition by a commensal bacterium modifies host nutritional immunity during *Salmonella* infection.

## Discussion

The commensal bacterial community maintains a mutualistic relationship with its host and provides pivotal benefits ranging from aid in digestion to promoting the maturation of the immune system^7^. For example, commensals, including members of the *Bacteroidetes* phylum, produce short-chain fatty acids and bio-transform primary bile acids to regulate T cell differentiation^71, 72^, and prevent host-to-host spread of pathogens by signaling the immune system^73^. This ecosystem constantly faces perturbations, including changes in diet, the use of antibiotics, and inflammatory diseases, impacting the structure of the gut microbial community. The altered community profiles, typically characterized by reduced diversity and depletion of commensal species, are often associated with exacerbation of disease outcomes^8, 74–78^. As such, microbial resilience is key to maintaining the structural and functional stability of the gut microbiome over time^3, 7, 79^.

Intestinal inflammation is frequently accompanied by changes in the intestinal immune and nutritional landscape^80–83^, including markedly decreased bioavailability of essential micronutrients such as iron^11^. While the molecular mechanisms underlying the adaptation to altered nutritional and immune landscapes as key resilience strategies are beginning to be uncovered^74, 84^, how commensal bacteria survive metal starvation remains to be defined. Here, we identified XusB, a surface-exposed lipoprotein that is enriched onto outer membrane vesicles, functions as an integral part of the siderophore acquisition machinery required for *B. thetaiotaomicron* to sustain resilience during infectious colitis. This represents a distinct capture mechanism compared to the canonical siderophore uptake machinery in Gram-negative bacteria, which involves an outer membrane TonB-dependent receptor that transports iron-laden siderophores into the periplasmic space, where a siderophore binding protein relays the siderophore onto an ABC-transporter for transport into the cytoplasm^85^. Xus-mediated siderophore uptake also differs from the mechanisms in Gram-positive bacteria, which typically employ an outer membrane-anchored siderophore binding protein coupled with an ABC-transporter instead of a TonB-dependent receptor for siderophore transport^86^. In contrast, *B. thetaiotaomicron*, a Gram-negative bacterium, requires an extracellular siderophore binding protein and a TonB-dependent receptor for the initial steps of siderophore uptake. In addition to the mechanism of action, XusB adopts a unique seven-blade β-propeller fold that is structurally distinct from siderophore binding proteins in Gram-positives^87^, and known extracellular nutrient binding proteins in *Bacteroidetes*^88–91^(**Fig. 2C&G**, **Supplementary Fig. 3A**). Of note, XusB is not unique to *B. thetaiotaomicron*, as we identified homologs of XusB in diverse members of *Bacteroidetes* that are predicted to be structurally conserved (**Fig. 2G** and **Supplementary Fig. 2**). Although these homologs are predicted to adopt the conserved seven-blade β-propeller fold as found in XusB, they are predicted to harbor distinct ligand binding pockets (**Supplementary Fig. 3B**). While the ligands for XusB homologs remains to be defined, our results indicate that members of the *Bacteroidetes* phylum employ a unique transport machinery to capture nutrients including xenosiderophores.

Some strains that encode XusB homologs may capture xenosiderophores in a selfish manner, suggesting that species-specific factors are required to extract siderophores from XusB or its homologs. Because cross-feeding of Fe-Ent from OMV associated XusB requires cell-surface exposed XusB in the recipient cells, it is plausible that species-specificity is conferred by XusB-XusB interactions, although we cannot exclude the possibility that species-specific factor(s) dock the OMVs to the recipient cells for subsequent enterobactin extraction. While the underlying molecular basis remains to be investigated, our data suggest that XusB and its homologs can shape the gut microbial community structure under inflammatory conditions by contributing to the competition of xenosiderophore in related species.

Remarkably, XusB-bound enterobactin is readily accessible to phylogenetically distant bacteria such as *S*. Tm and *E. coli* (phylum *Proteobacteria*), the archetypical examples of microbial competition for iron with the host nutritional immunity^20–22^. Moreover, we show that XusB may modify nutritional immunity as enterobactin bound by XusB is less accessible to lipocalin-2 but remains accessible to *S*. Tm and *E. coli*. One contributing factor is that both XusB and lipocalin-2 are high-affinity enterobactin binders, and the transfer of enterobactin from XusB to lipocalin, while thermodynamically favorable, may take too long to occur to impact microbial growth in the gut. Additionally, *Bacteroidetes* such as *B. thetaiotaomicron* are lumen-dwelling while lipocalin-2 is produced by epithelial cells and infiltrating myeloid cells^20, 92^. Therefore, it is plausible that XusB and lipocalin-2 may not be present in high enough concentrations in the same geographical location for the competition to occur. Further, the presence of a dense bacterial community and large molecules such as mucus and glycans likely create physical hindrance, viscous drag, transient traps, and non-specific interactions, which together reduce the rate of molecular diffusion by orders of magnitudes compared to a pure solvent^63^. As a population of the enterobactin produced by pathogens (e.g. *S*. Tm) are sequestered by lipocalin-2^19, 20^, accessing enterobactin through the commensal siderophore binding proteins can promote further evasion of nutritional immunity. Consistent with this notion, the *S*. Tm wild-type strain displayed an enhanced fitness advantage in the presence of XusB. As multiple *Bacteroidetes* members encode homologs of XusB, our results suggest that commensal iron acquisition represents a previously unrecognized layer to host nutritional immunity.

## STAR METHODS

### Iron-laden siderophores

The iron-free siderophores enterobactin (Sigma-Aldrich) and salmochelin S4 (Dr. Winkelmann, University of Tubingen, Germany) were dissolved in trace element grade dimethyl sulfoxide (Sigma-Aldrich) at a concentration of 2 mmol/L. Sterile 1 or 2 mmol/L Fe(III) chloride (Sigma-Aldrich) was incubated with the iron-free siderophores at 1:1 (v/v) overnight at 4 °C to obtain a 1 mmol/L siderophore stock solution at 50% or 100% saturation, respectively.

For Fe^58^ transfer experiments, an equal volume of 2 mM FeCl_3_ (the stock was made 100 mM in 12N HCl) and 2 mM apo-enterobactin was mixed at RT for 5 minutes. The mixture was then diluted into 0.3 M Tris-HCl (pH 8.0) to give a final concentration of 333 μM.

### Bacterial strains and growth conditions

The bacterial strains used in this study are listed in the Key Resource Table. *B. thetaiotaomicron, B. salyersiae,* and *B. vulgatus* were grown anaerobically (90% N_2_, 5% CO_2_, 5% H_2_; vinyl anaerobic chamber, Coy Lab) in brain-heart-infusion supplemented (BHIS) media (0.8% brain heart infusion from solid, 0.5% peptic digest of animal tissue, 1.6% pancreatic digest of casein, 0.5% sodium chloride, 0.2% glucose, 0.25% disodium hydrogen phosphate, 0.005% haemin, 0.0001% /vitamin K, pH 7.4) or BHIS plates (BHIS broth, 15 g/L agar) containing 50 μg/mL gentamycin (Gen) or 15 μg/mL chloramphenicol (Cm) or 25 μg/mL erythromycin (Erm) for 2 days at 37 °C.

For OMV harvest and siderophore-cross-feeding, single colonies of indicated strains were cultured anaerobically for 24 hours in BHIS, and the electron dense layer (EDL) fraction of the culture was purified using gradient centrifuge^93^ before subculturing into modified semi-defined media (SDM)^94^ (1.5 g/L KH_2_PO_4_, 0.5 g/L NH_4_SO_4_, 0.9 g/L NaCl, 150 mg/L L-methionine, 5 μg/L vitamin B_12_, 1 mg/L resazurin, 1 g/L tryptone, 0.2% NaHCO_3_, 0.005% protoporphyrin IX) for 16 hours.

For OMV harvest, bacterial culture was stimulated with 200 μM of bathophenanthroline disulfonate (BPS) for 3 hours, pelleted at 5,000 x*g* for 15 minutes at 4 °C, and the supernatant was passed through 0.22 μM polyethersulfone (PES) membranes (PALL Corporation) to remove cells and debris. The sterility of the vesicle containing filtrates was confirmed by plating onto BHIS agar. Iron-laden enterobactin or salmochelin (at 50% saturation) were added at a final concentration of 0.5 μmol/L and 2 μmol/L, respectively, and the binding of siderophores to XusB was allowed to proceed for 2 hours at RT (25 °C). The cell-free culture supernatant was then centrifuged at 150,000 x*g* (Beckman TLA-100.3 or 45 Ti) for 2 hours at 4 °C to recover the OMV fraction (pellet) and the supernatant fraction. Both fractions were then incubated with 2 μM of 100% saturated Fe-Ent for 1 hour at RT. The unbound Fe-Ent was removed by washing the OMV and supernatant fractions using ultrafiltration (Amicon Ultra, 30 kDa MWCO, Sigma Aldrich) for 4 times. For OMV cross-feeding experiments, the OMVs fraction was resuspended in SDM before adding to recipient cells. Recipient cells were cultured in BPS-supplemented SDM anaerobically. The inoculum for these experiments was 10^4^ CFU/ml. For *in vitro* competition assays, each bacterial strain was inoculated at 5×10^4^ CFU/ml. To confirm that other trace elements are not responsible for the growth defect observed in BPS-supplemented media, ammonium iron (III) citrate was supplemented at a concentration of 200 μmol/L. In experiments where purified, recombinant XusB instead of OMVs was added, the recombinant proteins were first incubated with Chelex 100 Resin (1 g/L, Bio-Rad Laboratories) for 3 hours at 4 °C to remove possible contaminating iron. For *in vitro* competition assays described in Fig. 6, the *S*. Tm wild-type and the Δ*fepA iroN* strains were inoculated at 5×10^4^ CFU/ml in SDM supplemented with 50 μM BPS and 43.48 nM of LCN-2 (R&D Systems), in the absence or presence of 5.7 μM of recombinant XusB.

*E. coli* and *S*. Typhimurium strains were routinely cultured in LB broth (10 g/L tryptone, 5 g/L yeast extract, 10 g/L sodium chloride) or on LB plates (LB broth, 15 g/L agar) at 37 °C, when appropriate, with antibiotics were added at the following concentrations: 100 µg/mL streptomycin (Strep), 100 µg/mL carbenicillin, or 100 µg/mL kanamycin (Kan).

### Plasmids

All the primers and plasmids used in this study are listed in the Key Resource Table. Suicide plasmids were routinely propagated in *E. coli* DH5α λ*pir* or S17 λ*pir*. The flanking regions of the *B. thetaiotaomicron* genes BT_2064 (XusB), BT_2064-65 were amplified and assembled into pExchange-*tdk* using the Gibson Assembly Cloning Kit (New England Biolab, Boston) to give rise to pKR112 and pWZ1138 respectively. To complement the BT_2064 deletion, the promoter of BT_2065 (the first gene in the BT_2063/4/5 (XusABC) operon) and the ORF of WT BT_2064 were synthesized (Twist Bioscience) and assembled into pKI, pNBU2-3xFlag or pNBU2-3XHA using the Gibson Assembly Cloning Kit. For point mutations of XusB, the promoter of BT_2065 and the mutated ORF of BT_2064 were synthesized (Twist Bioscience) and assembled into pNBU2-3xFlag or pNBU2-3XHA.

The flanking regions of the *S.* Typhimurium *fepA*, *iroN* genes were amplified and ligated into pGP706 to generate pWZ1077 and WZ1074. Relevant plasmid inserts were verified by Nanopore sequencing (Plasmidsaurus, USA). In bacterial competition experiments, *B. thetaiotaomicron* strains were marked by inserting a chloramphenicol resistant cassette into a neutral locus in the genome^28^. *S*. Tm strains were differentially marked by introducing the low-copy number plasmid pWSK29 or pWSK129 through electroporation (e.g., *S*. Typhimurium + pWSK29, WZ1110 and *S*. Typhimurium Δ*fepA iroN* + pWSK29, WZ1108)^95^.

To generate expression systems for recombinant HisTagged, truncated XusBΔN (residues 41-464), the mature coding region of WT XusB was amplified by PCR, and fragments were subsequently cloned via the Gibson Assembly Cloning Kit (New England Biolab, Boston) into pBG106 expression vector (Center for Structural Biology, Vanderbilt University) linearized using NdeI. All constructs were verified by Sanger or Oxford Nanopore Sequencing.

### Construction of mutants by allelic exchange

All bacterial mutant strains constructed using the method below are listed in the Key Resource Table. For *B. thetaiotaomicron* mutants, suicide plasmid pExchange-*tdk* containing the flanking regions of genes of interest was conjugated using *E. coli* S17-1 λ*pir* as the conjugative donor strain into the *B. thetaiotaomicron*. Exconjugants with suicide plasmid integrated into the recipient chromosome were selected on blood plates supplemented with appropriate antibiotics. 5-fluoro-2-deoxy-uridine (FudR, 200 µg/mL in BHIS) plates were used to select for the second crossover event. To create the *S.* Typhimurium Δ*fepA* mutant, pWZ1077 conjugated into the *S.* Typhimurium SL1344 using *E. coli* S17-1 λ*pir*. Exconjugants were recovered on LB agar containing appropriate antibiotics. The second crossover event was selected using sucrose plates (8 g/L nutrient broth base, 5 % sucrose, 15 g/L agar), thus creating WZ1093. Deletion of the target gene was confirmed by PCR. Similar strategies were used to construct the Δ*iroN* mutant in *S.* Typhimurium Δ*fepA* background, resulting in the *S.* Typhimurium Δ*fepA iroN* (WZ1096).

### Animal models

All experiments were conducted in accordance with the policies of the Institutional Animal Care and Use Committee at Vanderbilt University Medical Center. C57BL/6J wild-type (cat# 000664), originally obtained from Jackson Laboratory (Bar Harbor), were procured, bred, and housed under specific pathogen-free conditions at Vanderbilt University Medical Center.

Seven to nine-week-old male and female mice were semi-randomly assigned into treatment groups before the experiment. Antibiotic cocktails (5 mg of each ampicillin (Sigma-Aldrich), metronidazole (Sigma-Aldrich), vancomycin (Chem Impex International), and neomycin (Sigma-Aldrich) per mouse) or mock treatment (water) were administered by oral gavage daily for 5 days. After antibiotic treatment, fecal pellets were collected and tested for bacterial growth on blood agar and blood agar supplemented with 50 μg/mL gentamycin. Only mice with no detectable bacterial growth on both media were included in the study to allow for quantification of experimentally-introduced *B. thetaiotaomicron* strains in luminal content and feces. At day 5, mice were then inoculated with 1 x 10^9^ CFU of the indicated *B. thetaiotaomicron* strains or remained uninfected. In competitive colonization experiments, animals were inoculated with an equal mixture of 0.5 x 10^9^ CFU of the *B. thetaiotaomicron* wild-type strain and 0.5 x 10^9^ CFU of the indicated mutants. Two days later, mice were challenged by 1 x 10^5^ CFU of the *S*. Typhimurium strain SL1344 for 4 days (**Fig. 4**). For the experiments involving *S*. Typhimurium competition, mice were challenged by 0.5 x 10^5^ CFU of *S*. Typhimurium strain SL1344 wild-type strain and 0.5 x 10^5^ CFU of indicated isogenic mutant for 4 days.

For *in vivo* OMV administration, bacterial culture was centrifuged, and bacterial cells and debris were removed using PES membrane filters as described above. OMVs were collected by centrifuging filtrate at 200,000 x*g* (Beckman 45 Ti) at 4 °C for 2 hours. The quantity of OMVs was normalized by total protein content (Bradford Assay, Bio-Rad laboratories). OMVs with 20 μg total protein in 100 μL PBS were administered intragastrically.

For all experiments, fecal pellets were collected at the indicated time points. After euthanasia, cecal and colonic tissue was collected, flash-frozen and stored at −80 °C for subsequent mRNA analysis or fixed in 10% formalin for histopathological analysis. For culture-dependent quantification of bacterial load, colonic and cecal contents were harvested in sterile PBS, and the load of *B. thetaiotaomicron*, *S*. Typhimurium, or *E. coli* were quantified by plating serial-diluted intestinal contents on selective agar.

### Gnotobiotic mouse experiments

Germ-free Swiss-Webster mice were purchased from The National Gnotobiotic Rodent Resource Center and maintained in Sentry SPP Mouse caging system (Allentown, USA) on a 12-hour light cycle. Male and female mice were randomized and orally gavaged with 1×10^9^ CFU of indicated *B. thetaiotaomicron* strains. Seven days later, mice were challenged with 1 x 10^5^ CFU of indicated *S*. Typhimurium SL1344 strains for 3 days. For the experiments involving *S*. Typhimurium competition, mice were challenged by 0.5 x 10^5^ CFU of *S*. Typhimurium strain SL1344 wild-type strain and 0.5 x 10^5^ CFU of indicated isogenic mutant for 3 days. After euthanasia, colonic and cecal contents were harvested in sterile PBS, and the load of *B. thetaiotaomicron* and *S*. Typhimurium were quantified by plating serial-diluted intestinal contents on selective agar.

For OMV isolation from intestinal contents, intestinal contents were harvested and homogenized in sterile PBS. A 100 μL aliquot of the homogenates was removed and plated on selective agar to enumerate bacteria load. The rest of the homogenate was centrifuged at 5,500 x*g*, 4 °C for 15 min. The supernatants were filter-sterilized using 0.22 μm PES membranes (PALL) and the sterility of the filtrate was confirmed by plating onto BHIS plates. The filtrate was centrifuged (200,000 x*g* at 4 °C for 2 hours) in a TLA-100.3 rotor (Beckman Instruments). After centrifugation, the supernatant was removed, and the presence of XusB in the OMV fraction was determined using a proteomics approach. Briefly, proteins in the OMV pellets were dissolved in 50 μL of 5% SDS was added to each protein pellet, following which 2 μL of 500 mM TCEP was added for disulfide bond reduction and incubated at 56 °C for 30 min. After cooling, 2 μL of 500 mM iodoacetamide was added and samples were alkylated for 30 min at room temperature in the dark. 5.4 μL of 12% phosphoric acid was added followed by 300 uL of S-Trap binding buffer (100 mM TEAB in 90% MeOH), and each sample was loaded onto an S-Trap column (Protifi). Samples were digested overnight with 1µg of trypsin (Pierce) at 37 °C. Following digestion, the peptide eluates were cleaned using an Oasis HLB microelution plate (Waters), dried in a SpeedVac and reconstituted in a 2% acetonitrile, 0.1% TFA buffer for analysis.

Peptides were injected onto an Orbitrap Fusion Lumos mass spectrometer coupled to an Ultimate 3000 RSLC-Nano liquid chromatography system (Thermo Fisher). Samples were injected onto a 75 um i.d., 75-cm long EasySpray column (Thermo Fisher) and eluted with a gradient from 1-28% buffer B over 90 min. Buffer A contained 2% (v/v) ACN and 0.1% formic acid in water, and buffer B contained 80% (v/v) ACN, 10% (v/v) trifluoroethanol, and 0.1% formic acid in water. The mass spectrometer operated in positive ion mode with a source voltage of 1.8 kV and an ion transfer tube temperature of 275 °C. MS scans were acquired at 120,000 resolution in the Orbitrap and up to 10 MS/MS spectra were obtained in the ion trap for each full spectrum acquired using higher energy collisional dissociation (HCD) for ions with charges 2-7. Dynamic exclusion was set for 25 s after an ion was selected for fragmentation.

Raw MS data files were analyzed using Proteome Discoverer v2.4 (Thermo Fisher), with peptide identification performed using Sequest HT searching against the mouse and *Bacteroides thetaiotaomicron* protein databases from UniProt. Precursor and fragment mass tolerances of 10 ppm and 0.6 Da were specified, and three missed cleavages were allowed. Carbamidomethylation of Cys was set as a fixed modification and oxidation of Met set as a variable modification. The false-discovery rate (FDR) cutoff was 1% for all peptides. The proteomics data has been deposited in the MassIVE database under the accession number MSV000091422.

### Targeted quantification of mRNA levels in intestinal tissue and contents

Colonic or cecal tissue was homogenized in a bead beater (FastPrep 24, MP Bio) and RNA was extracted using TRI reagent (Molecular Research Center, Cincinnati). DNA contamination was removed using the Turbo DNA-free Kit (Ambion, USA) per the manufacturer’s recommendations. SuperScript VILO cDNA Synthesis Kit (Thermo Fisher, USA) was used to generate cDNA. Real-time PCR was performed using PowerUp SYBR Green Master Mix (Applied Biosystem, USA), data were acquired in a CFX maestro 2.3 (Bio Rad, USA). The final concentration of primers listed in the Key Resource Table was 250 nM. Target gene transcription of each sample was normalized to the respective levels of *Gapdh* (mouse) or *gyrB* (bacterial) mRNA.

### Inductively Coupled Plasma Mass Spectrometry

For measurement of XusB enterobactin mediated iron transfer, XusB containing OMVs were concentrated using Amicon Ultra (30 kDa MWCO, Sigma-Aldrich), followed by incubation with Fe-laden enterobactin at RT for 3 hours. Unbound Fe-laden enterobactin was removed by ultrafiltration (Amicon Ultra, 30 kDa MWCO, Sigma-Aldrich), and the retentate was incubated with indicated *B. thetaiotaomicron* strains anaerobically in SDM for 16 hours at 37 °C. Bacterial cells were then collected and digested using freshly prepared trace element grade 50% nitric acid (Thermo Fisher Scientific, USA) and 6% H_2_O_2_ (Thermo Fisher Scientific, USA). Sample digestion was allowed to proceed for 48 hours to ensure the complete dissolution of the metals to be analyzed. Samples were diluted in metal-free H_2_O to achieve a 3% nitric acid final concentration, followed by centrifugation to remove any particulates. The supernatant was analyzed for iron by inductively coupled plasma mass spectrometry (ICP-MS) using Agilent 7700 single quadrupole ICP-MS (Agilent Technologies). The measurement was repeated three times for each sample and the Fe (^56^Fe) content was normalized to the total sulfur (^34^S) content of the cells.

### Bacterial cell fractionation assay

Single colonies of *B. thetaiotaomicron* wild type strain were cultured in BHIS for 16 hours before subculture in SDM until reached the exponential phase (OD_600_= 0.8). BPS was added at 200 μM and the culture was incubated anaerobically for 3 hours at 37 °C. Bacterial culture was pelleted at 5,000 x*g* for 15 minutes at 4 °C, and the supernatant was passed through 0.22 μM polyethersulfone (PES) membranes (PALL Corporation) to remove cells and debris. The culture supernatant was concentrated (Amicon Ultra, 30 MWKO, Sigma-Aldrich) and centrifuged as described above to separate the OMV fraction (pellet) and supernatant. For the fractionation of bacterial cells, bacterial pellet was resuspended in lysis buffer (50 mM Tris pH 8, 5 mM EDTA, 2 mM PMSF, 10% glycerol)^88^ supplemented with 1X protease inhibitor cocktail (Halt Protease Inhibitor Cocktail, Thermo Fisher) per the manufacturer’s recommendation. Bacterial cells were then lysed at 4 °C by sonication (40% Amps; 15 s ‘on’ and 30 s ‘off’; 3 min total, Brandson, USA). The lysate was passed through 0.22 μM PES membrane to remove intact cells before centrifugation at 13,200 x*g* for 30 minutes at 4 °C. The resulting membrane fraction (insoluble) and cytoplasm fraction (soluble) were resolved in SDS-PAGE gels and the presence of XusB was probed using a custom-made rabbit anti-XusB polyclonal antibody (Cocalico Biologicals, Reamstown, PA, USA).

### Protease K assay

Single colonies of *B. thetaiotaomicron* wild-type strain was cultured in BHIS for 16 hours before subculture in SDM until reached the exponential phase (OD600 = 0.8). BPS was added at 200 μM and the culture was incubated anaerobically for 3 hours at 37°C. The bacterial culture was pelleted at 5,000 x*g* for 15 minutes at 4 °C, and the pellet was resuspended in PBS supplemented with the protease K (0, 10, 50 or 100 µg/mL, Thermo Scientific). Protease K digestion was allowed to proceed for 1 hour at 37 °C. The reaction was stopped by washing bacterial cells 3 times with PBS supplemented with 1X protease inhibitor cocktail (Halt Protease Inhibitor Cocktail, Thermo Fisher). Bacterial cells were then collected and lysed using BugBuster reagent (Sigma-Aldrich, USA), clarified protein lysate was resolved in an SDS-PAGE gel, and the presence of XusB and RpoB (cytoplasmic marker) was probed using specific antibodies.

### Recombinant XusBΔN protein purification

His-tagged XusBΔN was overexpressed in *E. coli* C41 (DE3) cells. Cells were grown at 37 °C in the Lysogeny Broth (LB) medium containing 100 μg/ml kanamycin until reaching an optical density at 600 nm (OD_600_) of 0.6. The expression of recombinant protein was induced with 1 mM (final concentration) of isopropyl-β-D-thiogalactopyranoside for 6 hours at 37 °C.

Cells were resuspended in lysis buffer (20 mM Tris, pH 8, 150 mM NaCl, 40 mM imidazole, EDTA-free protease inhibitor tablets (Roche), and lysozyme (1 mg per 10 mL buffer). Cells were lysed by sonication for 15 min, and the resulting lysate was centrifuged at 13,000 x*g* for 20 min at 4 °C to remove cellular debris. The resulting supernatant was injected into a HisTrap FF (Cytiva Life Sciences) column at room temperature and eluted with a gradient from 0 to 500 mM imidazole. Eluted protein was loaded onto a gel filtration Cytiva Superdex S200 gel filtration column in 20 mM Tris-HCl (pH 8.0), 150 mM NaCl. The fraction (single peak) containing monomeric XusBΔN was collected and concentrated using Amicon Ultra Centrifugal Filter Units (10 kDa MWCO, Millipore Sigma).

### Chromatographic analysis of Fe-Ent binding by XusB

Recombinant XusBΔN was incubated with 100% saturated Fe-Ent at a 1:1 molar ratio for 1 hour at RT (25 °C) and analyzed by by size-exclusion chromatography with multi-angle light scattering (SEC-MALS). XusB and Fe-Ent were traced using absorbance at 280 nm (aromatic amino acids) and 550 nm^43^, respectively.

### Crystallization and structure determination

A XusBΔN stock (20 mg/mL; 20 mM Tris, pH 8, 150 mM NaCl) was used for all crystal screens. Initial screening was carried out using a Mosquito Crystal Robot (STP LabTech) with 300 nL of protein solution plus 300 nL of reservoir solution in 96-well format plates (MRC two-well crystallization microplate; Hampton Research). Scaled-up crystallization experiments were carried out in VDX 24-well crystallization trays (Hampton Research) using the hanging drop vapor diffusion method, in which 1.5 μl of protein solution (20 mg/mL) was mixed with 1.5 μl of reservoir solution containing 2 M ammonium sulfate, 0.1 M sodium cacodylate (pH 6.5), and 0.2 M sodium chloride. 0.1 M Sodium citrate tribasic dihydrate was used as an additive along with seeding to improve anisotropy and mosaicity. XusB crystals appeared after 7 days at 20 °C. The crystals were soaked into a cryoprotective solution corresponding to the reservoir of their crystallization drop with 25 % glycerol, followed by vitrification in liquid nitrogen.

The X-ray diffraction data were collected at the beamline 21-ID-G (0.97857 Å), at the Argonne National Laboratory’s Advanced Photon Source. XusBΔN crystal displayed a P3^1^21 space group with six molecules in the asymmetric unit. These crystals diffracted to a maximal resolution of 2.5 Å. The data were processed with autoPROC^96^. The XusBΔN structure was determined by molecular replacement using the coordinates obtained from AlphaFold2^45^ using the program PHASER^97^. Models were manually refined with COOT^98^, then further refinement was carried out using Phenix Refine^99^. The final refinement statistics are given in **Supplementary Table 1**. Atomic coordinates and structure factors have been deposited in the Protein Data Bank under the accession number 8GEX. The quality of all structures was checked with MOLPROBITY^100^. There are 0.3% Ramachandran outliers in the structures and at least 96% of the residues are in the most favored region. All figures were generated using PYMOL^101^.

### Intrinsic tryptophan fluorescence

The intrinsic tryptophan fluorescence of XusBΔN was measured using a Jobin Yvon Fluoromax-3 (Horiba Scientific) with a 1 ml quartz cuvette. The excitation wavelength was fixed to 285 nm, and emission spectra were acquired between 300 and 500 nm with a 10 mm sample pathlength. The temperature was maintained constant at 25 °C using an external thermostatic water circulator. To measure XusBΔN-Ent interactions, XusBΔN at 0.3 µM, was titrated with 0.05, 0.1, 0.15, 0.2, 0.3, 0.5, 0.6, 0.8, 1.0, 1.2, 1.3, 1.4, and 1.5 µM Ent. Data from three independent experiments were analyzed using nonlinear regression with ‘One Site-Specific Binding’ model (Y = B_max_ * X/(K_D_ + X) where X is the ligand concentration, Y is the fluorescence intensity, B_max_ is the maximum specific binding and K_D_ is the equilibrium binding constant) in GraphPad Prism 8.0.2.

### Small-angle X-ray scattering (SAXS)

Data were collected at the Structurally Integrated. Biology for the Life Sciences (SIBYLS) beamline at Lawrence Berkeley National Laboratory (Berkeley, CA). 50 μL samples containing free-XusB (5 mg/mL) and XusB-Ent complex (2 mg/mL) were prepared in 20 mM Tris (pH 8), and 150 mM NaCl. The sample volume run through the size exclusion chromatography (SEC^28^) was 50 μl and the eluent was split between the SEC-MALS and SAXS channels. 10.0-s X-ray exposures were collected continuously during a ∼35 min elution. All experiments were performed at 20 °C and data was processed as described below. The SAXS frames recorded before the protein elution peak were used as buffer blanks to subtract from all other frames. The subtracted frames were monitored by the radius of gyration (*Rg*) and scattering intensity at q = 0 Å−1 (I(0)), derived using the Guinier approximation I(q) = I(0)e-q*Rg*2/3 with the limits q\**Rg* <1.5. I(0) and *Rg* values were compared for each collected SAXS curve across the entire elution peak. The elution peak was mapped by plotting the scattering intensity at q = 0 Å−1 (I(0)), relative to the recorded frame. Uniform *Rg* values across an elution peak represent a homogenous assembly. The merged experimental SAXS data were additionally investigated for aggregation by inspecting Guinier plots. The SAXS data were processed using BioXTAS Raw ^102^. The *ab initio* electron maps obtained from the SAXS data of XusB and XusB-Ent complex were generated using DENSS ^103^. The SAXS data and models have been deposited in the SASDB databank under accession codes SASDRG2 (XusB) and SASDRH2 (Fe-Ent XusB complex). The final PDB model aligned with electron density map was visualized using ChimeraX.

### Ligand docking

Docking calculations were performed with two different starting conformations: (1) Ent coordinates extracted from the X-ray crystal structure of its complex with *Pseudomonas aeruginosa receptor* PfeA (PDBID 6R1F); (2) this structure after refinement by semi-empirical quantum mechanical calculations in Gaussian^104^ 16. The optimization used the PBE0-D3BJ method with a basis set def2-SVP. The two Fe-Ent structures were then docked to the X-ray crystal structure of XusBΔN. All docking calculations were performed in AutoDock Vina using the AutoDock server^47, 105^. Each calculation generated 20 candidate structures and the most favorable binding pose was chosen based on the Autodock Vina score. Both models were used to analyze the properties of the complex and the interactions of Fe-ent with XusB.

### Calculation of electrostatic fields

XusBΔN PDB files were converted to PQR format using the PDB2PQR server^106^. The surface electrostatic profile was then analyzed using the adaptive Poisson-Boltzmann solver ^107^ and viewed in PyMOL^101^.

### Quantification and Statistical analysis

Unless noted otherwise, data analysis was performed in GraphPad Prism v8.1.1. Values of bacterial population sizes, competitive indices, fold changes in mRNA levels, and normalized colon length were normally distributed after transformation by the natural logarithm. A two-tailed Student’s *t*-test was used for ln-transformed data. Unless otherwise stated, *, *P* < 0.05; **, *P* < 0.01; ***, *P* < 0.001; ns, not statistically significant. In all mouse experiments, *N* refers to the number of animals from which samples were taken. Sample sizes (i.e. the number of animals per group) were not estimated *a priori* since effect sizes in our system cannot be predicted. No predicted statistical outliers were removed since the presence or absence of these potential statistical outliers did not affect the overall interpretation. Mice that were euthanized early due to health concerns were excluded from analysis.

## Acknowledgments

We thank Dr. Melissa Ellermann (University of South Carolina) and Dr. Regis Stentz (Quadram Institute Bioscience) for their insightful discussions and comments on the concepts and methodology of the work. R. T. F was supported by NIH (T32AI112541). Work in W. Z.’s lab was funded by the NIH (1R35GM147470 and 1R01DK134692). M. J. M was supported by NIH (T32ES007028). Work in E. P. S lab was funded by the NIH (R01AI164587). N. G. S. was supported by Howard Hughes Medical Institute through the James H. Gilliam Fellowships for Advanced Study program (GT15104). T. T. was supported by National Science Foundation (NSF-GRFP No. 1937963). Work in M. X. B. lab was funded by the NIH (1R01DK131104 and 1R01AI168302). Any opinions, findings, conclusions, or recommendations expressed in this material are those of the author(s) and do not necessarily reflect the views of the funding agencies. The funders had no role in study design, data collection, and interpretation, or the decision to submit the work for publication.

## Author contributions

L.S., R. T. F., Y. R. P., W. J. C., and W. Z. designed the study. L.S. and R. T. F. performed and analyzed all the *in vitro* and *in vivo* experiments with help from H. D., and K. R.. Y. R. P. obtained the crystal and performed the structural analyses. X. R. and Z. J. Y. performed ligand docking calculations. L. S., N. G. S., T. P. S., and M. X. B. contributed to the gnotobiotic animal experiments. M. J. M and E. P. S contributed to metal measurements. A. L. performed proteomic experiments and analysis. N. P. and E. C. M. provided key reagents. L.S., R. T. F., Y. R. P., W. J. C., and W. Z. interpreted the data, W. J. C., and W. Z. wrote the manuscript, and all authors commented on the manuscript.

---

**Supplementary Figure 1: XusB coordinates enterobactin via hydrogen bonds and hydrophobic interactions. Related to Fig. 2**

(**A**) *B. thetaiotaomicron* cells expressing XusB LES mutant (XusB_lesMT_) were fractionated into outer membrane vesicles (P, OMV), cell-free supernatant (S, OMV), membrane, and cytoplasm/periplasm fractions, and probed for XusB and RpoB. RpoB is a cytoplasmic control. (**B**) Comparison of the experimental (blue) vs. theoretical (red) scattering curve of XusBΔN. (**C**) Small-angle X-ray scattering (SAXS) profiles of apo-XusB or Fe-Ent by DENSS with no symmetry constraints. Normalized pair distance distribution functions (P(r)) were calculated from the scattering profiles for XusB (red) and XusB+Ent complex (black). The P(r) curves for both constructures are consistent with globular protein shapes. (**D**) *B. thetaiotaomicron* cells expressing enterobactin-binding XusB mutant (XusB_entMT_) were fractionated into outer membrane vesicles (P, OMV), cell-free supernatant (S, OMV), membrane, and cytoplasm/periplasm fractions, and probed for XusB and RpoB. RpoB is a cytoplasmic control.

**Supplementary Figure 2: *Bacteroidetes* members encode homologous *xusABC* systems. Related to Fig. 2**

(**A**) A Blastp query of XusB was performed in a custom database consistent with 1,500 human metagenome-assembled genomes. The resulting homologs were aligned using MUSCLE, and the phylogenetic tree (**A**) was constructed using the neighbor-joining method. (**B**) Genetic synteny of the *xusABC* operon in *Bacteroidetes* that encode *xusB* homologs.

**Supplementary Figure 3: XusB homologs are structurally distinct from other surface exposed ligand binding proteins and harbor a positively charged calyx. Related to Fig. 2**

(**A**) Quantification of structural similarity between XusB homologs and other bacterial surface exposed ligand-binding proteins. The root-mean-square deviation (RMSD) of each structure to XusB was measured using PyMOL. SPB: Siderophore binding proteins in Gram-positive bacteria. VitB12: Vitamin B12. (**B**) Overall surface electrostatic profiles of XusB homologs, calculated using Adaptive Poisson-Boltzmann Solver (ABPS). The color scale represents electrostatic potentials in units of kT/e ranging from −5 (red, negatively charged) to +5 (blue, positively charged).

**Supplementary Figure 4: Enterobactin binding is required for XusB-mediated siderophore cross-feeding. Related to Fig. 2 & 3**

(**A**) Supernatant from donor *B. thetaiotaomicron* strains grown in iron-limited medium (BPS-supplemented SDM medium) was filter-sterilized, loaded with Fe-Ent, ultracentrifuged to collect OMV fraction, washed to remove unbound ligand, and introduced to recipient cells in BPS-supplemented SDM medium. Bacterial growth was measured by OD_600_. Bars represent the geometric mean. ***, *P* < 0.001; n.s., not statistically significant.

**Supplementary Figure 5: Comparison of inflammatory markers in the infectious colitis model. Related to Fig. 4**

(**A**) Groups of C57BL/6 mice were treated with a cocktail of antibiotics, followed by intragastrical inoculation of an equal mixture of the *B. thetaiotaomicron* wild-type strain and an isogenic mutant (Δ*xusB*), or a strain expressing the enterobactin-binding deficient mutant (Δ*xusB xusB*_entMT_). Mice were then either mock-treated (*N* = 6), intragastrically challenged with *S*. Tm SL1344 (WT vs. Δ*xusB*, *N* = 4; WT vs. Δ*xusB xusB*_entMT_, *N* = 5), or an isogenic *entB* mutant (WT vs. Δ*xusB*, *N* = 6). Four days after infection, the cecal tissue was collected, RNA extracted, and the transcript levels of indicated genes were measured using RT-qPCR (**A**). (**B**) Groups of C57BL/6 mice were treated with a cocktail of antibiotics, followed by intragastrical inoculation of either the *B. thetaiotaomicron* wild type strain or an isogenic Δ*xusB* mutant. Mice were challenged with the *S*. Tm SL1344 wild-type strain (*B. thetaiotaomicron* wild-type*, N* = 13; *B. thetaiotaomicron* Δ*xusB, N =* 14) or an isogenic *entB* mutant (*B. thetaiotaomicron* wild-type*, N* = 8; *B. thetaiotaomicron* Δ*xusB, N =* 7) for 4 days. The transcript levels of indicated genes in cecal tissue were measured using RT-qPCR (**B**). Bars represent the geometric mean. *, *P* < 0.05; ***; *P* < 0.001; ns, not statistically significant.

**Supplementary Figure 6: XusB homologs may mediate interbacterial competition or cross-feeding for xenosiderophores. Related to Fig. 5**

(**A**) XusB-containing culture supernatant was introduced into 10^8^ CFU of indicated recipient cells in SDM medium in the presence or absence of protease inhibitors. The culture supernatant recovered after 24 hours of anaerobic bacterial growth was resolved on SDS-PAGE gel and probed for XusB. (**B**) Supernatant from donor *B. vulgatus* strains grown in iron-limited medium (BPS-supplemented SDM medium) was filter-sterilized, loaded with Fe-Ent, washed to remove unbound ligand, and introduced to recipient cells in BPS-supplemented SDM medium. Bacterial growth was measured by OD_600_. (**C**) Supernatant from indicated donor strains grown in iron-limited medium (BPS supplemented SDM medium) was filter-sterilized, loaded with Fe-Ent, washed to remove unbound ligand, and introduced to *S*. Tm *entB* cells in BPS-supplemented SDM medium. Bacterial growth was measured by OD_600_. BV: *Bacteroides vulgatus*; BV CL: *Bacteroides vulgatus* CL09T03C04; BS: *Bacteroides salyersiae* DSM 18765; BTH: *Bacteroides thetaiotaomicron*. Bars represent the geometric mean. ***, *P* < 0.001; n.s., not statistically significant.

**Supplementary Figure 7: Comparison of inflammatory markers in gnotobiotic and the infectious colitis model. Related to Fig. 7**

(**A**) Groups of C57BL/6 mice were treated with a cocktail of antibiotics, followed by intragastrical inoculation of either the *B. thetaiotaomicron* wild-type strain (*N* = 10) or an isogenic Δ*xusB* mutant (*N* = 12). Mice were inoculated with an equal mixture of the *S*. Tm SL1344 wild-type strain and the isogenic Δ*fepA iroN* mutant strain. 4 days later, cecal tissue was collected, RNA extracted, and the transcript levels of indicated genes were measured using RT-qPCR. (**B**) Groups of C57BL/6 mice were treated with a cocktail of antibiotics, followed by intragastrical inoculation of a Δ*xusB* mutant strain. Mice were then inoculated with an equal mixture of the *S*. Tm SL1344 wild-type strain and the isogenic Δ*fepA iroN* mutant strain, followed by intragastrical administration of OMVs derived from the *B. thetaiotaomicron* wild-type strain (*N* = 11) or the isogenic Δ*xusB* mutant strain (*N* = 10) daily. mRNA levels of indicated genes in the cecal tissue were determined using RT-qPCR. **(C)** Groups of gnotobiotic Swiss Webster mice were monoassociated with either the *B. thetaiotaomicron* wild-type strain (*N* = 12) or the isogenic Δ*xusB* mutant (*N* = 11) for 7 days. Mice were then challenged with an equal mixture of the *S*. Tm SL1344 wild type strain and the isogenic Δ*fepA iroN* mutant strain. The cecal tissue was collected 3 days post-infection and the mRNA levels of indicated genes were determined using RT-qPCR. Bars represent the geometric mean. *, *P* < 0.05; n.s., not statically significant.

**Supplementary Figure 8: *B. thetaiotaomicron* abundance in an infectious colitis model. Related to Fig. 7**

(**A**) Groups of C57BL/6 mice were treated with a cocktail of antibiotics, followed by intragastrical inoculation of a Δ*xusB* mutant strain. Mice were then inoculated with an equal mixture of the *S*. Tm SL1344 wild-type strain and the isogenic Δ*fepA iroN* mutant strain, followed by intragastrical administration of an equal amount of OMVs derived from the *B. thetaiotaomicron* wild-type strain (*N* = 11) or the isogenic Δ*xusB* mutant strain (*N* = 10) daily. The abundance of *B. thetaiotaomicron* in cecal contents was determined 4 days after infection by plating on selective agar. Bars represent the geometric mean. n.s., not statistically significant.

## KEY RESOURCES TABLE

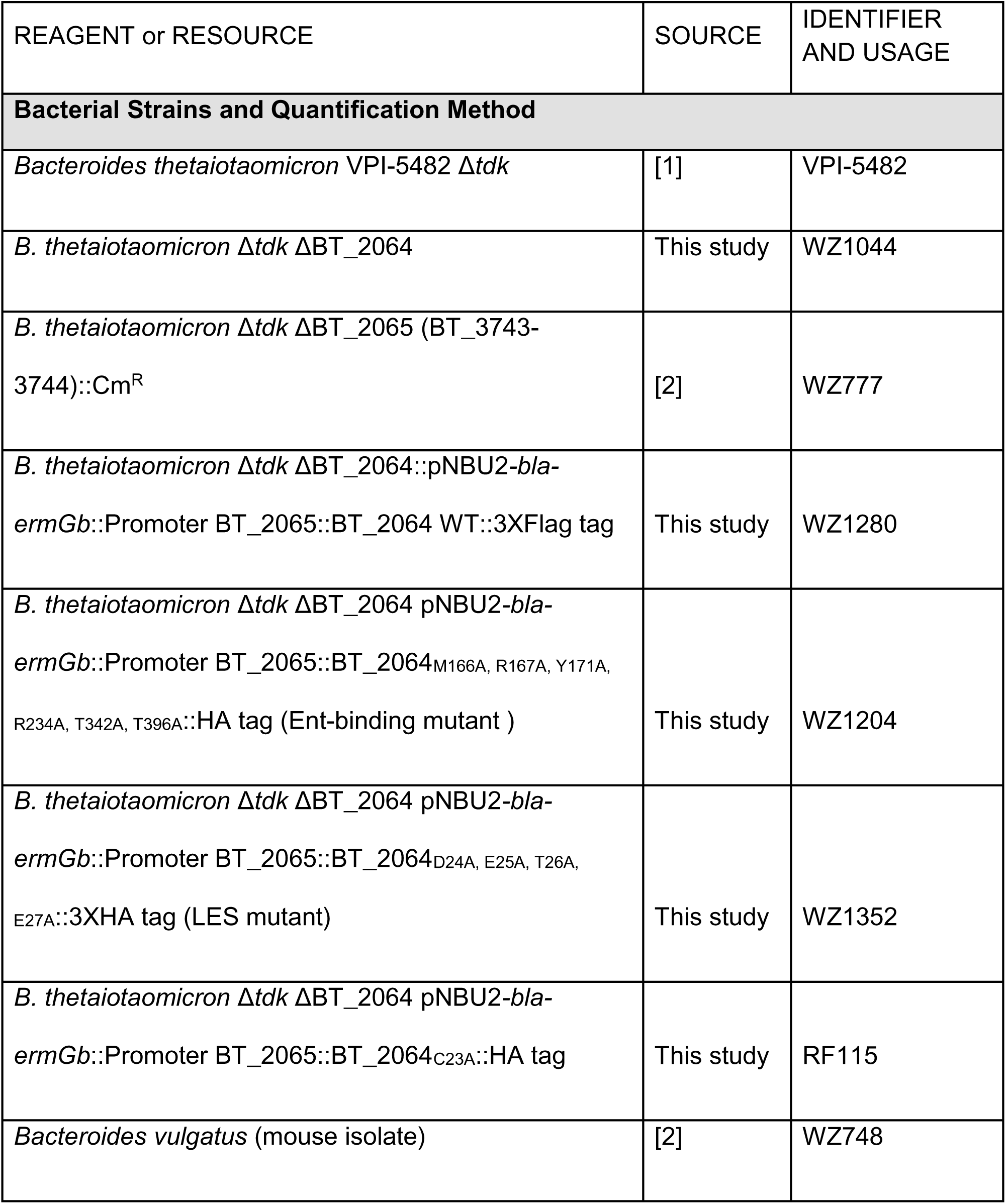
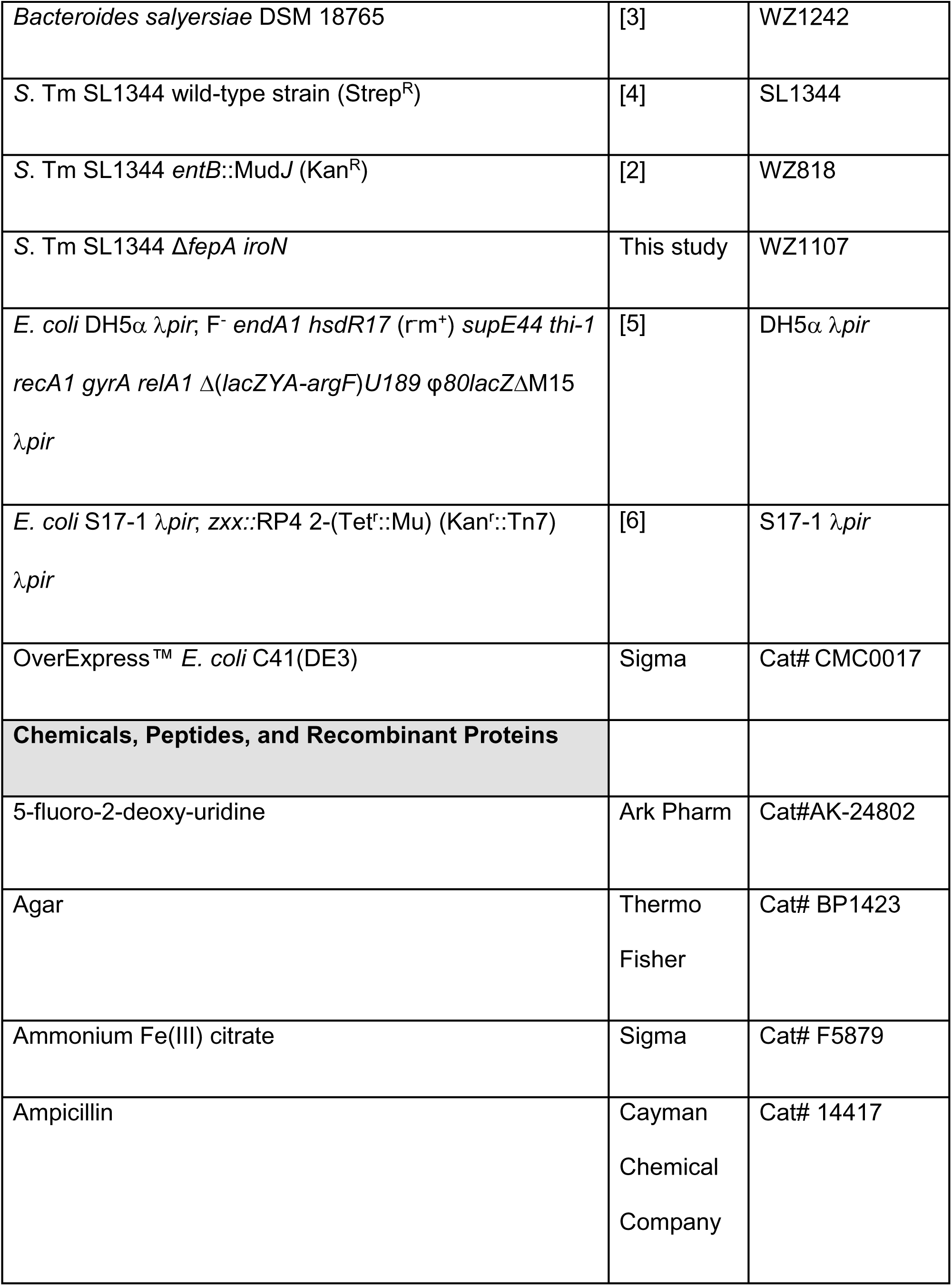
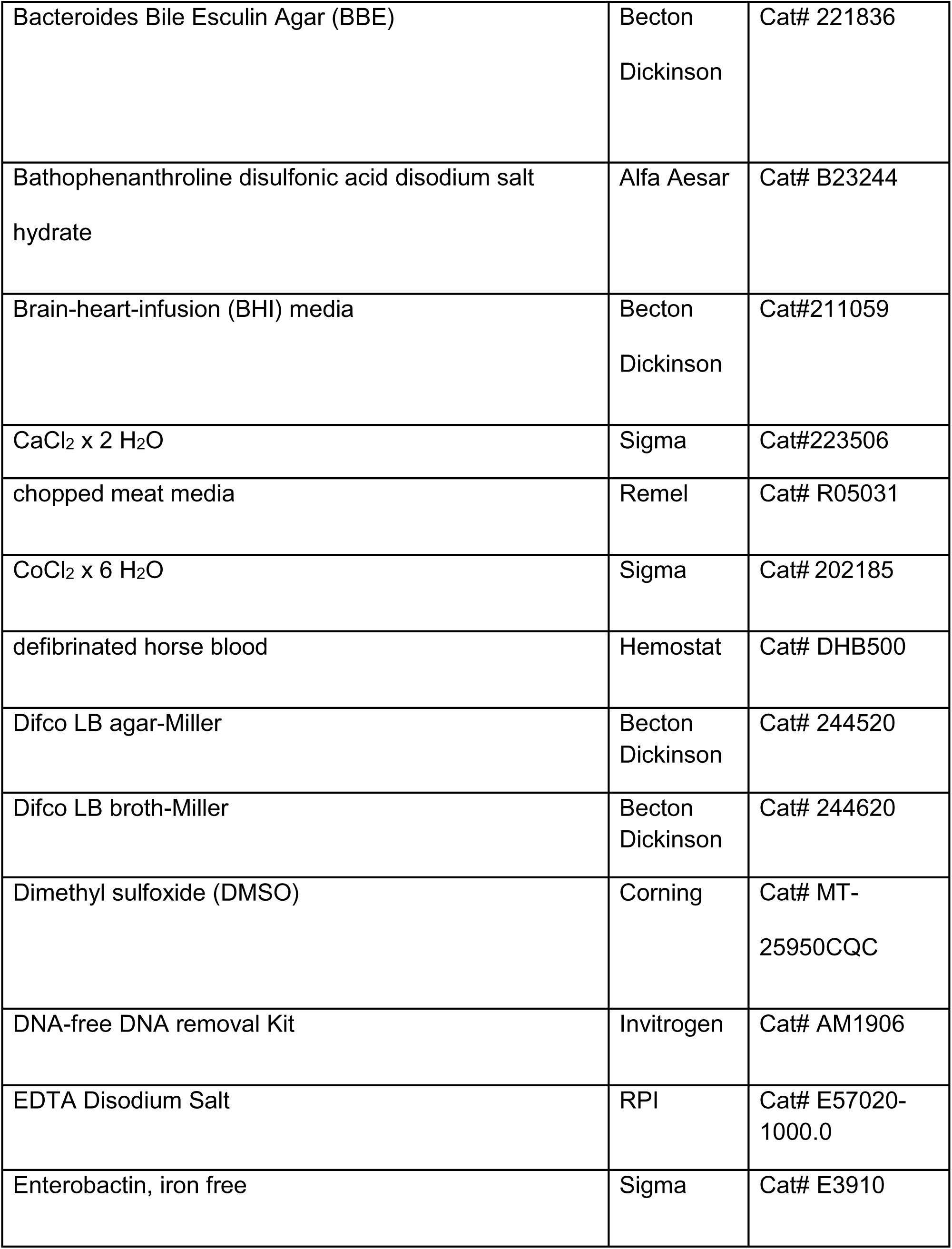
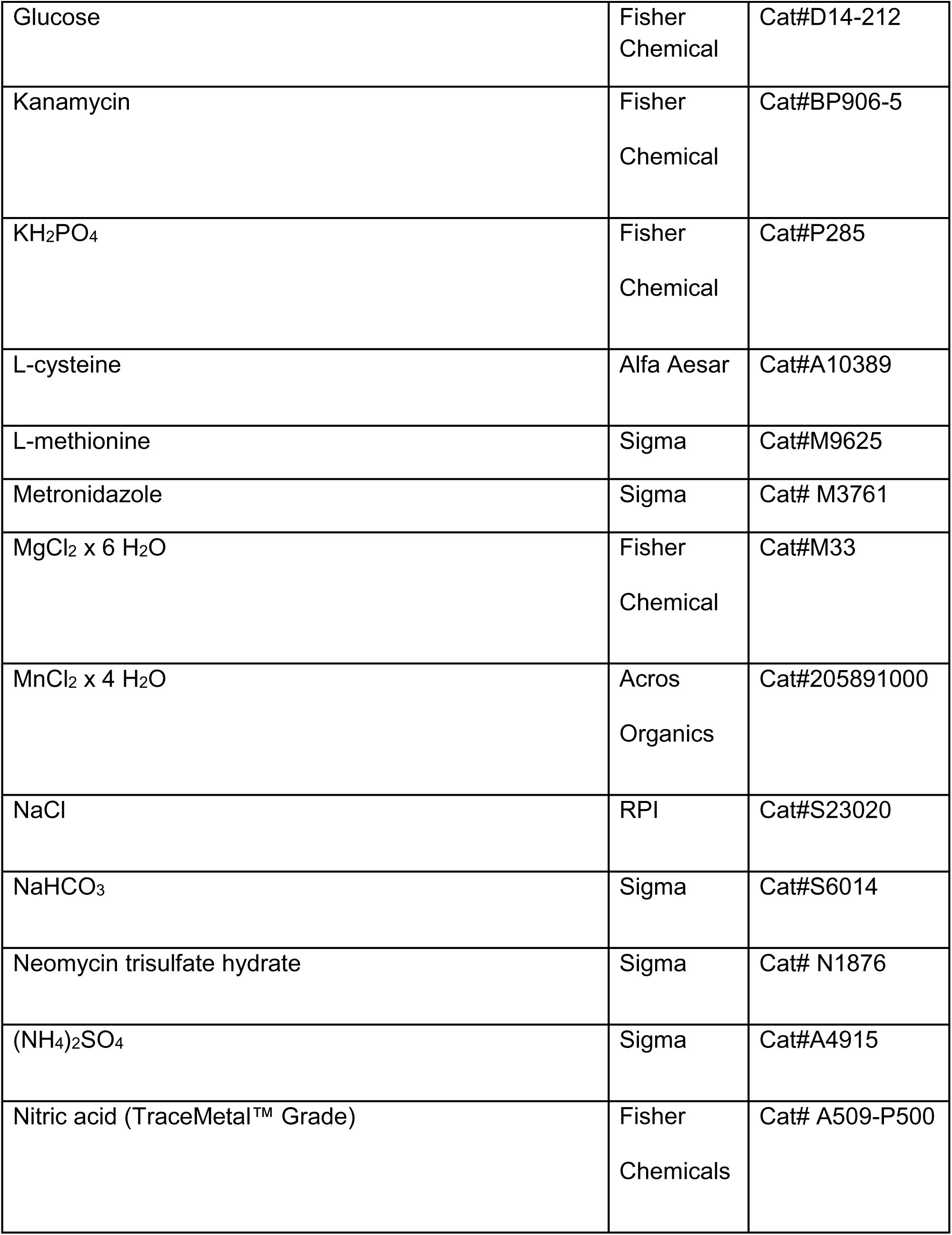
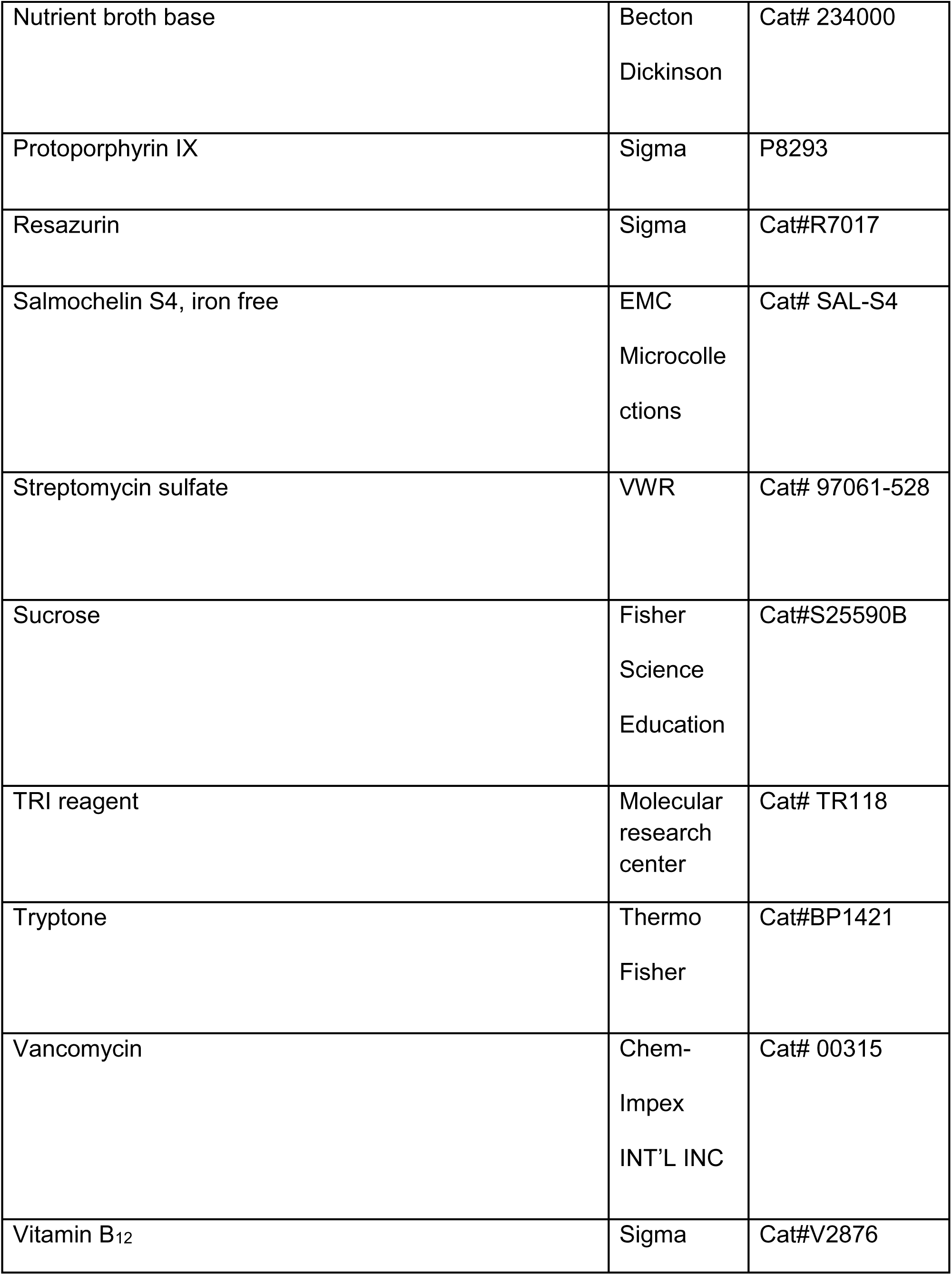
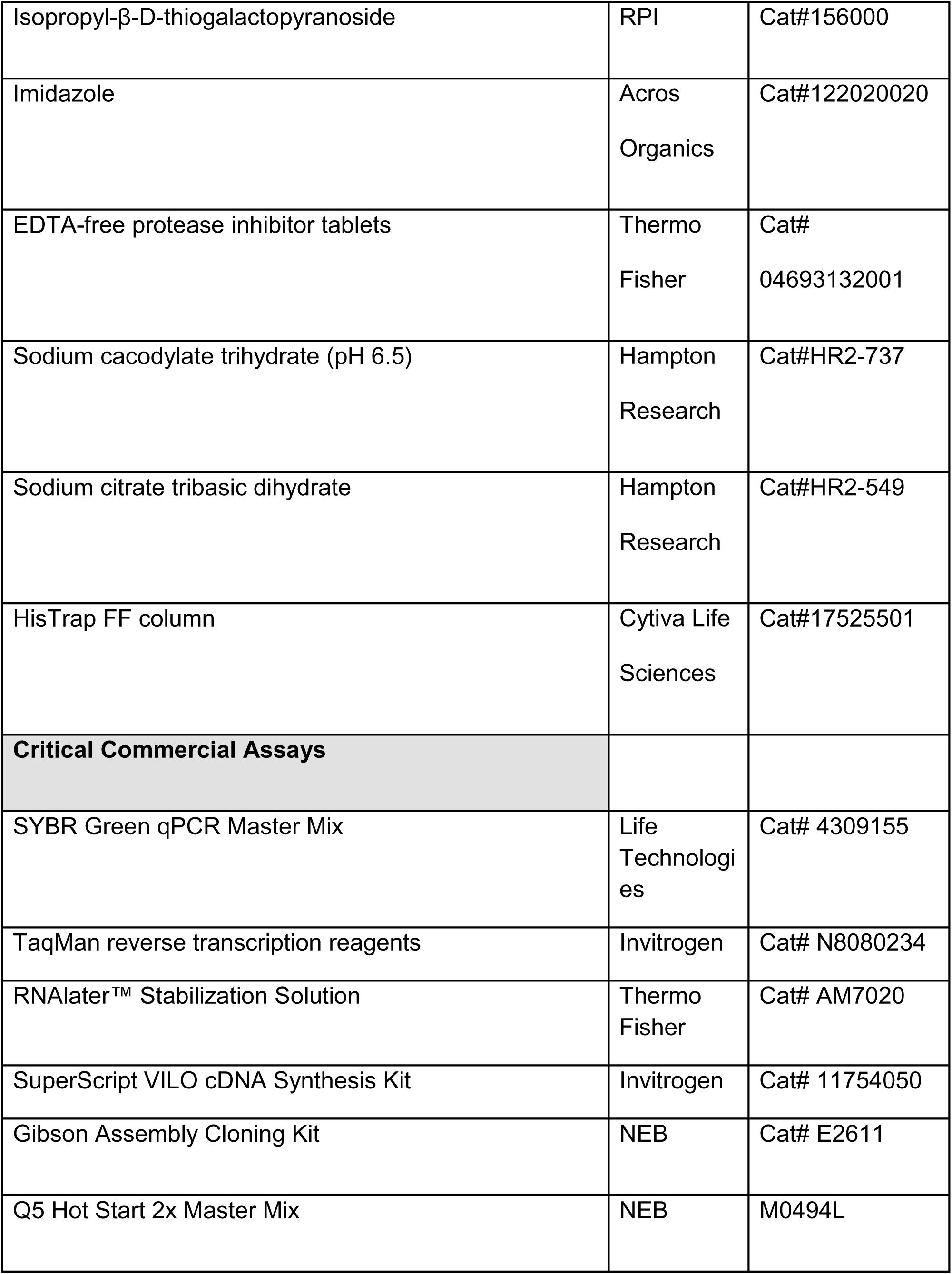
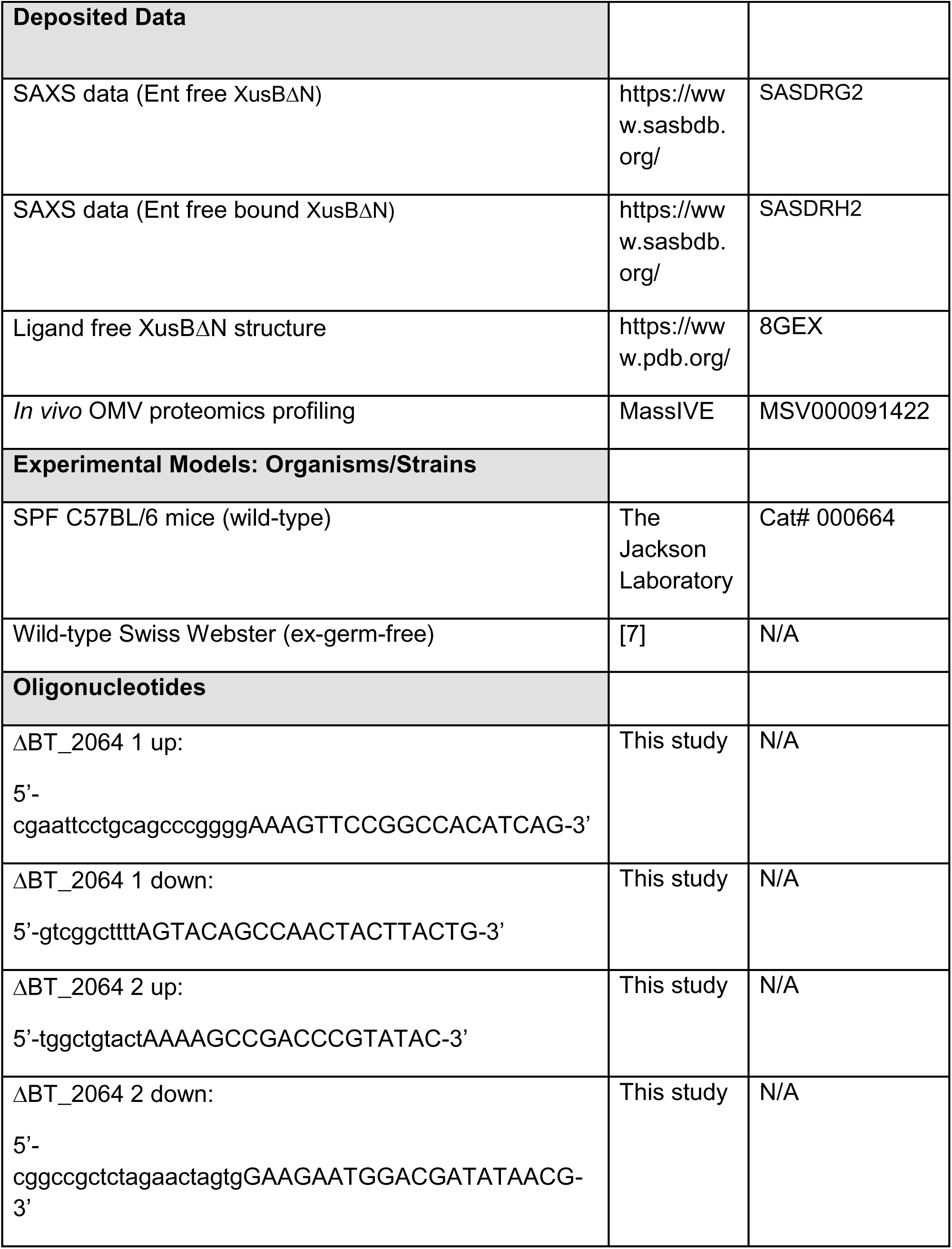
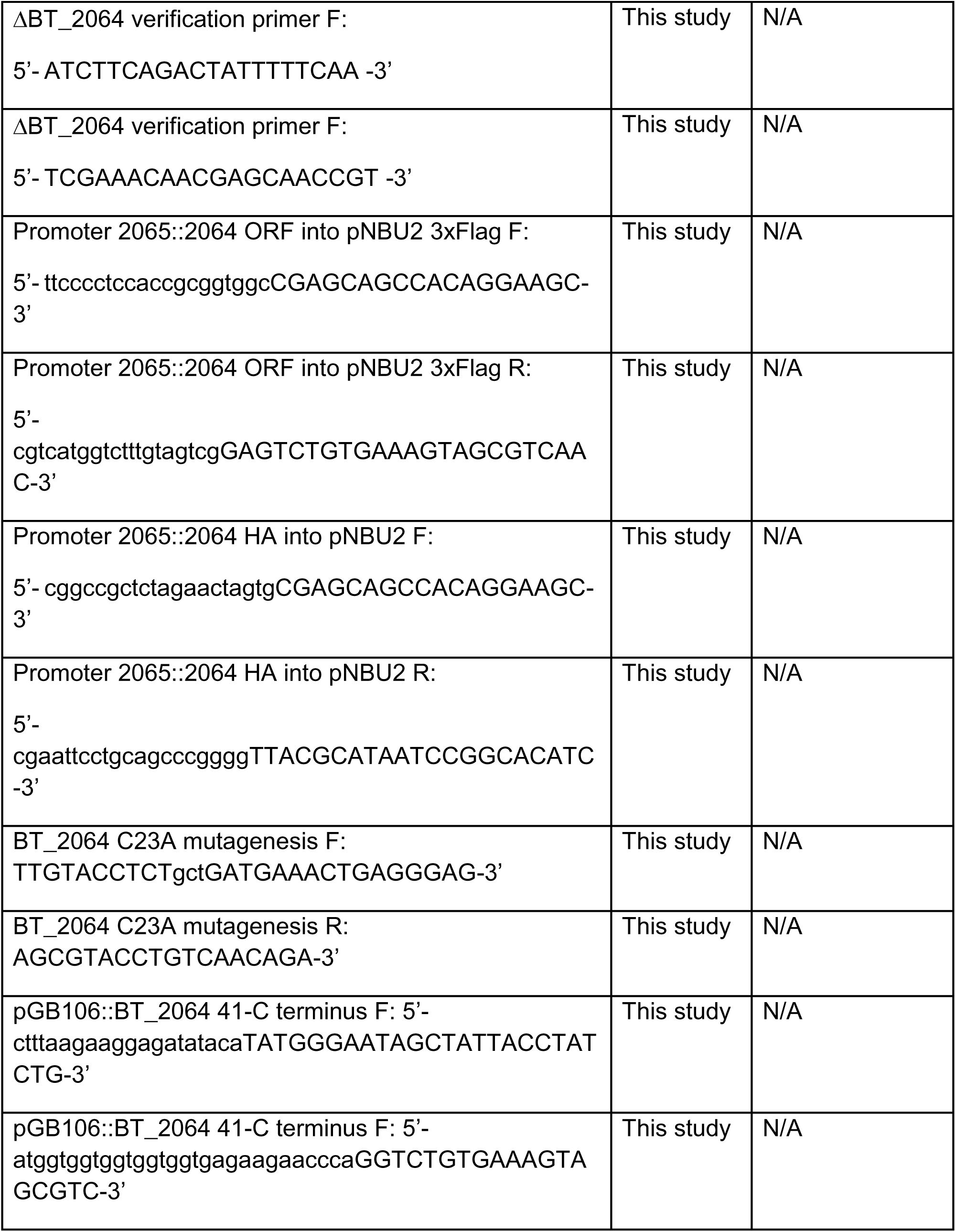
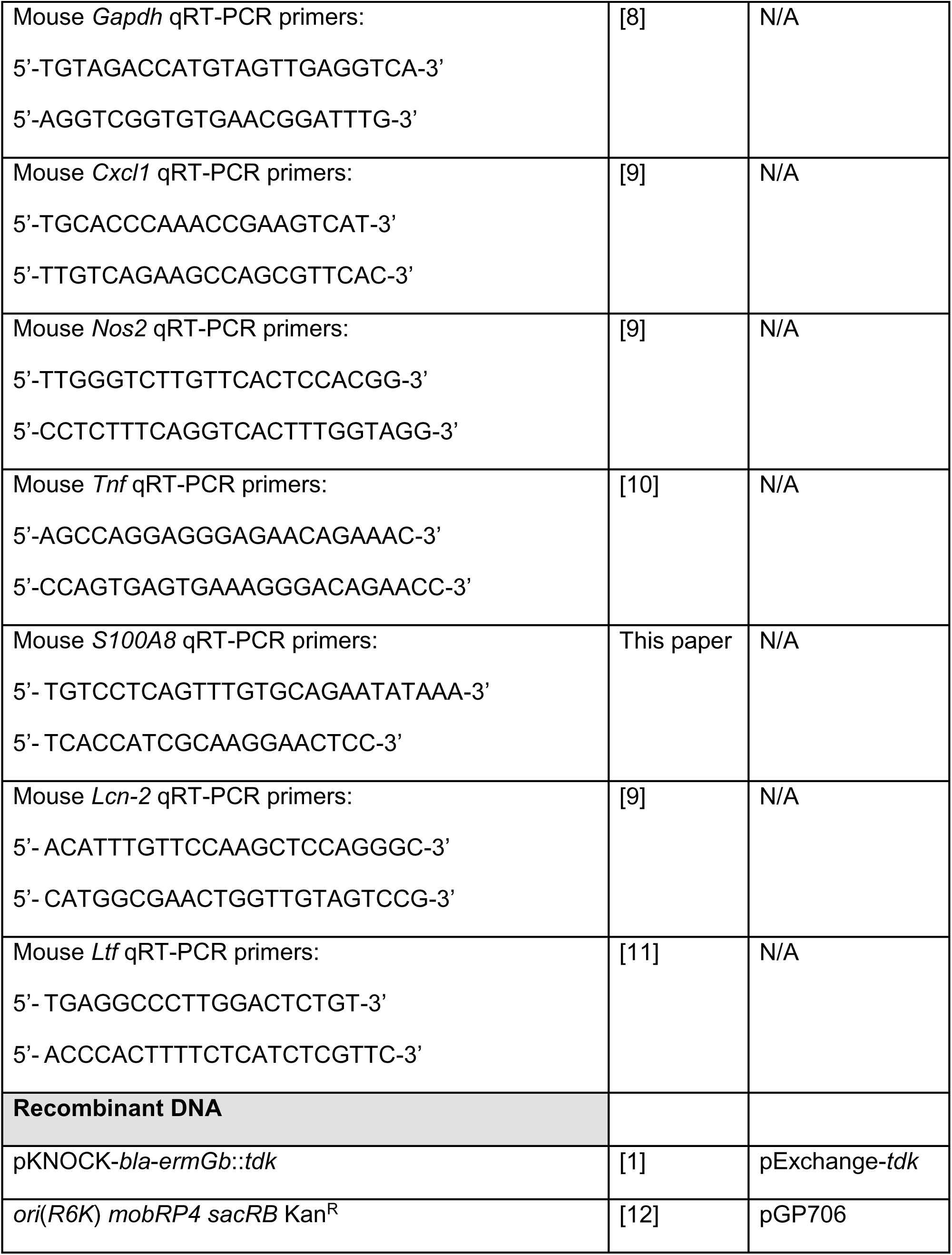
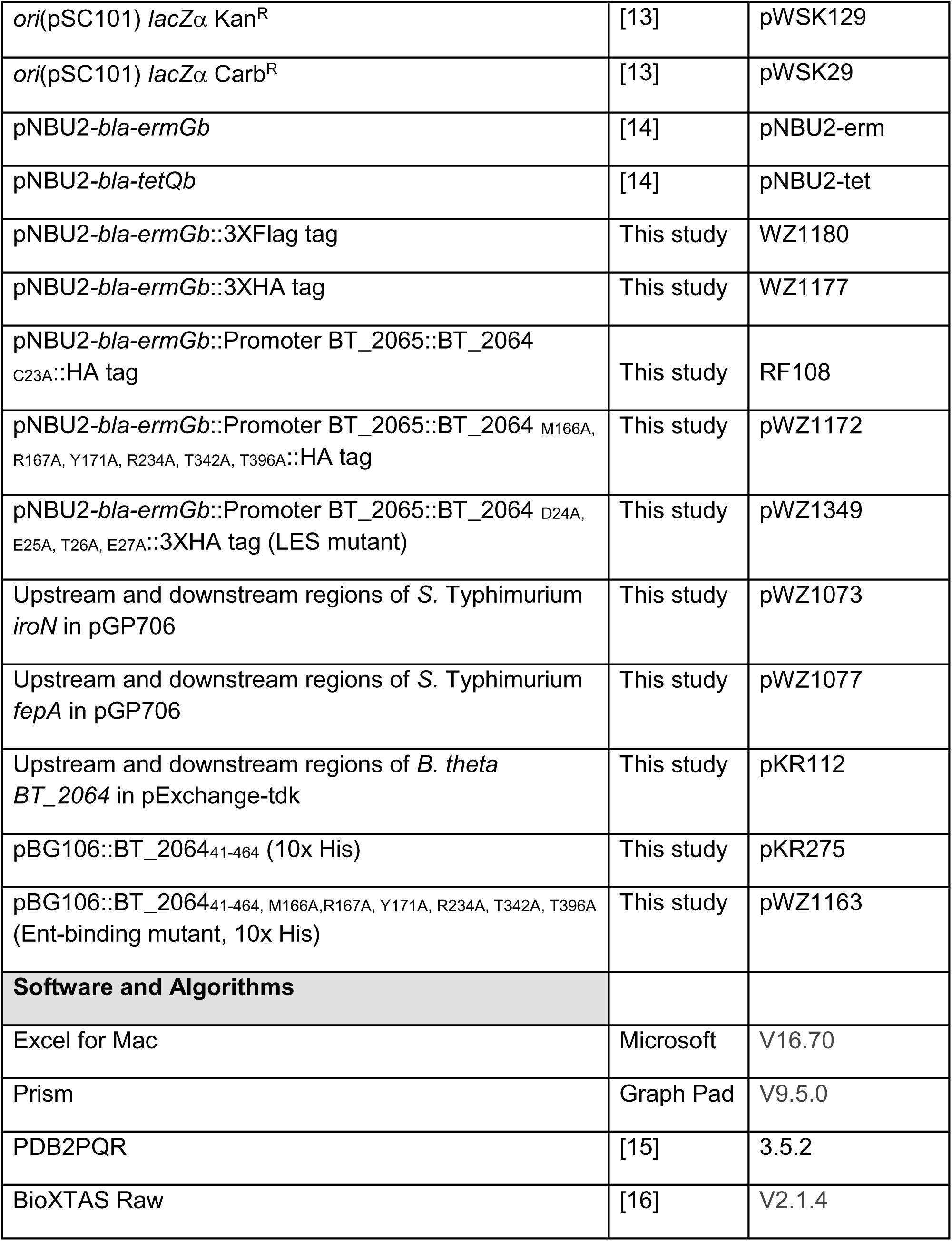
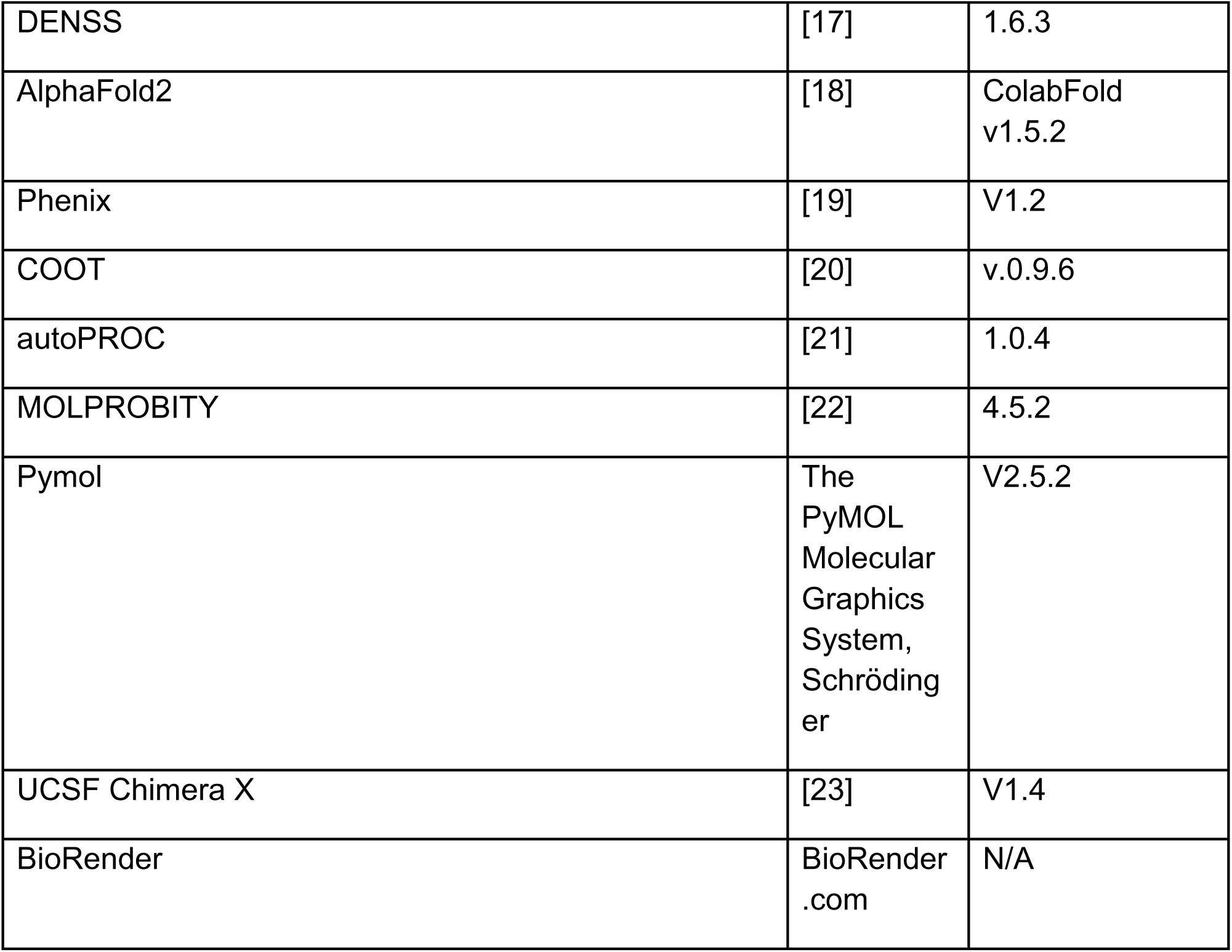

---

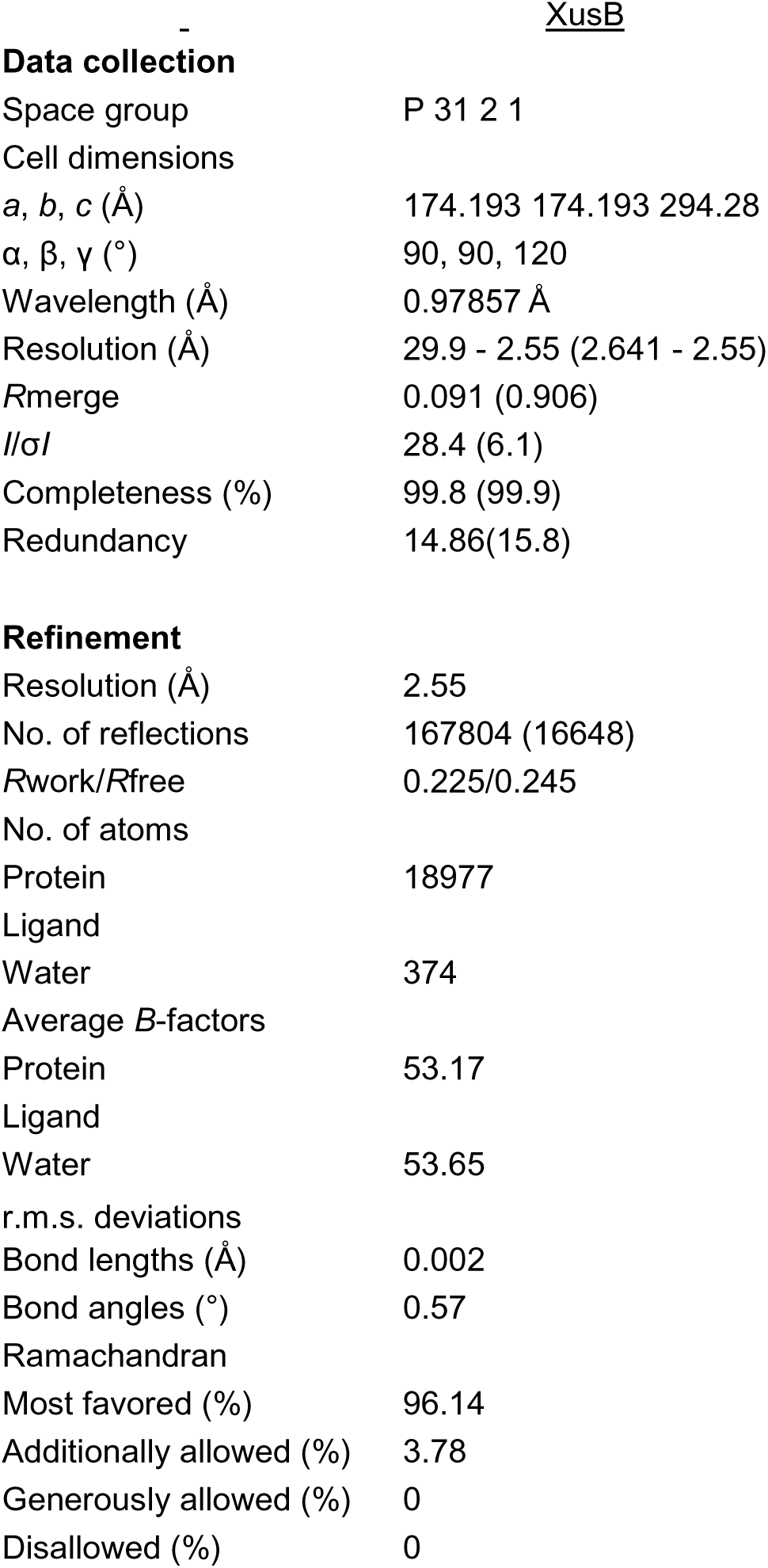

|  |  |
| --- | --- |
|  | <u>XusB</u> |
| **Data collection** |  |
| Space group | P 31 2 1 |
| Cell dimensions |  |
| *a*, *b*, *c* (Å) | 174.193 174.193 294.28 |
| $\alpha$, $\beta$, $\gamma$ (°) | 90, 90, 120 |
| Wavelength (Å) | 0.97857 Å |
| Resolution (Å) | 29.9 - 2.55 (2.641 - 2.55) |
| *R*<sub>merge</sub> | 0.091 (0.906) |
| *I*/σ*I* | 28.4 (6.1) |
| Completeness (%) | 99.8 (99.9) |
| Redundancy | 14.86(15.8) |
| **Refinement** |  |
| Resolution (Å) | 2.55 |
| No. of reflections | 167804 (16648) |
| *R*<sub>work</sub>/*R*<sub>free</sub> | 0.225/0.245 |
| No. of atoms |  |
| Protein | 18977 |
| Ligand |  |
| Water | 374 |
| Average *B*-factors |  |
| Protein | 53.17 |
| Ligand |  |
| Water | 53.65 |
| r.m.s. deviations |  |
| Bond lengths (Å) | 0.002 |
| Bond angles (°) | 0.57 |
| Ramachandran |  |
| Most favored (%) | 96.14 |
| Additionally allowed (%) | 3.78 |
| Generously allowed (%) | 0 |
| Disallowed (%) | 0 |

## Reference

1. Rocha, E.R. and C.J. Smith, Transcriptional regulation of the Bacteroides fragilis ferritin gene (ftnA) by redox stress. Microbiology, 2004. 150(Pt 7): p. 2125–34. doi:10.1099/mic.0.26948-0

2. Zhu, W., et al., Xenosiderophore Utilization Promotes Bacteroides thetaiotaomicron Resilience during Colitis. Cell Host Microbe, 2020. 27(3): p. 376–388 e8. doi:10.1016/j.chom.2020.01.010

3. Wang, R.F. and S.R. Kushner, Construction of versatile low-copy-number vectors for cloning, sequencing and gene expression in Escherichia coli. Gene, 1991. 100: p. 195–9.

1. Almeida, A., et al., A new genomic blueprint of the human gut microbiota. Nature, 2019. 568(7753): p. 499–504.

2. Vemuri, R., et al., Beyond Just Bacteria: Functional Biomes in the Gut Ecosystem Including Virome, Mycobiome, Archaeome and Helminths. Microorganisms, 2020. 8(4).

3. Sommer, F., et al., The resilience of the intestinal microbiota influences health and disease. Nat Rev Microbiol, 2017. 15(10): p. 630–638.

4. Scheffer, M., et al., Catastrophic shifts in ecosystems. Nature, 2001. 413(6856): p. 591–6.

5. Ingrisch, J. and M. Bahn, Towards a Comparable Quantification of Resilience. Trends Ecol Evol, 2018. 33(4): p. 251–259.

6. Winter, S.E. and A.J. Baumler, Dysbiosis in the inflamed intestine: chance favors the prepared microbe. Gut Microbes, 2014. 5(1): p. 71–3.

7. Lozupone, C.A., et al., Diversity, stability and resilience of the human gut microbiota. Nature, 2012. 489(7415): p. 220–30.

8. Lloyd-Price, J., et al., Multi-omics of the gut microbial ecosystem in inflammatory bowel diseases. Nature, 2019. 569(7758): p. 655–662.

9. Miller, B.M., et al., The longitudinal and cross-sectional heterogeneity of the intestinal microbiota. Curr Opin Microbiol, 2021. 63: p. 221–230.

10. Lee, J.Y., R.M. Tsolis, and A.J. Baumler, The microbiome and gut homeostasis. Science, 2022. 377(6601): p. eabp9960.

11. Murdoch, C.C. and E.P. Skaar, Nutritional immunity: the battle for nutrient metals at the host-pathogen interface. Nat Rev Microbiol, 2022. 20(11): p. 657–670.

12. Murdoch, C.C. and E.P. Skaar, Nutritional immunity: the battle for nutrient metals at the host-pathogen interface. Nat Rev Microbiol, 2022.

13. Cassat, J.E. and E.P. Skaar, Iron in infection and immunity. Cell Host Microbe, 2013. 13(5): p. 509–519.

14. Raymond, K.N., E.A. Dertz, and S.S. Kim, Enterobactin: an archetype for microbial iron transport. Proc Natl Acad Sci U S A, 2003. 100(7): p. 3584–8.

15. Goetz, D.H., et al., The neutrophil lipocalin NGAL is a bacteriostatic agent that interferes with siderophore-mediated iron acquisition. Mol Cell, 2002. 10(5): p. 1033–43.

16. Baumler, A.J., et al., IroN, a novel outer membrane siderophore receptor characteristic of Salmonella enterica. J Bacteriol, 1998. 180(6): p. 1446–53.

17. Baumler, A.J., et al., Identification of a new iron regulated locus of Salmonella typhi. Gene, 1996. 183(1-2): p. 207–13.

18. Bister, B., et al., The structure of salmochelins: C-glucosylated enterobactins of Salmonella enterica. Biometals, 2004. 17(4): p. 471–81.

19. Fischbach, M.A., et al., The pathogen-associated iroA gene cluster mediates bacterial evasion of lipocalin 2. Proc Natl Acad Sci U S A, 2006. 103(44): p. 16502–7.

20. Raffatellu, M., et al., Lipocalin-2 resistance confers an advantage to Salmonella enterica serotype Typhimurium for growth and survival in the inflamed intestine. Cell Host Microbe, 2009. 5(5): p. 476–86.

21. Behnsen, J., et al., The cytokine IL-22 promotes pathogen colonization by suppressing related commensal bacteria. Immunity, 2014. 40(2): p. 262–73.

22. Deriu, E., et al., Probiotic bacteria reduce salmonella typhimurium intestinal colonization by competing for iron. Cell Host Microbe, 2013. 14(1): p. 26–37.

23. Costa, L.F., et al., Iron acquisition pathways and colonization of the inflamed intestine by Salmonella enterica serovar Typhimurium. Int J Med Microbiol, 2016. 306(8): p. 604–610.

24. Bratburd, J.R., et al., Gut Microbial and Metabolic Responses to Salmonella enterica Serovar Typhimurium and Candida albicans. mBio, 2018. 9(6).

25. Jacobson, A., et al., A Gut Commensal-Produced Metabolite Mediates Colonization Resistance to Salmonella Infection. Cell Host Microbe, 2018. 24(2): p. 296–307 e7.

26. Gillis, C.C., et al., Dysbiosis-Associated Change in Host Metabolism Generates Lactate to Support Salmonella Growth. Cell Host Microbe, 2018. 23(1): p. 54–64 e6.

27. Sana, T.G., et al., Salmonella Typhimurium utilizes a T6SS-mediated antibacterial weapon to establish in the host gut. Proc Natl Acad Sci U S A, 2016. 113(34): p. E5044–51.

28. Zhu, W., et al., Xenosiderophore Utilization Promotes Bacteroides thetaiotaomicron Resilience during Colitis. Cell Host Microbe, 2020. 27(3): p. 376–388 e8.

29. Jautzus, T., J. van Gestel, and A.T. Kovacs, Complex extracellular biology drives surface competition during colony expansion in Bacillus subtilis. ISME J, 2022. 16(10): p. 2320–2328.

30. Bomberger, J.M., et al., Long-distance delivery of bacterial virulence factors by Pseudomonas aeruginosa outer membrane vesicles. PLoS Pathog, 2009. 5(4): p. e1000382.

31. Wang, T., et al., Type VI Secretion System Transports Zn2+ to Combat Multiple Stresses and Host Immunity. PLoS Pathog, 2015. 11(7): p. e1005020.

32. Endicott, N.P., et al., Emergence of Ferrichelatase Activity in a Siderophore-Binding Protein Supports an Iron Shuttle in Bacteria. ACS Cent Sci, 2020. 6(4): p. 493–506.

33. Elhenawy, W., M.O. Debelyy, and M.F. Feldman, Preferential packing of acidic glycosidases and proteases into Bacteroides outer membrane vesicles. mBio, 2014. 5(2): p. e00909–14.

34. McMillan, H.M. and M.J. Kuehn, The extracellular vesicle generation paradox: a bacterial point of view. EMBO J, 2021. 40(21): p. e108174.

35. Buchanan, S.K., et al., Crystal structure of the outer membrane active transporter FepA from Escherichia coli. Nat Struct Biol, 1999. 6(1): p. 56–63.

36. Josts, I., et al., Structural insights into a novel family of integral membrane siderophore reductases. Proc Natl Acad Sci U S A, 2021. 118(34).

37. von Heijne, G., The structure of signal peptides from bacterial lipoproteins. Protein Eng, 1989. 2(7): p. 531–4.

38. Lauber, F., G.R. Cornelis, and F. Renzi, Identification of a New Lipoprotein Export Signal in Gram-Negative Bacteria. mBio, 2016. 7(5).

39. Schwechheimer, C. and M.J. Kuehn, Outer-membrane vesicles from Gram-negative bacteria: biogenesis and functions. Nat Rev Microbiol, 2015. 13(10): p. 605–19.

40. Valguarnera, E., et al., Surface Exposure and Packing of Lipoproteins into Outer Membrane Vesicles Are Coupled Processes in Bacteroides. mSphere, 2018. 3(6).

41. Stentz, R., et al., The Proteome of Extracellular Vesicles Produced by the Human Gut Bacteria Bacteroides thetaiotaomicron In Vivo Is Influenced by Environmental and Host Derived Factors. Appl Environ Microbiol, 2022. 88(16): p. e0053322.

42. Rocha, E.R. and A.S. Krykunivsky, Anaerobic utilization of Fe(III)-xenosiderophores among Bacteroides species and the distinct assimilation of Fe(III)-ferrichrome by Bacteroides fragilis within the genus. Microbiologyopen, 2017. 6(4).

43. Moynie, L., et al., The complex of ferric-enterobactin with its transporter from Pseudomonas aeruginosa suggests a two-site model. Nat Commun, 2019. 10(1): p. 3673.

44. Ghisaidoobe, A.B. and S.J. Chung, Intrinsic tryptophan fluorescence in the detection and analysis of proteins: a focus on Forster resonance energy transfer techniques. Int J Mol Sci, 2014. 15(12): p. 22518–38.

45. Jumper, J., et al., Highly accurate protein structure prediction with AlphaFold. Nature, 2021. 596(7873): p. 583–589.

46. Chen, C.K., N.L. Chan, and A.H. Wang, The many blades of the beta-propeller proteins: conserved but versatile. Trends Biochem Sci, 2011. 36(10): p. 553–61.

47. Morris, G.M., et al., AutoDock4 and AutoDockTools4: Automated docking with selective receptor flexibility. J Comput Chem, 2009. 30(16): p. 2785–91.

48. Zou, Y., et al., 1,520 reference genomes from cultivated human gut bacteria enable functional microbiome analyses. Nat Biotechnol, 2019. 37(2): p. 179–185.

49. Rocha, E.R. and C.J. Smith, Transcriptional regulation of the Bacteroides fragilis ferritin gene (ftnA) by redox stress. Microbiology (Reading), 2004. 150(Pt 7): p. 2125–2134.

50. Lee, S.M., et al., Bacterial colonization factors control specificity and stability of the gut microbiota. Nature, 2013. 501(7467): p. 426–9.

51. Curtis, M.M., et al., The gut commensal Bacteroides thetaiotaomicron exacerbates enteric infection through modification of the metabolic landscape. Cell Host Microbe, 2014. 16(6): p. 759–69.

52. Hantke, K., et al., Salmochelins, siderophores of Salmonella enterica and uropathogenic Escherichia coli strains, are recognized by the outer membrane receptor IroN. Proc Natl Acad Sci U S A, 2003. 100(7): p. 3677–82.

53. Sassone-Corsi, M., et al., Microcins mediate competition among Enterobacteriaceae in the inflamed gut. Nature, 2016. 540(7632): p. 280–283.

54. Rakoff-Nahoum, S., K.R. Foster, and L.E. Comstock, The evolution of cooperation within the gut microbiota. Nature, 2016. 533(7602): p. 255–9.

55. Leventhal, G.E., M. Ackermann, and K.T. Schiessl, Why microbes secrete molecules to modify their environment: the case of iron-chelating siderophores. J R Soc Interface, 2019. 16(150): p. 20180674.

56. Kramer, J., O. Ozkaya, and R. Kummerli, Bacterial siderophores in community and host interactions. Nat Rev Microbiol, 2020. 18(3): p. 152–163.

57. Young, I.G., et al., Biosynthesis of the iron-transport compound enterochelin: mutants of Escherichia coli unable to synthesize 2,3-dihydroxybenzoate. J Bacteriol, 1971. 106(1): p. 51–7.

58. Luke, R.K. and F. Gibson, Location of three genes concerned with the conversion of 2,3-dihydroxybenzoate into enterochelin in Escherichia coli K-12. J Bacteriol, 1971. 107(2): p. 557–62.

59. Baba, T., et al., Construction of Escherichia coli K-12 in-frame, single-gene knockout mutants: the Keio collection. Mol Syst Biol, 2006. 2: p. 2006 0008.

60. McIntosh, M.A., S.S. Chenault, and C.F. Earhart, Genetic and physiological studies on the relationship between colicin B resistance and ferrienterochelin uptake in Escherichia coli K-12. J Bacteriol, 1979. 137(1): p. 653–7.

61. Rogers, A.W.L., R.M. Tsolis, and A.J. Baumler, Salmonella versus the Microbiome. Microbiol Mol Biol Rev, 2021. 85(1).

62. Raffatellu, M. and A.J. Baumler, Salmonella’s iron armor for battling the host and its microbiota. Gut Microbes, 2010. 1(1): p. 70–72.

63. Peppas, N.A., P.J. Hansen, and P.A. Buri, A theory of molecular diffusion in the intestinal mucus. International Journal of Pharmaceutics, 1984. 20(1): p. 107–118.

64. Stecher, B., et al., Comparison of Salmonella enterica serovar Typhimurium colitis in germfree mice and mice pretreated with streptomycin. Infect Immun, 2005. 73(6): p. 3228–41.

65. Ng, K.M., et al., Microbiota-liberated host sugars facilitate post-antibiotic expansion of enteric pathogens. Nature, 2013. 502(7469): p. 96–9.

66. Shelton, C.D., et al., Salmonella enterica serovar Typhimurium uses anaerobic respiration to overcome propionate-mediated colonization resistance. Cell Rep, 2022. 38(1): p. 110180.

67. Bronner, D.N., et al., Genetic Ablation of Butyrate Utilization Attenuates Gastrointestinal Salmonella Disease. Cell Host Microbe, 2018. 23(2): p. 266–273 e4.

68. Deschemin, J.C., et al., The microbiota shifts the iron sensing of intestinal cells. FASEB J, 2016. 30(1): p. 252–61.

69. Reddy, B.S., J.R. Pleasants, and B.S. Wostmann, Effect of intestinal microflora on iron and zinc metabolism, and on activities of metalloenzymes in rats. J Nutr, 1972. 102(1): p. 101–7.

70. Das, N.K., et al., Microbial Metabolite Signaling Is Required for Systemic Iron Homeostasis. Cell Metab, 2020. 31(1): p. 115–130 e6.

71. Song, X., et al., Microbial bile acid metabolites modulate gut RORgamma(+) regulatory T cell homeostasis. Nature, 2020. 577(7790): p. 410–415.

72. Arpaia, N., et al., Metabolites produced by commensal bacteria promote peripheral regulatory T-cell generation. Nature, 2013. 504(7480): p. 451–5.

73. Sequeira, R.P., et al., Commensal Bacteroidetes protect against Klebsiella pneumoniae colonization and transmission through IL-36 signalling. Nat Microbiol, 2020. 5(2): p. 304–313.

74. Desai, M.S., et al., A Dietary Fiber-Deprived Gut Microbiota Degrades the Colonic Mucus Barrier and Enhances Pathogen Susceptibility. Cell, 2016. 167(5): p. 1339–1353 e21.

75. Zhu, W., et al., Precision editing of the gut microbiota ameliorates colitis. Nature, 2018. 553(7687): p. 208–211.

76. Zhu, W., et al., Editing of the gut microbiota reduces carcinogenesis in mouse models of colitis-associated colorectal cancer. J Exp Med, 2019. 216(10): p. 2378–2393.

77. Federici, S., et al., Targeted suppression of human IBD-associated gut microbiota commensals by phage consortia for treatment of intestinal inflammation. Cell, 2022. 185(16): p. 2879–2898 e24.

78. Sinha, A., et al., Transplantation of bacteriophages from ulcerative colitis patients shifts the gut bacteriome and exacerbates the severity of DSS colitis. Microbiome, 2022. 10(1): p. 105.

79. Falony, G., et al., Population-level analysis of gut microbiome variation. Science, 2016. 352(6285): p. 560–4.

80. Rivera-Chavez, F. and A.J. Baumler, The Pyromaniac Inside You: Salmonella Metabolism in the Host Gut. Annu Rev Microbiol, 2015. 69: p. 31–48.

81. Fletcher, J.R., et al., Clostridioides difficile exploits toxin-mediated inflammation to alter the host nutritional landscape and exclude competitors from the gut microbiota. Nat Commun, 2021. 12(1): p. 462.

82. Pruss, K.M. and J.L. Sonnenburg, C. difficile exploits a host metabolite produced during toxin-mediated disease. Nature, 2021. 593(7858): p. 261–265.

83. Lopez, L.R., et al., Microenvironmental Factors that Shape Bacterial Metabolites in Inflammatory Bowel Disease. Front Cell Infect Microbiol, 2022. 12: p. 934619.

84. Cullen, T.W., et al., Gut microbiota. Antimicrobial peptide resistance mediates resilience of prominent gut commensals during inflammation. Science, 2015. 347(6218): p. 170–5.

85. Wilson, B.R., et al., Siderophores in Iron Metabolism: From Mechanism to Therapy Potential. Trends Mol Med, 2016. 22(12): p. 1077–1090.

86. Krewulak, K.D. and H.J. Vogel, Structural biology of bacterial iron uptake. Biochim Biophys Acta, 2008. 1778(9): p. 1781–804.

87. Grigg, J.C., et al., The Staphylococcus aureus siderophore receptor HtsA undergoes localized conformational changes to enclose staphyloferrin A in an arginine-rich binding pocket. J Biol Chem, 2010. 285(15): p. 11162–71.

88. Wexler, A.G., et al., Human gut Bacteroides capture vitamin B(12) via cell surface exposed lipoproteins. Elife, 2018. 7.

89. Degnan, P.H., et al., Human gut microbes use multiple transporters to distinguish vitamin B(1)(2) analogs and compete in the gut. Cell Host Microbe, 2014. 15(1): p. 47–57.

90. Glenwright, A.J., et al., Structural basis for nutrient acquisition by dominant members of the human gut microbiota. Nature, 2017. 541(7637): p. 407–411.

91. Ernits, K., et al., First crystal structure of an endo-levanase - the BT1760 from a human gut commensal Bacteroides thetaiotaomicron. Sci Rep, 2019. 9(1): p. 8443.

92. Langelueddecke, C., et al., Expression and function of the lipocalin-2 (24p3/NGAL) receptor in rodent and human intestinal epithelia. PLoS One, 2013. 8(8): p. e71586.

93. Patrick, S., et al., A comparison of the haemagglutinating and enzymic activities of Bacteroides fragilis whole cells and outer membrane vesicles. Microb Pathog, 1996. 20(4): p. 191–202.

94. Rocha, E.R. and C.J. Smith, Transcriptional regulation of the Bacteroides fragilis ferritin gene (ftnA) by redox stress. Microbiology, 2004. 150(Pt 7): p. 2125–34.

95. Wang, R.F. and S.R. Kushner, Construction of versatile low-copy-number vectors for cloning, sequencing and gene expression in Escherichia coli. Gene, 1991. 100: p. 195–9.

96. Vonrhein, C., et al., Data processing and analysis with the autoPROC toolbox. Acta Crystallogr D Biol Crystallogr, 2011. 67(Pt 4): p. 293–302.

97. McCoy, A.J., et al., Phaser crystallographic software. J Appl Crystallogr, 2007. 40(Pt 4): p. 658–674.

98. Emsley, P., et al., Features and development of Coot. Acta Crystallogr D Biol Crystallogr, 2010. 66(Pt 4): p. 486–501.

99. Liebschner, D., et al., Macromolecular structure determination using X-rays, neutrons and electrons: recent developments in Phenix. Acta Crystallogr D Struct Biol, 2019. 75(Pt 10): p. 861–877.

100. Chen, V.B., et al., MolProbity: all-atom structure validation for macromolecular crystallography. Acta Crystallogr D Biol Crystallogr, 2010. 66(Pt 1): p. 12–21.

101. Schrodinger, LLC, The PyMOL Molecular Graphics System, Version 1.8. 2015.

102. Hopkins, J.B., R.E. Gillilan, and S. Skou, BioXTAS RAW: improvements to a free open source program for small-angle X-ray scattering data reduction and analysis. J Appl Crystallogr, 2017. 50(Pt 5): p. 1545–1553.

103. Grant, T.D., Ab initio electron density determination directly from solution scattering data. Nat Methods, 2018. 15(3): p. 191–193.

104. Frisch, M.J., et al., Gaussian 16 Rev. C.01. 2016: Wallingford, CT.

105. Eberhardt, J., et al., AutoDock Vina 1.2.0: New Docking Methods, Expanded Force Field, and Python Bindings. J Chem Inf Model, 2021. 61(8): p. 3891–3898.

106. Dolinsky, T.J., et al., PDB2PQR: an automated pipeline for the setup of Poisson Boltzmann electrostatics calculations. Nucleic Acids Res, 2004. 32(Web Server issue): p. W665–7.

107. Jurrus, E., et al., Improvements to the APBS biomolecular solvation software suite. Protein Sci, 2018. 27(1): p. 112–128.

## Key Resource Table Reference

1. Koropatkin, N.M., et al., Starch catabolism by a prominent human gut symbiont is directed by the recognition of amylose helices. Structure, 2008. 16(7): p. 1105–15.

2. Zhu, W., et al., Xenosiderophore Utilization Promotes Bacteroides thetaiotaomicron Resilience during Colitis. Cell Host Microbe, 2020. 27(3): p. 376–388 e8.

3. Song, Y.L., et al., “Bacteroides nordii„ sp. nov. and “Bacteroides salyersae„ sp. nov. isolated from clinical specimens of human intestinal origin. J Clin Microbiol, 2004. 42(12): p. 5565–70.

4. Hoiseth, S.K. and B.A. Stocker, Aromatic-dependent Salmonella typhimurium are non-virulent and effective as live vaccines. Nature, 1981. 291(5812): p. 238–9.

5. Pal, D., et al., Multipartite regulation of rctB, the replication initiator gene of Vibrio cholerae chromosome II. J Bacteriol, 2005. 187(21): p. 7167–75.

6. Simon, R., U. Priefer, and A. Puhler, A Broad Host Range Mobilization System for In Vivo Genetic Engineering: Transposon Mutagenesis in Gram Negative Bacteria. Nature Biotechnology, 1983. 1: p. 784–791.

7. Spiga, L., et al., An oxidative central metabolism enables Salmonella to utilize microbiota-derived succinate. Cell Host Microbe, 2017. In press.

8. Overbergh, L., et al., The use of real-time reverse transcriptase PCR for the quantification of cytokine gene expression. J Biomol Tech, 2003. 14(1): p. 33–43.

9. Godinez, I., et al., T cells help to amplify inflammatory responses induced by Salmonella enterica serotype Typhimurium in the intestinal mucosa. Infect Immun, 2008. 76(5): p. 2008–17.

10. Wilson, R.P., et al., The Vi-capsule prevents Toll-like receptor 4 recognition of Salmonella. Cell Microbiol, 2008. 10(4): p. 876–90.

11. Fullerton, P.T., Jr., et al., Follistatin is critical for mouse uterine receptivity and decidualization. Proc Natl Acad Sci U S A, 2017. 114(24): p. E4772–E4781.

12. Hughes, E.R., et al., Microbial Respiration and Formate Oxidation as Metabolic Signatures of Inflammation-Associated Dysbiosis. Cell Host Microbe, 2017. 21(2): p. 208–219.

13. Wang, R.F. and S.R. Kushner, Construction of versatile low-copy-number vectors for cloning, sequencing and gene expression in Escherichia coli. Gene, 1991. 100: p. 195–9.

14. Wang, J., et al., Characterization of a Bacteroides mobilizable transposon, NBU2, which carries a functional lincomycin resistance gene. J Bacteriol, 2000. 182(12): p. 3559–71.

15. Dolinsky, T.J., et al., PDB2PQR: an automated pipeline for the setup of Poisson-Boltzmann electrostatics calculations. Nucleic Acids Res, 2004. 32(Web Server issue): p. W665–7.

16. Hopkins, J.B., R.E. Gillilan, and S. Skou, BioXTAS RAW: improvements to a free open-source program for small-angle X-ray scattering data reduction and analysis. J Appl Crystallogr, 2017. 50(Pt 5): p. 1545–1553.

17. Grant, T.D., Ab initio electron density determination directly from solution scattering data. Nat Methods, 2018. 15(3): p. 191–193.

18. Jumper, J., et al., Highly accurate protein structure prediction with AlphaFold. Nature, 2021. 596(7873): p. 583–589.

19. Adams, P.D., et al., PHENIX: building new software for automated crystallographic structure determination. Acta Crystallogr D Biol Crystallogr, 2002. 58(Pt 11): p. 1948–54.

20. Ng, W.M., et al., Structure of trimeric pre-fusion rabies virus glycoprotein in complex with two protective antibodies. Cell Host Microbe, 2022. 30(9): p. 1219–1230 e7.

21. Vonrhein, C., et al., Data processing and analysis with the autoPROC toolbox. Acta Crystallogr D Biol Crystallogr, 2011. 67(Pt 4): p. 293–302.

22. Chen, V.B., et al., MolProbity: all-atom structure validation for macromolecular crystallography. Acta Crystallogr D Biol Crystallogr, 2010. 66(Pt 1): p. 12–21.

23. Pettersen, E.F., et al., UCSF ChimeraX: Structure visualization for researchers, educators, and developers. Protein Sci, 2021. 30(1): p. 70–82.

