## Supplementary material for "Iron acquisition by a commensal bacterium modifies host nutritional immunity during *Salmonella* infection": N/A

**Supplementary Fig. 1**

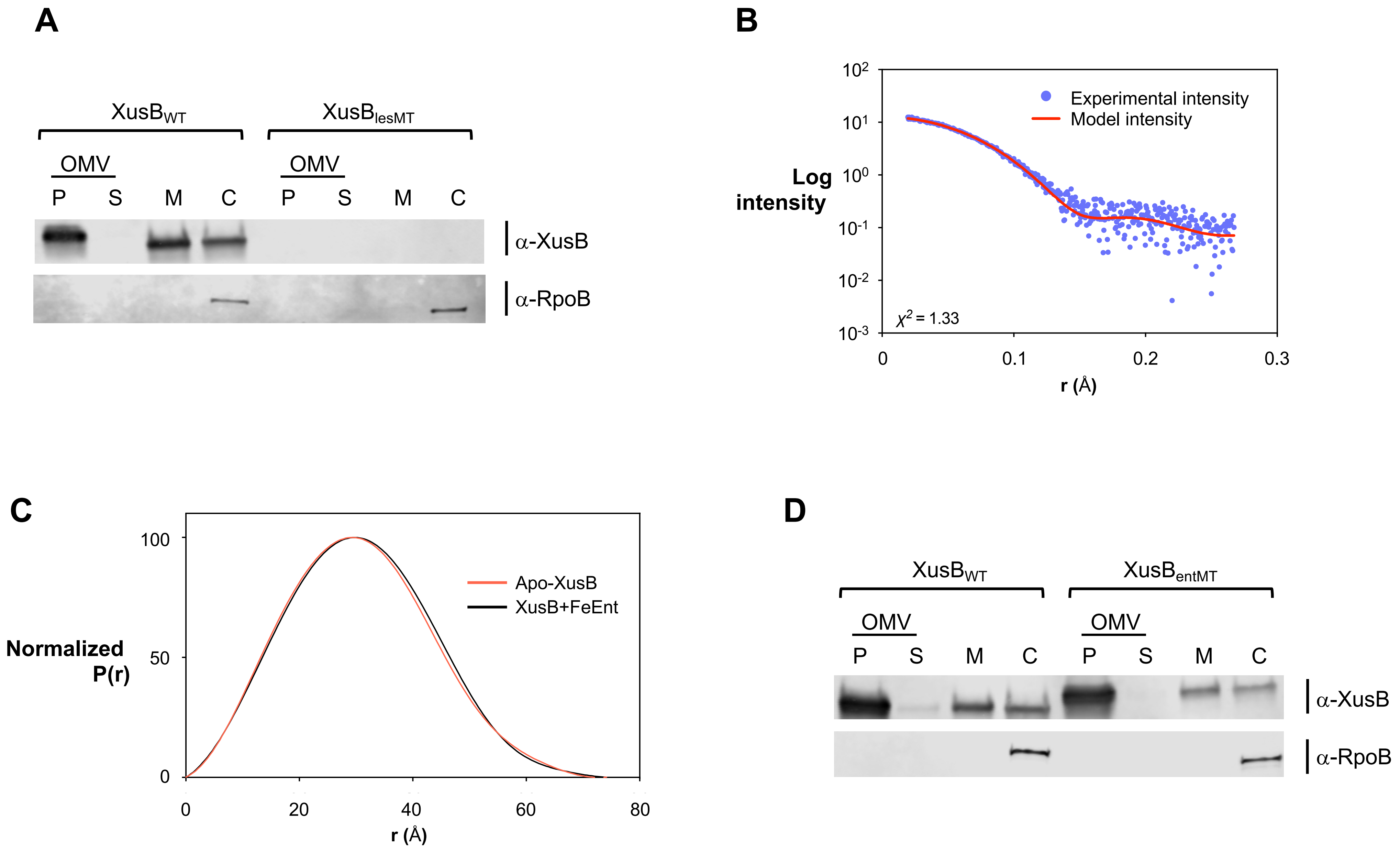

**Supplementary Figure 1: XusB coordinates enterobactin via hydrogen bonds and hydrophobic interactions**

(**A**) *B. thetaiotaomicron* cells expressing XusB LES mutant ($XusB_{lesMT}$) were fractionated into outer membrane vesicles (P, OMV), cell-free supernatant (S, OMV), membrane, and cytoplasm/periplasm fractions, and probed for XusB and RpoB. RpoB is a cytoplasmic control. (**B**) Comparison of the experimental (blue) vs. theoretical (red) scattering curve of XusBΔN. (**C**) Small-angle X-ray scattering (SAXS) profiles of apo-XusB or Fe-Ent by DENSS with no symmetry constraints. Normalized pair distance distribution functions (P(r)) were calculated from the scattering profiles for XusB (red) and XusB+Ent complex (black). The P(r) curves for both constructures are consistent with globular protein shapes. (**D**) *B. thetaiotaomicron* cells expressing enterobactin-binding XusB mutant ($XusB_{entMT}$) were fractionated into outer membrane vesicles (P, OMV), cell-free supernatant (S, OMV), membrane, and cytoplasm/periplasm fractions, and probed for XusB and RpoB. RpoB is a cytoplasmic control.

#### Supplementary Fig. 2 (Neighbor joining)

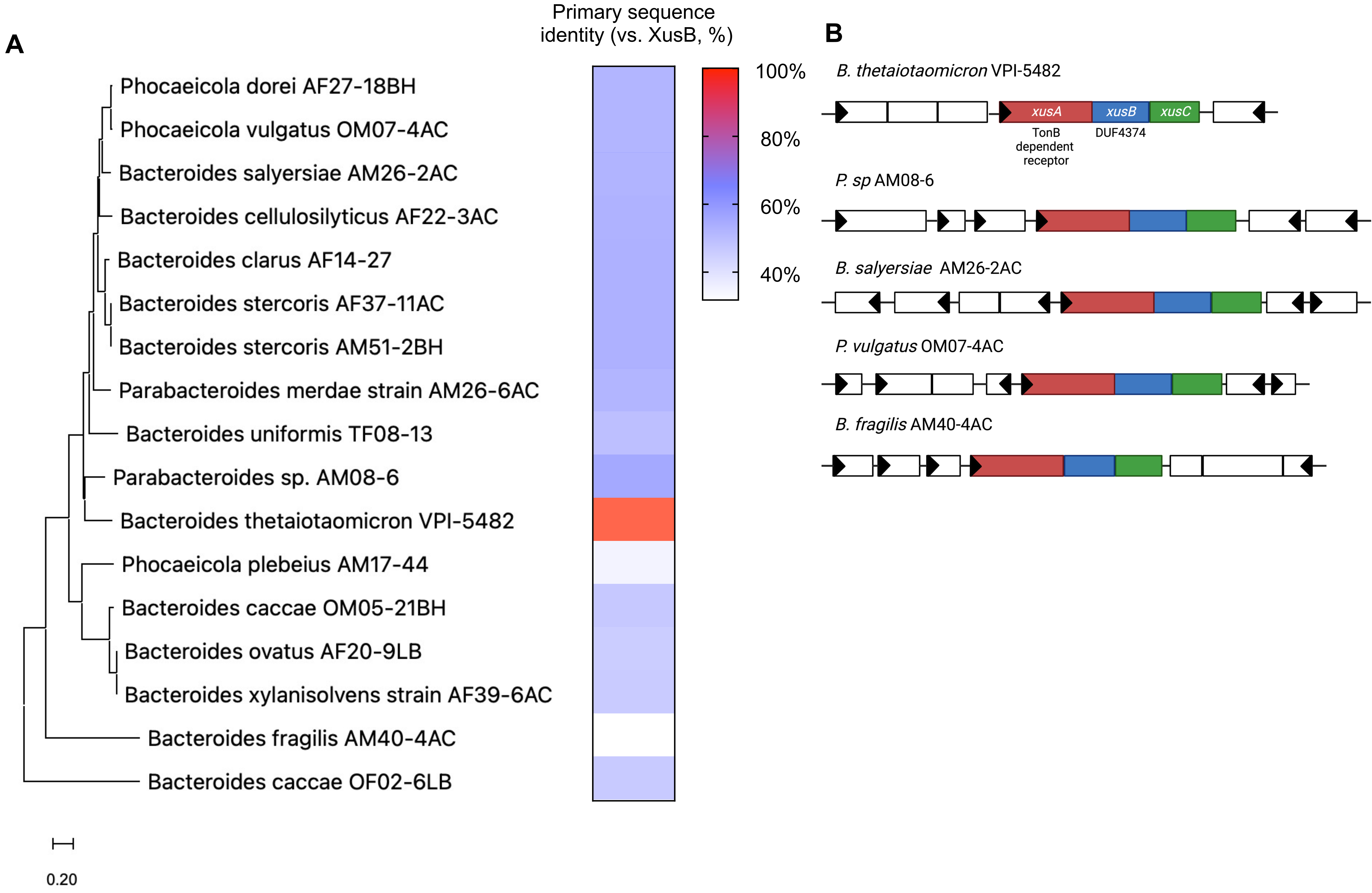

**Supplementary Figure 2: *Bacteroidetes* members encode homologous *xusABC* systems**

(**A**) A Blastp query of XusB was performed in a custom database consistent with 1,500 human metagenome-assembled genomes. The resulting homologs were aligned using MUSCLE, and the phylogenetic tree (**A**) was constructed using the neighbor-joining method. (**B**) Genetic synteny of the *xusABC* operon in *Bacteroidetes* that encode *xusB* homologs.

### Supplementary Fig. 3

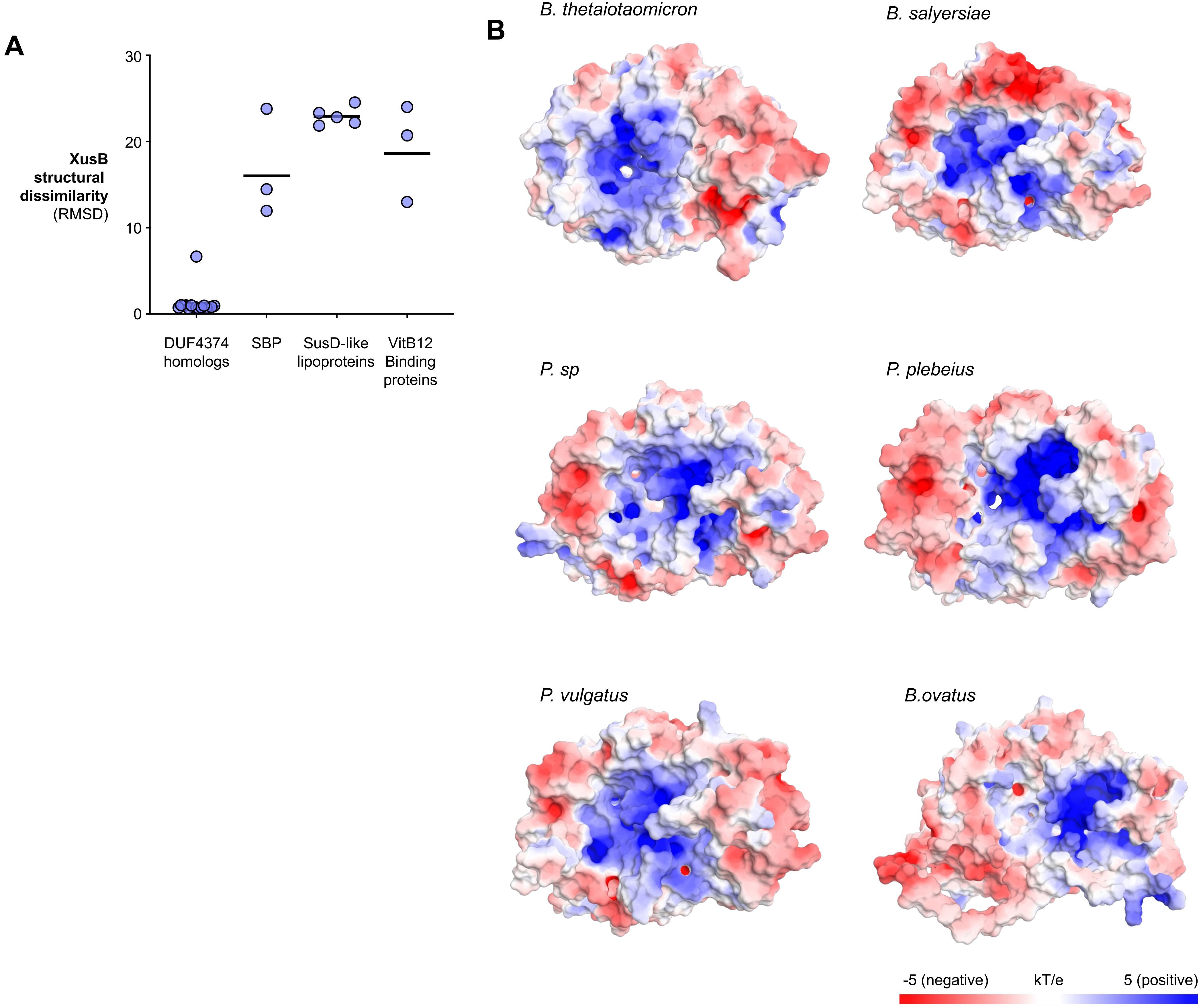

**Supplementary Figure 3: XusB homologs are structurally distinct from other surface exposed ligand binding proteins and harbor a positively charged calyx**

(**A**) Quantification of structural similarity between XusB homologs and other bacterial surface exposed ligand-binding proteins. The root-mean-square deviation (RMSD) of each structure to XusB was measured using PyMOL. SPB: Siderophore binding proteins in Gram-positive bacteria. VitB12: Vitamin B12. (**B**) Overall surface electrostatic profiles of XusB homologs, calculated using Adaptive Poisson-Boltzmann Solver (ABPS). The color scale represents electrostatic potentials in units of kT/e ranging from −5 (red, negatively charged) to +5 (blue, positively charged).

### Supplementary Fig. 4

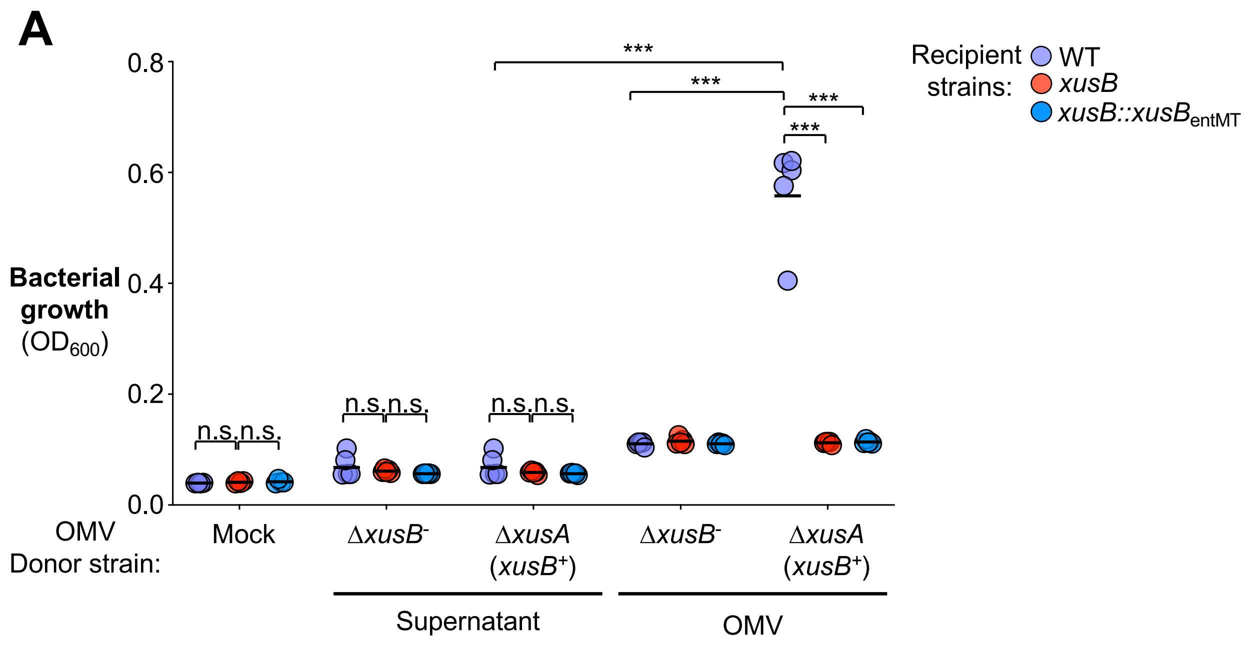

**Supplementary Figure 4: Enterobactin binding is required for XusB-mediated siderophore cross-feeding**

(**A**) Supernatant from donor *B. thetaiotaomicron* strains grown in iron-limited medium (BPS-supplemented SDM medium) was filter-sterilized, loaded with Fe-Ent, ultracentrifuged to collect OMV fraction, washed to remove unbound ligand, and introduced to recipient cells in BPS-supplemented SDM medium. Bacterial growth was measured by $OD_{600}$. Bars represent the geometric mean. ***, $P < 0.001$; n.s., not statistically significant.

#### Supplementary Fig. 5

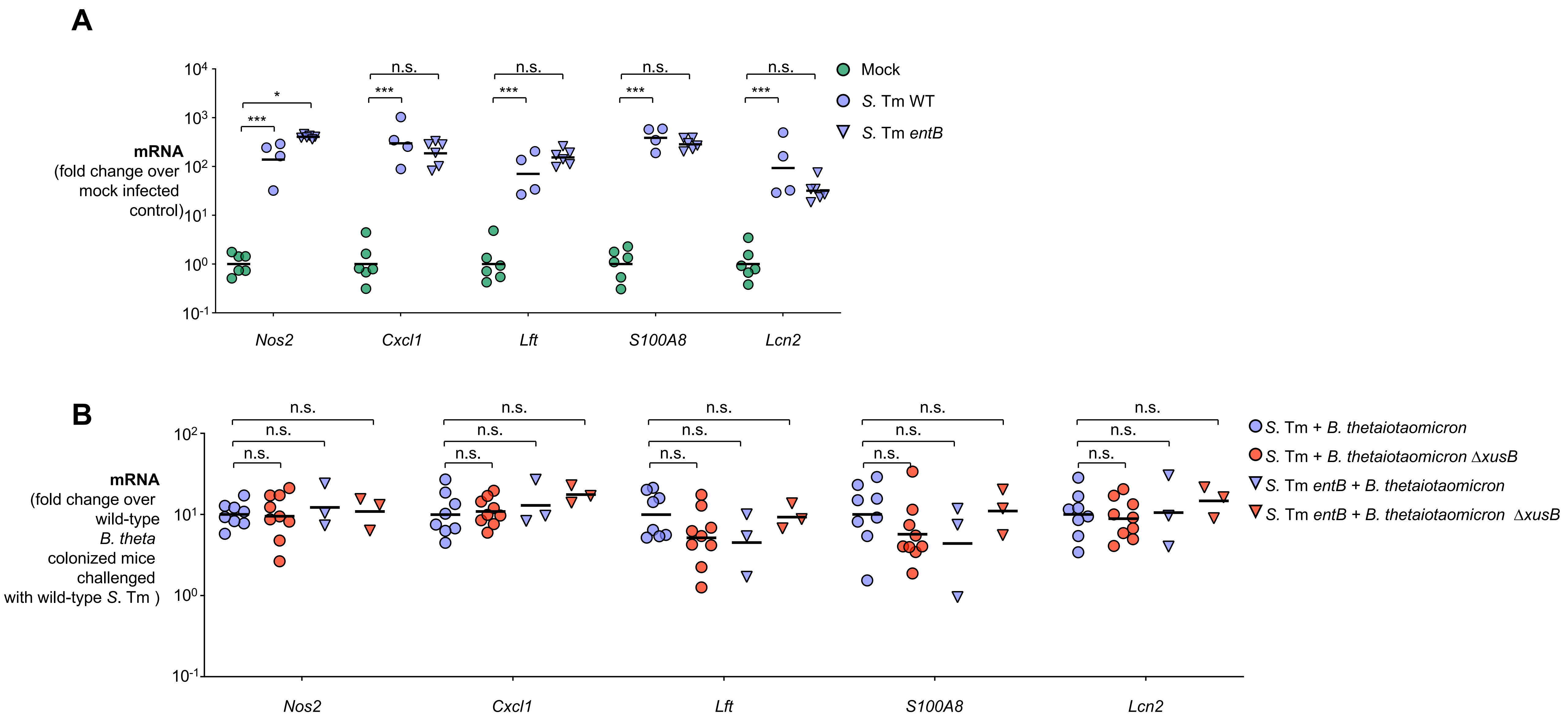

**Supplementary Figure 5: Comparison of inflammatory markers in the infectious colitis model. Related to Fig. 4**

(**A**) Groups of C57BL/6 mice were treated with a cocktail of antibiotics, followed by intragastrical inoculation of an equal mixture of the *B. thetaiotaomicron* wild-type strain and an isogenic mutant (Δ*xusB*), or a strain expressing the enterobactin-binding deficient mutant (Δ*xusB* $xusB_{entMT}$). Mice were then either mock-treated (*N* = 6), intragastrically challenged with *S*. Tm SL1344 (WT vs. Δ*xusB*, *N* = 4; WT vs. Δ*xusB* $xusB_{entMT}$, *N* = 5), or an isogenic *entB* mutant (WT vs. Δ*xusB*, *N* = 6). Four days after infection, the cecal tissue was collected, RNA extracted, and the transcript levels of indicated genes were measured using RT-qPCR (**A**). (**B**) Groups of C57BL/6 mice were treated with a cocktail of antibiotics, followed by intragastrical inoculation of either the *B. thetaiotaomicron* wild-type strain or an isogenic Δ*xusB* mutant. Mice were challenged with the *S*. Tm SL1344 wild-type strain (*B. thetaiotaomicron* wild-type, *N* = 13; *B. thetaiotaomicron* Δ*xusB, N* = 14) or an isogenic *entB* mutant (*B. thetaiotaomicron* wild-type, *N* = 8; *B. thetaiotaomicron* Δ*xusB, N* = 7) for 4 days. The transcript levels of indicated genes in cecal tissue were measured using RT-qPCR (**B**). Bars represent the geometric mean. *, $P < 0.05$; ***; $P < 0.001$; ns, not statistically significant.

### Supplementary Fig. 6

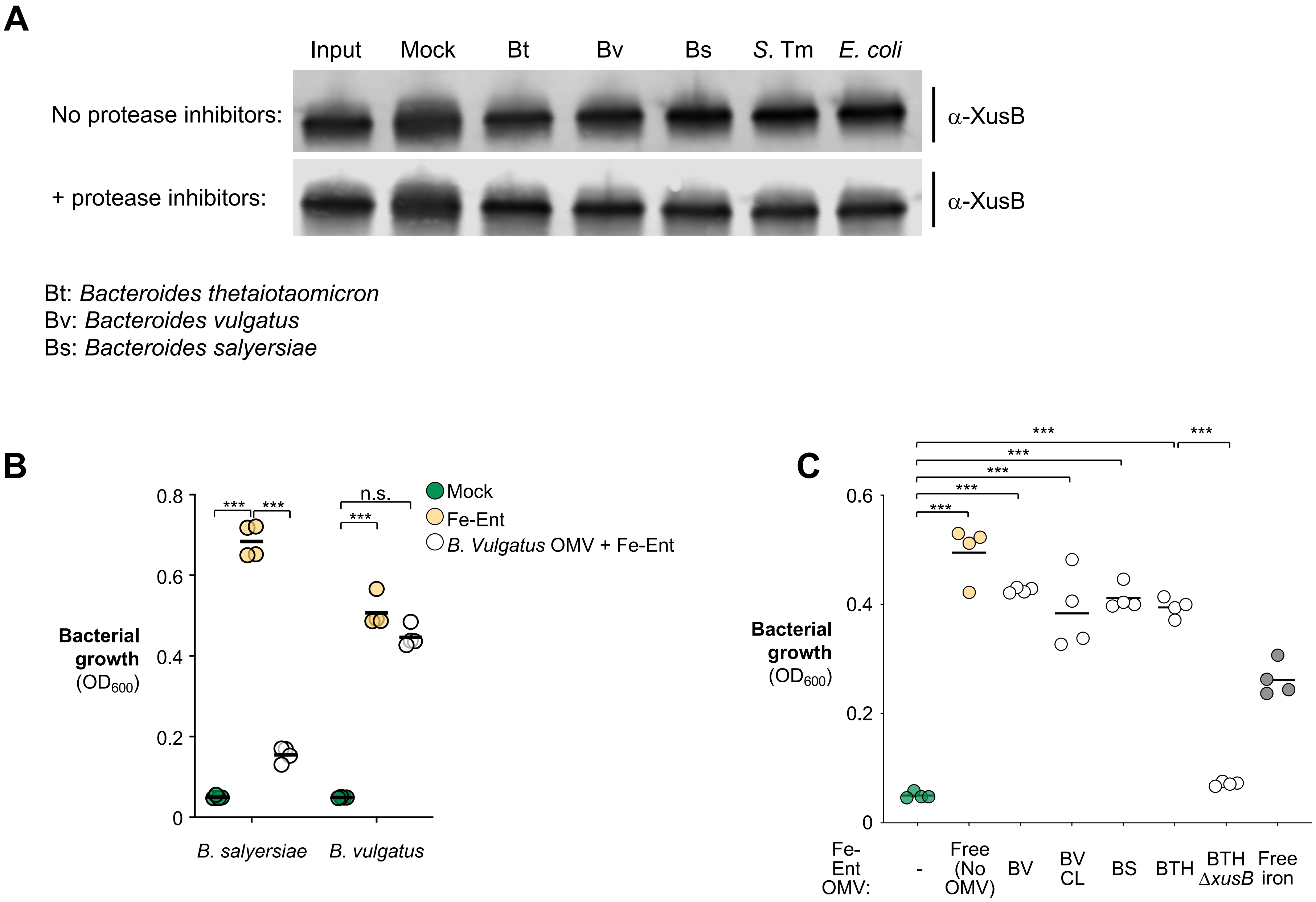

**Supplementary Figure 6: XusB homologs may mediate interbacterial competition for xenosiderophores. Related to Fig. 5**

(**A**) XusB-containing culture supernatant was introduced into $10^8$ CFU of indicated recipient cells in SDM medium in the presence or absence of protease inhibitors. The culture supernatant recovered after 24 hours of anaerobic bacterial growth was resolved on SDS-PAGE gel and probed for XusB. (**B**) Supernatant from donor *B. vulgatus* strains grown in iron-limited medium (BPS-supplemented SDM medium) was filter-sterilized, loaded with Fe-Ent, washed to remove unbound ligand, and introduced to recipient cells in BPS-supplemented SDM medium. Bacterial growth was measured by $OD_{600}$. (**C**) Supernatant from indicated donor strains grown in iron-limited medium (BPS-supplemented SDM medium) was filter-sterilized, loaded with Fe-Ent, washed to remove unbound ligand, and introduced to *S*. Tm *entB* cells in BPS-supplemented SDM medium. Bacterial growth was measured by $OD_{600}$. BV: *Bacteroides vulgatus*; BV CL: *Bacteroides vulgatus* CL09T03C04; BS: *Bacteroides salyersiae* DSM 18765; BTH: *Bacteroides thetaiotaomicron*. Bars represent the geometric mean. ***, $P < 0.001$; n.s., not statistically significant.

### Supplementary Fig. 7

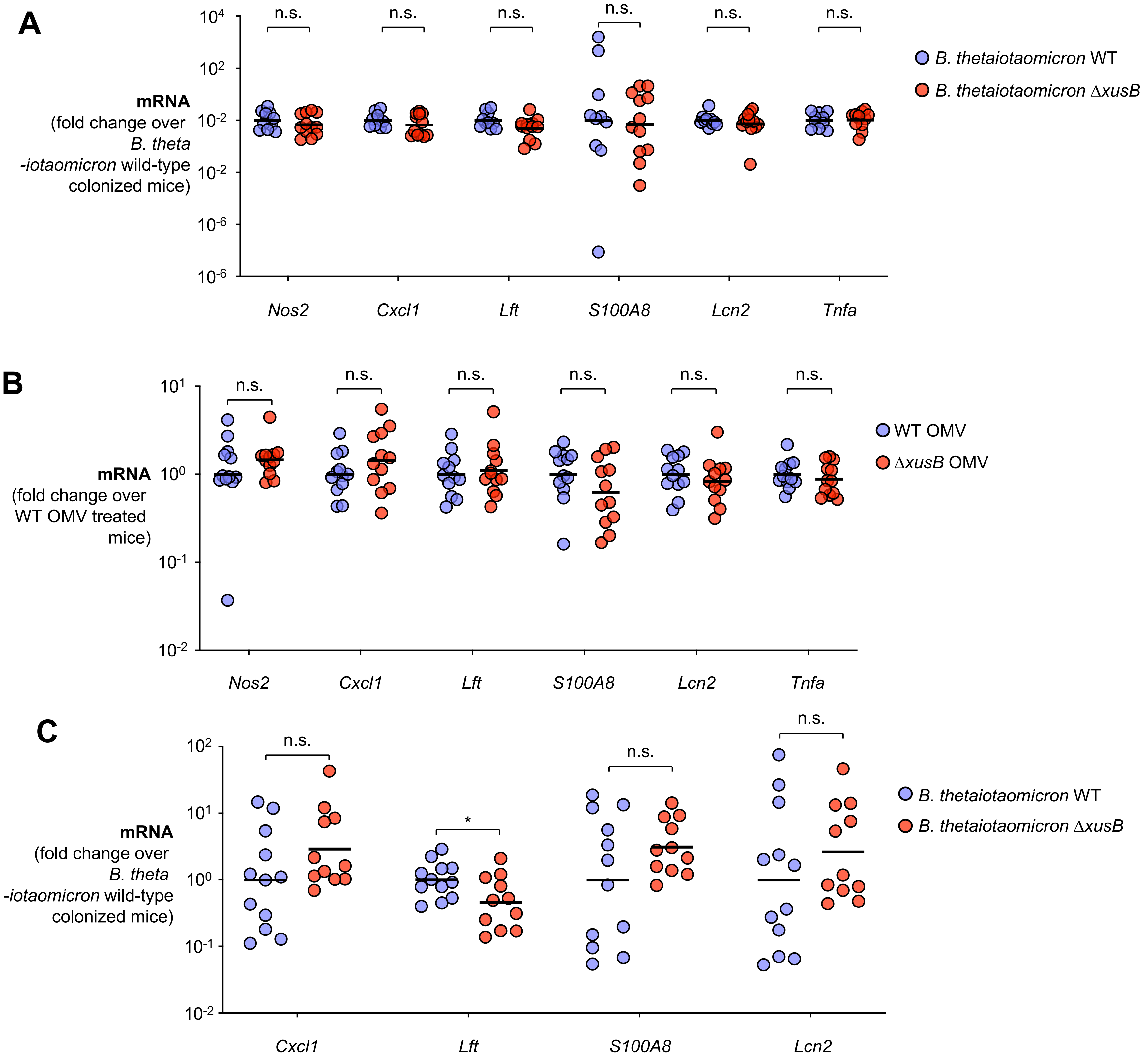

**Supplementary Figure 7: Comparison of inflammatory markers in gnotobiotic and the infectious colitis model. Related to Fig. 7**

(**A**) Groups of C57BL/6 mice were treated with a cocktail of antibiotics, followed by intragastrical inoculation of either the *B. thetaiotaomicron* wild-type strain ($N$ = 10) or an isogenic Δ*xusB* mutant ($N$ = 12). Mice were inoculated with an equal mixture of the *S*. Tm SL1344 wild-type strain and the isogenic Δ*fepA iroN* mutant strain. 4 days later, cecal tissue was collected, RNA extracted, and the transcript levels of indicated genes were measured using RT-qPCR. (**B**) Groups of C57BL/6 mice were treated with a cocktail of antibiotics, followed by intragastrical inoculation of a Δ*xusB* mutant strain. Mice were then inoculated with an equal mixture of the *S*. Tm SL1344 wild-type strain and the isogenic Δ*fepA iroN* mutant strain, followed by intragastrical administration of OMVs derived from the *B. thetaiotaomicron* wild-type strain ($N$ = 11) or the isogenic Δ*xusB* mutant strain ($N$ = 10) daily. mRNA levels of indicated genes in the cecal tissue were determined using RT-qPCR. **(C)** Groups of gnotobiotic Swiss Webster mice were monoassociated with either the *B. thetaiotaomicron* wild-type strain ($N$ = 12) or the isogenic Δ*xusB* mutant ($N$ = 11) for 7 days. Mice were then challenged with an equal mixture of the *S*. Tm SL1344 wild-type strain and the isogenic Δ*fepA iroN* mutant strain. The cecal tissue was collected 3 days post-infection and the mRNA levels of indicated genes were determined using RT-qPCR. Bars represent the geometric mean. *, $P < 0.05$; n.s., not statically significant.

### Supplementary Fig. 8

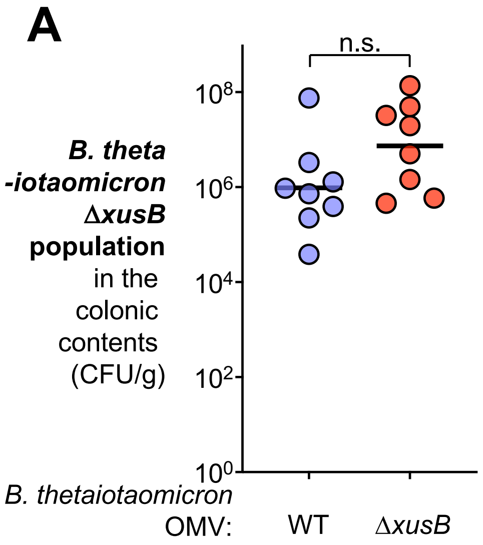


**Supplementary Figure 8: *B. thetaiotaomicron* abundance in an infectious colitis model. Related to Fig. 7**

(**A**) Groups of C57BL/6 mice were treated with a cocktail of antibiotics, followed by intragastrical inoculation of a Δ*xusB* mutant strain. Mice were then inoculated with an equal mixture of the *S*. Tm SL1344 wild-type strain and the isogenic Δ*fepA iroN* mutant strain, followed by intragastrical administration of an equal amount of OMVs derived from the *B. thetaiotaomicron* wild-type strain ($N$ = 11) or the isogenic Δ*xusB* mutant strain ($N$ = 10) daily. The abundance of *B. thetaiotaomicron* in cecal contents was determined 4 days after infection by plating on selective agar. Bars represent the geometric mean. n.s., not statistically significant.
